# Loss of ciliary proteins IFT20 and IFT88 results in defective phagocytosis and metabolism in the RPE

**DOI:** 10.1101/2025.10.16.682843

**Authors:** Peter Andreas Matthiessen, Lotta Elisabeth Wagner, Karsten Boldt, Gunnar Glaßer, Ingo Lieberwirth, Emeline F. Nandrot, Helen Louise May-Simera

## Abstract

A major proportion of retinal disease-causing genes are related to the primary cilium, a microtubule-based signalling organelle essential for multiple developmental pathways. Previous work has shown that the primary cilium plays a crucial role in the development of the retinal pigment epithelium (RPE) affecting homeostasis and function, in particular phagocytosis. We used a cell biology approach to analyse the influence of ciliary genes on RPE phagocytosis and dissect the underlying molecular mechanisms. We found that loss of ciliary trafficking via depletion of *Ift20* and *Ift88* in RPE-J cells resulted in impaired phagocytosis, specifically by reducing photoreceptor outer segments binding, changes in apical membrane morphology and altered mitochondrial metabolism, whereas loss of *Bbs6* showed no functionality phenotype. In addition, proteomics revealed mis-regulated pathways and targets, through which new phagocytosis-related proteins were identified. Our data highlight the role of primary cilia proteins in RPE function and metabolism, essential for visual health.

## Introduction

The primary cilium is a microtubule-based organelle, which extends from the apical membrane of most mammalian cell types and acts as a sensory antenna, receiving and sending signals related to cellular communication^1–3^. It has been shown to play a crucial role in essential developmental processes, regulating multiple important signalling pathways, including Sonic Hedgehog (SHH), Wingless (WNT) and Transforming growth factor-β (TGF-β) signalling^4–7^. Therefore, defects in the structure, function or maintenance of this organelle result in a wide variety of syndromic diseases, termed ciliopathies^8^. These ciliopathies exhibit a diverse collection of overlapping symptoms, which include polydactyly, *situs inversus*, kidney diseases and retinopathies, mostly related to ciliary signalling. The precise mechanisms underlying these dysfunctions are, however, often cell type specific and have not been fully elucidated.

Previous studies have shown that the primary cilium plays a crucial role in the development of the retinal pigment epithelium (RPE)^9–12^. The RPE is a monolayer of highly specialised and tightly connected polarised cells, located between the neural retina and the vascular choroid. Its highly infolded basolateral membrane faces the Bruch’s membrane, which separates the RPE from the choriocapillaris. The RPE further controls the transfer of nutrients and blood supply to the retina^13,14^. The apical membrane of the RPE consists of long actin-based microvilli, so-called apical processes, which ensheathe the light-sensitive photoreceptor outer segments (POS). This close connection enables the exchange of different molecules, for example 11-*cis*-retinal, an essential visual pigment of rod photoreceptors, which is recycled by the RPE and transported back to the photoreceptors^15–19^. Because of this tight interaction, it is not surprising that the RPE is crucial for maintenance and function of the photoreceptors, and together both are seen as one functional unit.

The arguably most important function of the RPE is the phagocytosis of oxidised POS extremities^13,15,20^. POS have to be constantly renewed to ensure photoreceptor function and are detached from the photoreceptor tip in a circadian manner^21^. The process of phagocytosis is highly controlled and regulated by multiple feedback-loops. In short, phagocytosis is initiated by the detection of phosphatidylserines (PtdSer) on the outer leaflet of the plasma membrane^22^ by the main receptors, namely the α_v_β_5_ integrin receptor, Mer tyrosine kinase (MerTK) and scavenger receptors Cluster of Differentiation 36 (CD36) and SR-B2/LIMP-2, either directly (scavenger receptors) or via specific receptor ligands present in the interphotoreceptor matrix such as MFG-E8 (α_v_β_5_ integrin), or Gas6 and Protein S (MerTK)^23–30^. During this process, the α_v_β_5_ integrin receptor contributes to POS binding to the cell and is needed for the circadian activation of the function^25,31,32^. Upon α_v_β_5_ integrin receptor activation, phosphorylation of the focal adhesion kinase (FAK) launches an intracellular signalling cascade and, subsequently, MerTK activates the re-organisation of the actin cytoskeleton initiating downstream pathways for phagosome formation and ultimately POS engulfment^25,33–36^.

This process of phagocytosis generates a high energy demand, therefore, the RPE has a high mitochondria content and mainly generates energy via oxidative metabolism^14,37^. Multiple studies have shown the importance of mitochondrial health in the RPE and that mitochondrial dysfunction can result in multiple RPE degeneration-associated diseases, including AMD or diabetic retinopathy^38,39^.

We have previously shown that the primary cilium not only influences RPE development, but also further aspects of RPE functions. Studies using iPSC-RPE could show that the primary cilium mediates RPE maturation via suppression of the canonical WNT pathway and PKCδ activation^9^. In addition, different ciliary dysfunction mouse models (*Bbs6^-/-^* and *Bbs8^-/-^*), in which ciliary trafficking is impaired, have shown incomplete RPE maturation, polarisation and changes in morphology, resulting in alterations in retinal adhesion. Moreover, there is also limited data for primary cilia dysfunction impacting RPE function. Si-RNA-mediated knockdown of *BBS8* in ARPE-19 cells directly resulted in a decrease in POS phagocytosis^10,12,40^. Furthermore, the *Ift20^null^;Tyrp2-Cre* mouse model, in which the primary cilium is exclusively ablated in the RPE, likewise displayed a defective phagocytosis phenotype, together with reduced retinal adhesion and morphological changes in the RPE^11^. These findings strongly suggest that the primary cilium is essential for RPE functionality and understanding the underlying mechanism is relevant for unravelling ciliary-associated mechanisms of visual function.

Therefore, we adopted a cell biology approach to analyse the influence of ciliary genes on RPE phagocytosis and dissect the underlying molecular mechanisms. For this, we utilised the immortalised RPE-J cell line, which was originally derived from 7-day-old Log-Evans rats. The unique selling point of this RPE cell line is, besides maintaining epithelial cell surface polarity and retaining essential RPE properties such as the expression of the RPE marker RET-PE2, that these cells display efficient phagocytosis activity, making them ideal for studying functional aspects of the RPE^27,28,31,41,42^.

To study the influence of primary cilia dysfunction in these cells, we targeted three ciliary genes that differentially affect ciliary function, namely intraflagellar transport (IFT) protein 20 (*Ift20)*, IFT protein 88 (*Ift88/Polaris)* and Bardet-Biedl syndrome protein 6 (*Bbs6/Mkks)*. IFT20 and IFT88 are both part of the IFT-B complex, which together with the IFT-A complex and BBSome facilitate the trafficking of components in and out of the primary cilium^2,43–46^. Loss of *Ift20* disrupts ciliary trafficking, including trafficking to the cilium and leads to loss of both the axoneme and transition zone^11,47^. Ciliary trafficking is equally impaired upon loss of *Ift88*, but by comparison most IFT88 models still retain the transition zone^5,48–51^. In contrast, BBS6 is part of the BBS chaperonin-like complex, which plays an important role in assembling protein complexes needed for ciliary trafficking, including the BBSome^52^. Compared to the IFT genes, loss of *Bbs6* does change ciliation as a consequence of defective signalling, since loss of *Bbs6* compromises BBSome function^53,54^.

Using these ciliary gene knockdown cells, we analysed the morphology of the cells, their phagocytic and mitochondrial functions, as well as adopted an OMIC approach to investigate alterations in cellular pathways. Our study aims to create new insights into how ciliary proteins affect the vital function of the RPE to phagocytose POS by uncovering the molecular mechanism underlying this phenotype, initiating a better understanding for ciliary-related retinal diseases. We could demonstrate that the loss of IFT proteins changes apical membrane morphology and alters protein expression and mitochondrial metabolism, resulting in defective phagocytosis in the RPE. Furthermore, we identified new players in phagocytosis. Taken together, we could show that primary cilia proteins influence RPE phagocytic function in part via modulation of metabolic processes, highlighting the interconnection between these cellular processes.

## Results

### Dysfunction of IFT results in a defective phagocytosis binding phenotype

We previously observed that RPE models of ciliary dysfunction, namely the *Ift20^null^;Tyrp2-Cre* mouse and ARPE-19 *BBS8* knockdown (KD) cells, displayed photoreceptor outer segment (POS) phagocytosis impairment^10,11^, however, the underlying molecular mechanism have not been investigated so far. To address this question, we generated cilia dysfunction cell models by knocking down the cilia genes *Ift20*, *Ift88* and *Bbs6/Mkks*. Target KD was validated via qPCR and western blot, showing significant reduction of expression for all targets (Figure S1A-B). In addition, we quantified the ciliary phenotype, and observed significant reduction in ciliation in *Ift20* and *Ift88* KD cells, in combination with significant changes in ciliary gene expression in all three KD cells (Figure S1C-E), as expected.

To analyse effects of ciliary gene KD on RPE phagocytosis, we incubated RPE-J cells with FITC-tagged POS for 1.5 h and 4 h, before measuring the amount of bound and internalised POS (Figure 1A). *Ift20* KD cells displayed a decrease in total and bound POS after 1.5 h (Figure 1A, left panel). This effect was even more pronounced after 4 h, showing a significant decrease in comparison to non-targeting control (NTC). Internalisation was unaffected, suggesting a defect specifically in POS binding. This phenotype was similarly observed in *Ift88* KD cells, however the decrease was less pronounced (Figure 1A, middle panel). In contrast, *Bbs6* KD cells displayed no differences in phagocytic function compared to control (Figure 1A, right panel). To further probe changes in the phagocytic pathway, we examined gene and protein expression of key phagocytosis components. *Ift20* KD cells displayed a significant decrease in *Mertk* and *Itgb5* gene expression (Figure 1B, left panel). The MerTK receptor is essential for POS internalisation, whereas ITGB5 together with ITGAV form the α_v_β_5_ integrin dimer receptor essential for POS binding. MerTK expression was similarly downregulated at the protein level. Interestingly, protein expression of ITGB5 was unchanged compared to NTC, however ITGAV expression was significantly decreased (Figure 1C, left panel). In *Ift88* KD cells the phagocytosis components showed a systematic decrease in gene expression (Figure 1B, middle panel). However, this effect was not consistently observed at the protein level, except for ITGAV, which was similarly downregulated (Figure 1C, middle panel). Despite no measurable changes in phagocytic function, *Bbs6* KD cells displayed a significant downregulation of MerTK at the gene and protein levels, and a slight downregulation of *Itgb5* gene expression (Figure 1B and C, right panel). ITGB5 protein expression, however, was unchanged. Interestingly, the data display that mostly the expression of membrane bound receptors was altered upon ciliary dysfunction, whereas the expression of ligands and other cytosolic proteins remained largely stable. These results suggest specific differences in phagocytic function, specifically outer segment binding, and changes in expression of key phagocytosis-related receptors based on the type of ciliary mutation.

**Figure 1.**
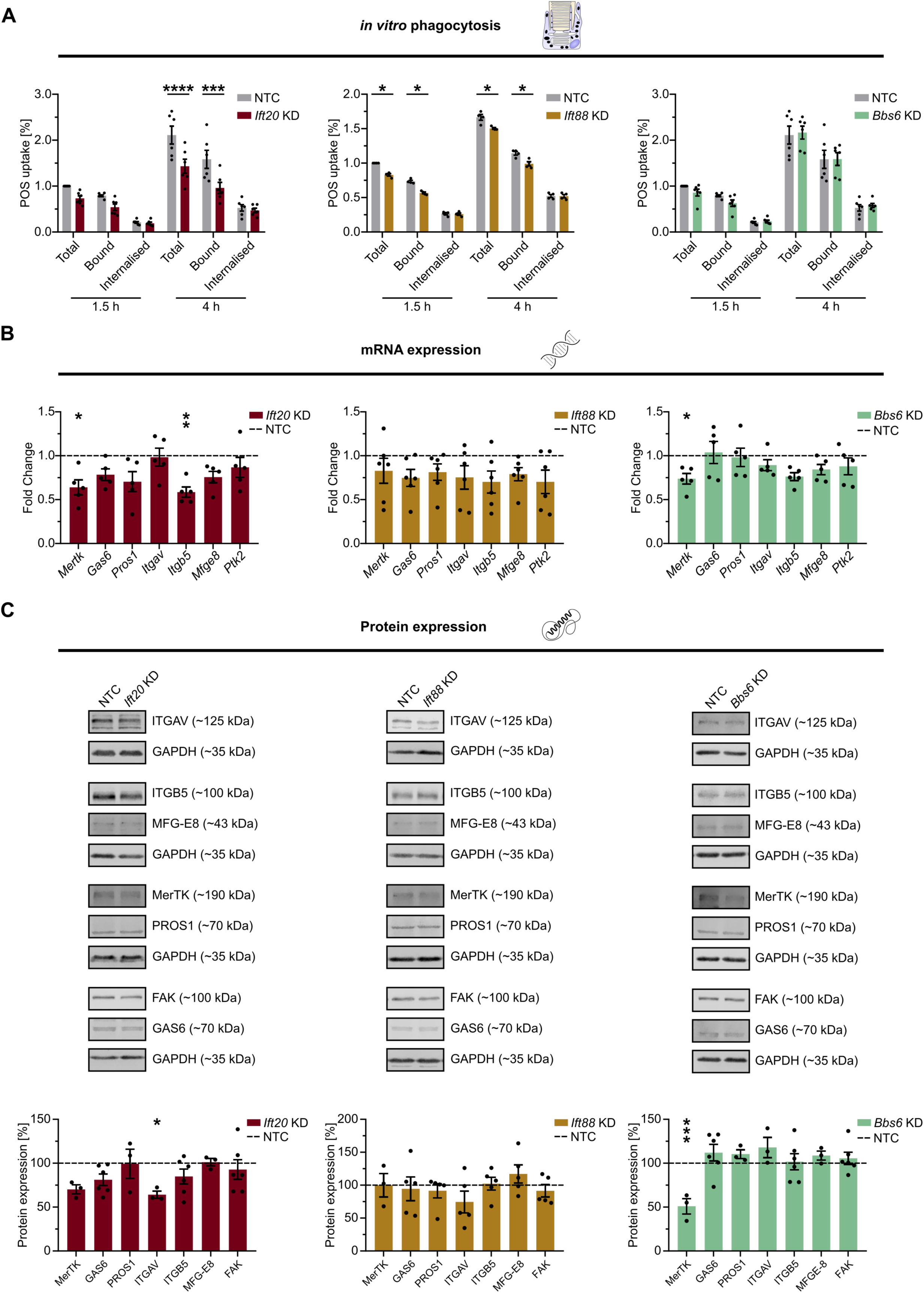
Knockdown of *Ift* genes decreases photoreceptor outer segment binding in RPE-J cells. (A) Phagocytic function of RPE-J KD cells was determined via phagocytosis assay after treatment with FITC-tagged photoreceptor outer segments (POS) for 1.5 and 4 h. Total, bound and internalised POS were measured. Data were normalised to NTC total POS after 1.5 h (n = 4-6). (B) mRNA expression of key phagocytosis pathway players *Mertk*, *Gas6*, *Pros*1, *Itgav*, *Itgb5*, *Mfge8* and *Ptk2* was analysed with RT-qPCR. Data were normalised to NTC (dashed line) and *Rplp0* was used as housekeeping gene (n = 5-6). (C) Protein lysates were immunoblotted to detect and quantify protein expression of key phagocytosis pathway proteins. GAPDH was used as housekeeping protein. Data were normalised to NTC (dashed line) (n = 3-6). Two-way ANOVA test with Sidak’s (A, C) or Tukey’s (B) multiple comparison was used as statistical analysis with the following p-values: * p < 0.05; ** p < 0.01; *** p < 0.001; **** p < 0.0001. NTC was used as control. Data are displayed as mean ± SEM.

### Loss of IFT20 results in aberrant apical microvilli morphology

Since primary cilia deficiencies have been shown to alter RPE maturation and polarisation^9^, we sought to examine overall cell and apical membrane morphology, especially considering that the microvilli-rich apical membrane is essential for phagocytosis of POS *in vivo*^55^. Upon *Ift20*, *Ift88* and *Bbs6* KD, overall cell morphology remained unchanged in the terms of cell area, perimeter, aspect ratio, hexagonality and polygonality (Figure S2A-G). However, a significant decrease in the number of cell neighbours was observed for *Ift88* and *Bbs6* KD cells. We further analysed the polarisation of KD cells via assessment of their transepithelial electrical resistance (TEER) and capacitance (Ccl). Both *Ift20* and *Ift88* KD cells displayed consistently higher TEER values throughout cultivation, suggesting increased tight junction integrity between the cells, whereas *Bbs6* KD TEER values were comparable to NTC (Figure S3A). Ccl can be used as a readout for the degree of apical membrane infolding. Both *Ift20* and *Bbs6* KD cells displayed a decrease in Ccl on cultivation days 5 and 8, which translates to a higher degree of apical membrane infolding (Figure S3B). In contrast, *Ift88* KD showed no difference compared to control. Examination of the apical membrane morphology via scanning electron microscopy revealed that consistent with decreased Ccl values, *Ift20* KD appeared to display a higher number of somewhat longer apical microvilli compared to the NTC control (Figure 2A). In contrast, *Ift88* KD and *Bbs6* KD cells displayed appeared to display shorter microvilli. To further examine apical microvilli, we performed immunofluorescence staining using an antibody against EZRIN, a protein-tyrosine kinase which localises inside microvilli (Figure 2B). Although EZRIN gene and protein expression was not altered upon KD (Figure 2C-D), EZRIN staining looked structurally different in *Ift20* KD cells compared to control, displaying more aggregate-like structures and less homogenous distribution (Figure 2B). In contrast, EZRIN staining of *Ift88* KD and *Bbs6* KD cells looked comparable to the NTC. In addition, visible differences in apical carbohydrate distribution were observed via staining of wheat germ agglutinin (Figure S3C). In specific, a potential increase of apical carbohydrates was seen with *Ift88* KD cells, whereas staining of *Ift20* and *Bbs6* KD cells looked comparable to the NTC. These data indicate changes in apical membrane morphology upon knockdown of *Ift20* and *Ift88*, with the *Ift20* KD cells potentially exhibiting alteration in microvilli structure.

**Figure 2.**
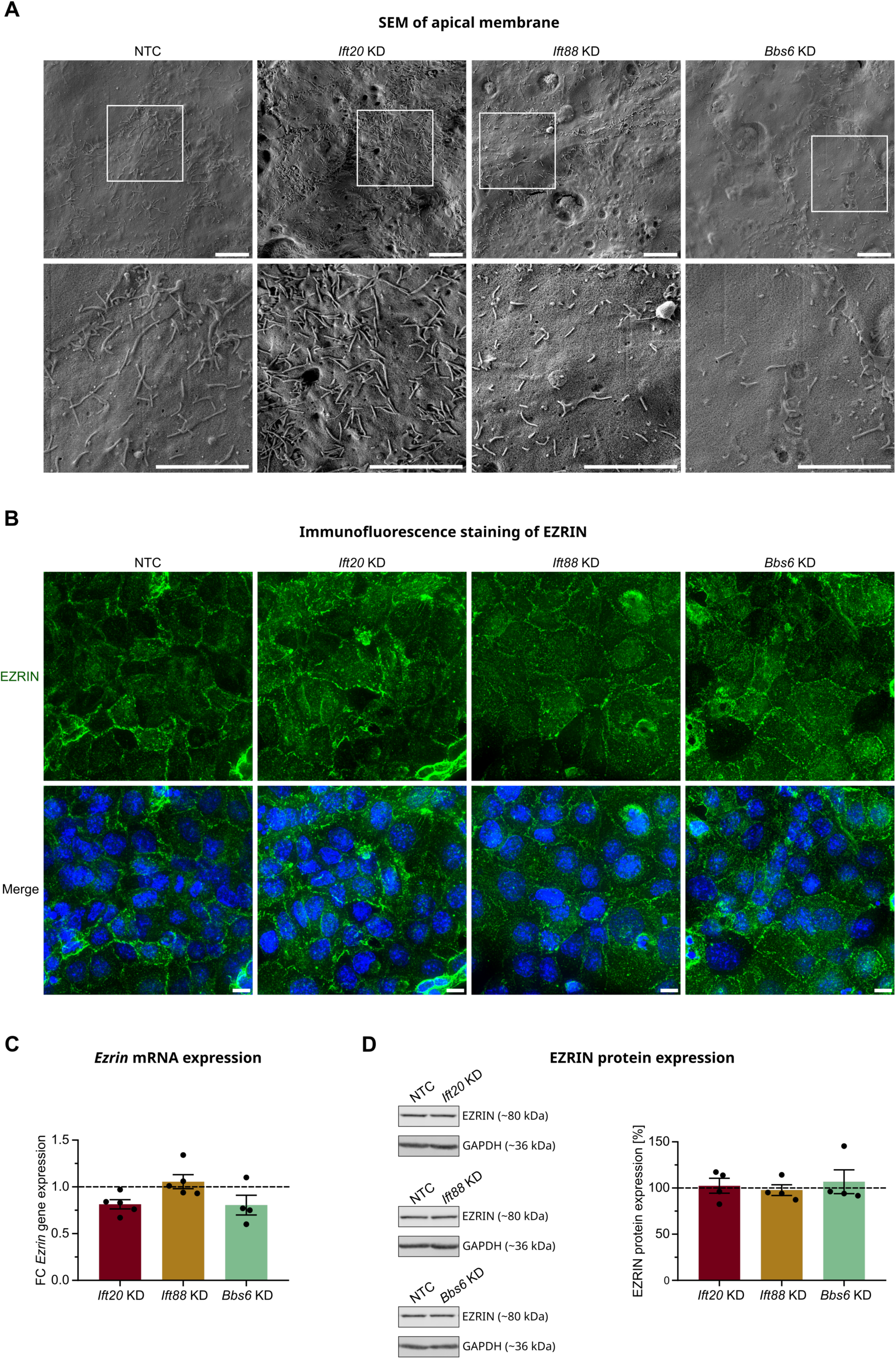
RPE apical membrane morphology seems altered upon ciliary dysfunction. (A) Scanning electron microscopy (SEM) was used to image the apical membrane of RPE-J KD cells and NTC as control. Scale bars represent 4 µm. (B) Apical processes were visualised using EZRIN immunofluorescence staining. EZRIN staining is shown in green and nuclei stained with DAPI in blue. Scale bars represent 10 µm. (C) RT-qPCR was utilized to measure *Ezrin* mRNA expression. Data were normalised to NTC (dashed line, n = 4-5). (D) EZRIN protein levels were determined and quantified via western blot analysis. GAPDH was used as a housekeeping protein and data were normalised to NTC (dashed line, n = 4). Two-way ANOVA test with Sidak’s (C) or Tukey’s (D) multiple comparison was used as statistical analysis with the following p-values: * p < 0.05; ** p < 0.01; *** p < 0.001; **** p < 0.0001. Data is shown as mean ± SEM.

### Ciliary dysfunction effects phagocytosis and cytoskeleton pathways

To identify further molecular processes that might be affected upon ciliary dysfunction in the RPE we performed mass spectrometry. Significantly mis-regulated proteins in the KD cells compared to the NTC were categorised into two tiers based on stringency criteria (see Methods). All three KD cells exhibited multiple mis-regulated proteins compared to the NTC control (Figure 3A-B). More specifically, *Ift20* KD resulted in 20 up- and 116 down-regulated Tier 2 proteins, whereas in *Ift88* KD cells 103 proteins were up- and 163 down-regulated. *Bbs6* KD resulted in 66 up- and 82 down-regulated proteins, clearly suggesting that ciliary dysfunction influences RPE gene expression. We further filtered these hits for functionality-related proteins, including phagocytosis, phago-/lysosomes, melanogenesis and visual cycle proteins (Table S3). Several mis-regulated targets were found upon all three KDs (Figure 3C). Most of these targets were related to phagocytosis or phago- and lysosome (Figure 3D), displaying a specific effect of ciliary dysfunction on the phagocytosis pathway. Ciliary dysfunction has already been shown to have a major impact on the cytoskeleton^56,57^, and actin cytoskeleton remodelling is important in the internalisation process of POS via formation of the phagocytic cup and then of the phagosome^58–60^. Therefore, it was encouraging to find mis-regulated expression of multiple cytoskeleton-related proteins upon all three KDs (Figure 3E; Table S3; Tier 2 *Ift20*: 11 up- and 24 down-regulated; *Ift88*: 27 up- and 34 down-regulated; *Bbs6*: 9 up- and 24 down-regulated), demonstrating a key impact of ciliary dysfunction on the cytoskeleton.

**Figure 3.**
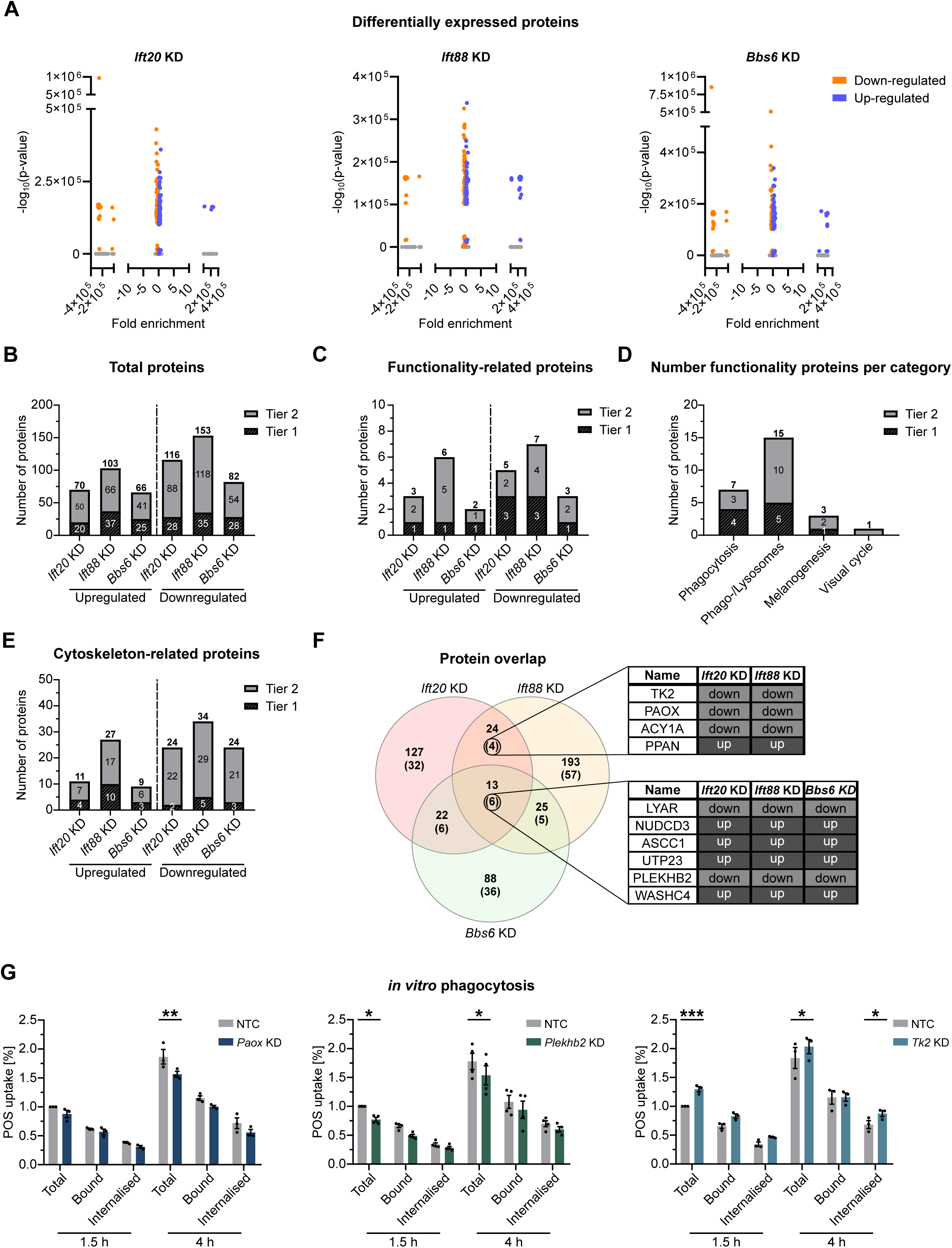
**Mass spectrometry of RPE-J KD cells displays potentially altered pathways and targets.** (A) Volcano plots of mass spectrometry values of RPE-J KD cells compared to the NTC. Upregulated hits are displayed in blue, downregulated hits in orange (n = 4). Tier 1: hits with Student’s t-test = log p > 1.3 and Significance A = p ≤ 0.05; Tier 2: hits with Student’s t-test = log p > 1.3. Data were displayed as diagrams showing total numbers of mis-regulated proteins (B), numbers of mis-regulated functionality-related proteins (C), including phagocytosis, phago-/lysosome, melanogenesis and visual cycle-related proteins, numbers of functionality-related proteins sorted by category (D) and numbers of mis-regulated cytoskeleton-related proteins (E). (F) VENN diagram displaying overlap of mis-regulated proteins between RPE-J KD cells. Number of Tier 1 hits displayed with, Tier 2 hits without brackets. Commonly mis-regulated Tier 1 proteins are listed in the tables on the right, down-regulated genes highlighted in grey and up-regulated genes highlighted in black. (G) Phagocytic function of RPE-J *Paox*, *Plekhb2* and *Tk2* KD cells was determined after incubation with FITC-tagged photoreceptor outer segments (POS) for 1.5 and 4 h. Total, bound and internalised POS were measured. NTC was used as control. Data were normalised to NTC total POS after 1.5 h (n = 3-4). Two-way ANOVA test with Sidak’s multiple comparisons was used as statistical analysis with the following p-values: * p < 0.05; ** p < 0.01; *** p < 0.001; **** p < 0.0001. Data are displayed as mean ± SEM.

### Proteomic approach reveals novel genes involved in RPE phagocytosis

We next analysed the overlap of mis-regulated proteins between our 3 genes (Figure 3F). 6 Tier 1 proteins were commonly affected in all 3 KD cells (ASCC1, LYAR, NUDCD3, PLEKHB2, WASHC4, UTP23). A further 4 Tier 1 proteins overlapped between *Ift20* and *Ift88* KD cells (ACY1A, PAOX, PPAN, TK2). Interestingly, PAOX is involved in polyamine metabolism^61,62^, whereas TK2 plays a role in the phosphorylation of the nucleotides thymidine, deoxycytidine, and deoxyuridine in the mitochondrial matrix for mitochondrial DNA synthesis and maintenance, and mitochondrial gene expression^63,64^. Both were commonly downregulated in both *Ift* KD cell models, which showed defective phagocytosis. PLEKHB2 (also known as EVT-2), down-regulated in all 3 KD cells, binds phosphatidylserines, similarly to phagocytosis receptor and/or ligands, and regulates the retrograde transport of recycling endosomes to the Golgi apparatus^65^. Although not previously reported to be directly associated with phagocytosis, we examined their potential to influence this. Upon *Paox* KD, cells displayed a significant decrease in total POS, as well as a slight decrease in bound and internalised POS after 4 h (Figure 3G, left panel; knockdown efficiency: Figure S4). *Plekhb2* KD cells also revealed a significant decrease in total POS, in addition to a slight decrease in bound POS at both time points, thus displaying a similar phenotype to the *Ift* KD cells (Figure 3G, middle panel). Conversely, *Tk2* KD cells showed a significant higher amount of total POS after 1.5 and 4 h and internalised POS after 4 h, together with a slight increase in bound and internalised POS after 1.5 h (Figure 3G, right panel). These results strengthen the hypothesis that dysfunction of ciliary proteins disrupts phagocytosis, and identifies potential new players involved.

### Ciliary dysfunction results in mis-regulation of membrane- and metabolism-related pathways

To create a detailed overview of which cellular pathways are impacted upon ciliary dysfunction we performed GO-Term and pathway analysis. The overlap of GO-Terms displayed 10 common terms in *Ift20*, *Ift88* and *Bbs6* KD cells (Figure 4A-B). These included terms important for RPE function and morphology such as ‘focal adhesion’, ‘cadherin binding’ and ‘membrane’, as well as the term ‘extracellular exosome’ (Figure 4B). We have previously described the role of primary cilia dysfunction on the release and composition of extracellular vesicles^66^. Two further terms overlapped between *Ift20* and *Ift88* KD cells (Figure 4A, C), namely ‘endoplasmic reticulum membrane’ and ‘ATP binding’, which could indirectly impact RPE function via metabolism or protein synthesis. We also checked for overlapping pathways, as defined by KEGG and REACT, and found 6 that are commonly affected for all 3 genes KD (Figure 4D-E). These included trafficking and metabolism pathways, both processes already known to be important for RPE functionality, as well as Wnt signalling, well known to be associated with the primary cilium and also related to RPE differentiation^5,67^. We found 24 pathways overlapping between *Ift20* and *Ift88* KD cells. The top 10 hits of these included the phagocytosis-related pathway ‘ER-phagosome pathway’ and pathways involving CyclinD, a target of Wnt signalling (Figure 4D, F).

**Figure 4.**
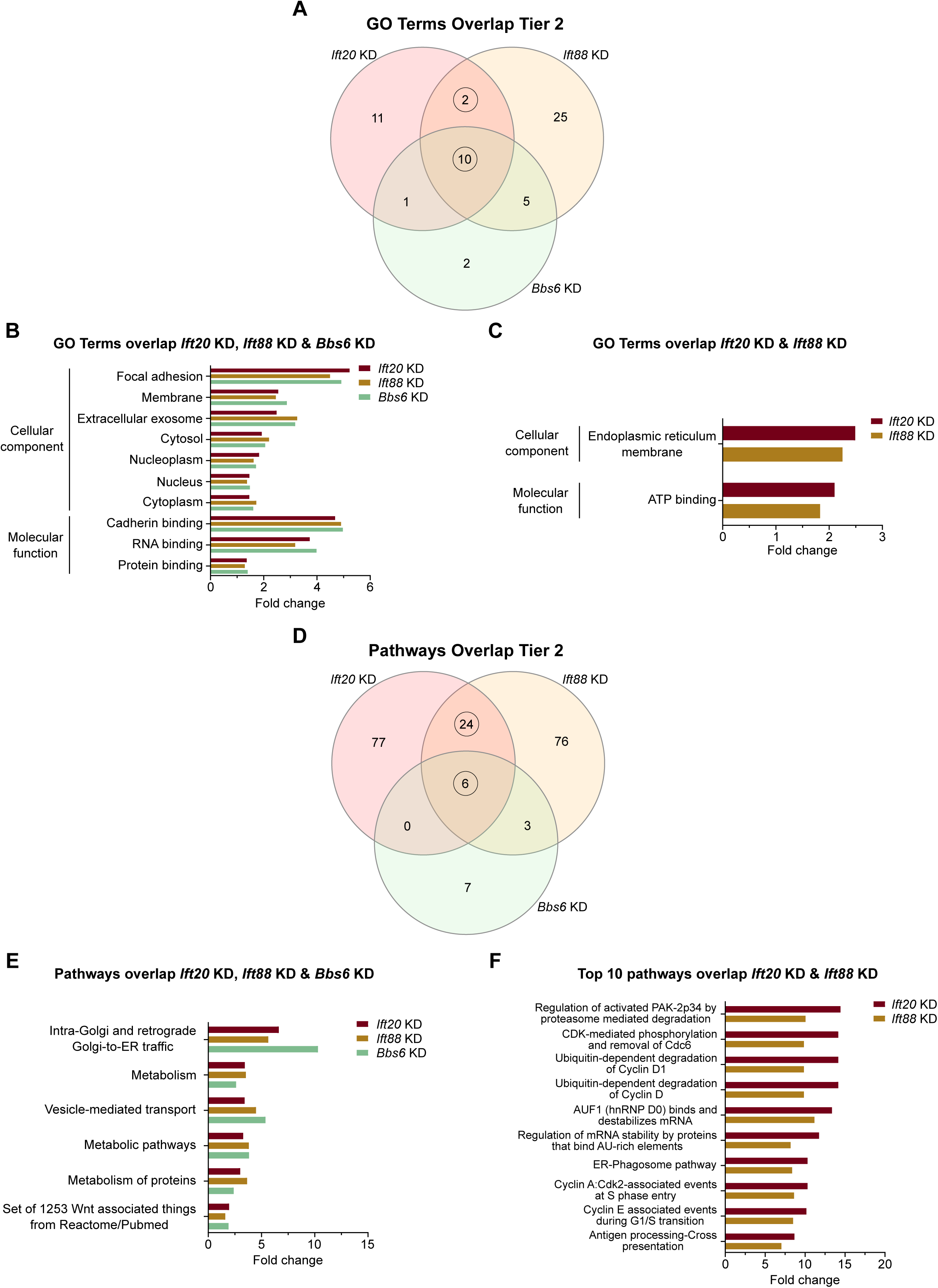
**Gene ontology and pathway analysis reveal altered cellular pathway upon ciliary gene knockdown.** (A) VENN diagram displaying overlap of mis-regulated GO terms between RPE-J KD cells. Bar charts display commonly mis-regulated GO terms in *Ift20*, *Ift88* and *Bbs6* KD cells (B) and GO terms commonly mis-regulated in only *Ift20* and *Ift88* KD cells (C). (D) VENN diagram displaying overlap of mis-regulated pathway terms between RPE-J KD cells. Bar charts display commonly mis-regulated GO terms in *Ift20*, *Ift88* and *Bbs6* KD cells (E) and GO terms commonly mis-regulated in only *Ift20* and *Ift88* KD cells (F). Analysis was performed using Tier 2 hits and GetGo (n = 4).

Upon analysis of GO-terms and pathways from the individual KD cells we identified processes uniquely mis-regulated in single cell lines. For *Ift20* KD cells 24 GO-terms and 107 pathways were identified, whereas *Ift88* KD cells displayed 42 GO-terms and 109 pathways. For *Bbs6* KD cells 18 GO-terms and 17 pathways were identified. The Top 20 hits for *Ift20* and *Ift88* KD were examined for the analysis, whereas for *Bbs6* KD all hits were observed. In addition to GO-terms and pathways related to transport and the proteasome, the term ‘ATP hydrolysis activity’ was detected for *Ift20* KD cells, again linking the KD to effects on protein synthesis and metabolism (Figure S5A, D). In contrast, *Ift88* KD cells had mis-regulated proteins associated with GO-terms related to the actin cytoskeleton, endocytic and phagocytic vesicles, and terms related to phagocytosis. In addition, proteins for mitophagy and glycogen synthesis pathways were affected, again suggesting a potential effect on metabolism (Figure S5B, E). Proteomic profiling of *Bbs6* KD cells identified changes in proteins involved in cytoskeleton- and membrane-related processes, namely ‘actin filament binding’, ‘integral component of membrane’ and pathway ‘membrane trafficking’ (Figure S5C), exhibiting potential effects on membrane composition and function. Further pathways related to metabolism were also observed, including ‘ATP hydrolysis activity’, ‘The citric acid cycle and respiratory electron transport’ (Figure S5C, F). In summary, this proteomic data further strengthens the findings that phagocytosis-related processes are mis-regulated in the KD cell lines and suggests cytoskeletal, membrane- and metabolism-related processes might be affected upon ciliary dysfunction and contribute to the phenotype. As some terms and pathways however do not overlap between all three KD cells and are only affected individually suggests involvement of not only ciliary-related, but also non-ciliary-related functions of the three targets in this process.

### Loss of IFT proteins leads to mitochondrial metabolism defect

As our OMIC analysis provided us with multiple links to mitochondrial metabolism, we set out to examine this in more details. Filtering the mass spectrometry data for MitoCarta 3.0^68^ proteins revealed several mis-regulated hits for *Ift20*, *Ift88* and *Bbs6* KD cells, strengthening this suspicion (Figure 5A; Tier 2 *Ift20*: 13 up- and 11 down-regulated; *Ift88*: 9 up- and 16 down-regulated; *Bbs6*: 7 up- and 4 down-regulated). Although no difference in expression of the essential mitochondrial genes *Cox4i*, *Atp6* and *Nd4* was observed, apparent differences in OxPhos complex expression were detected via western blot (Figure 5B-C). *Ift88* KD cells displayed a systematic decrease in the expression of all complexes, including a significant decrease in complex V (ATP synthase). In *Ift20* KD cells, complex III (Cytochrome BC1 complex) expression, was significantly reduced. In addition, complexes IV (Cytochrome C oxidase) and V also showed a similar pattern in down-regulation. In contrast, in *Bbs6* KD cells an upregulation of complex II (Fumarate reductase) was detected. We then analysed the mitochondrial function in detail and could observe changes in the oxygen consumption rates (OCR) (Figure S6A). *Ift20* KD cells displayed significant up-regulation of the basal respiration, in addition to a down-regulation of the spare respiratory capacity compared to NTC control cells (Figure 5C). *Ift88* KD cells similarly exhibited a significant up-regulation of basal respiration, together with significant higher proton leak and ATP production, and an increase in spare respiratory capacity compared to control cells (Figure 5D). Additionally, significant differences in the extracellular acidification rate (ECAR) were observed, especially after adding oligomycin, an ATP synthase inhibitor, and carbonyl cyanide-4 (trifluoromethoxy) phenylhydrazone (FCCP), an uncoupling agent that collapses the proton gradient and, thus, disrupts mitochondrial membrane potential (Figure S6B). This acidification of the extracellular milieu points towards *Ift88* KD cells potentially having a shift in metabolism. Looking at the energy map, a shift from aerobic to energetic metabolism could indeed be observed for *Ift88* KD cells in comparison to the NTC (Figure S6C). This change in metabolism was not visible for *Ift20* or *Bbs6* KD cells. In contrast, *Bbs6* KD cells displayed a trend of slight decrease in maximal respiration and ATP production compared to NTC, but no real significant difference (Figure 5D). We further calculated the bioenergetic health index (BHI) as another readout for mitochondrial fitness. *Ift20*, *Ift88* and *Bbs6* KD cells all showed a decreased BHI in comparison to the NTC, displaying mitochondrial dysfunction and reduced mitochondrial health (Figure S6D). Taken together, these data show that loss of ciliary genes, especially loss of *Ift*, affects mitochondrial function in the RPE and result in significant mis-regulation of mitochondrial metabolism.

**Figure 5.**
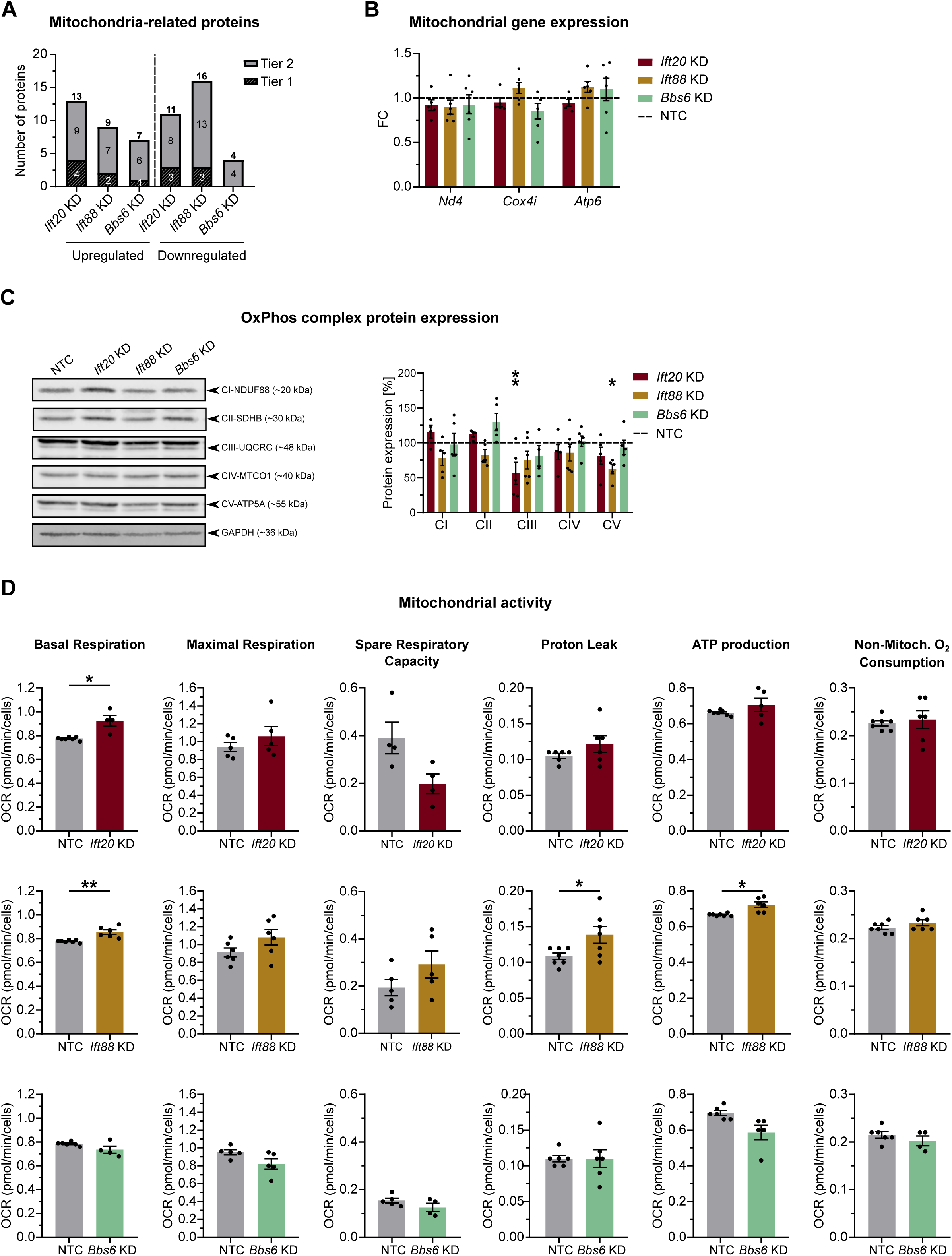
**Ciliary dysfunction affects in mitochondrial metabolism in RPE cells.** (A) Mass spectrometry data was filtered for mitochondria-related proteins using MitoCarta 3.0 and number of mis-regulated hits was displayed. Tier 1: hits with Student’s t-test = log p > 1.3 and Significance A = p ≤ 0.05); Tier 2: hits with Student’s t-test = log p > 1.3 (n = 4). (B) mRNA expression of mitochondrial genes *Nd4, Cox4i* and *Atp6* in RPE-J KD cells was analysed via RT-qPCR (n = 5-6). *Rplp0* was used as housekeeping gene and data were normalised to NTC (dashed line). Two-way ANOVA test with Sidak’s multiple comparison was used as statistical analysis with the following p-values: * p < 0.05; ** p < 0.01; *** p < 0.001; **** p < 0.0001. (C) OxPhos protein complex expression in RPE-J KD cells was determined and quantified using western blot analysis. GAPDH was used as housekeeping protein and data were normalised to NTC (dashed line, n = 4-6) (D) Evaluation of the mitochondrial function. NTC was used as control (n = 4-7). Two-way ANOVA test with Sidak’s (B) or Tukey’s (C) multiple comparison, or unpaired t-test with Welch’s correction (D) were used as statistical analysis with the following p-values: * p < 0.05; ** p < 0.01; *** p < 0.001; **** p < 0.0001. Data are displayed as mean ± SEM.

## Discussion

Primary cilia dysfunction disrupts multiple cellular signalling pathways, resulting in a plethora of diseases termed ciliopathies^4–8^. These display a variety of phenotypes, with retinal degeneration being one of the most prominent. We previously showed that the primary cilium does not only play a crucial role in the development and maturation of the RPE, but also influences homeostasis and functionality^9–12^. The mechanism of this however is not clear, nor how individual cilia proteins specifically influence particular aspects of RPE functionality. Therefore, as it constitutes one of its major functions dependent on microvilli structure, we analysed the influence of primary cilia on the phagocytic function of RPE cells. To dissect this mechanism in depth, we chose diverse forms of ciliary dysfunction by ablating different ciliary proteins, namely IFT20, IFT88 and BBS6. To recapitulate our findings, loss of *Ift20* or *Ift88* disrupts ciliary trafficking and results in structural changes of the primary cilium, whereas loss of *Bbs6* changes ciliation as a consequence of defective signalling^5,11,47–51,53,54^.

Here we demonstrated that loss of IFT proteins leads to defective phagocytosis, more specifically impaired binding of POS to the cell surface. We could show that this was attributable to differences in microvilli structure. Mass spectrometry of all three KD cells not only revealed mis-regulation of multiple functionality-related proteins, but also proteins involved in cytoskeletal processes, trafficking and metabolism. Analysing the overlapping mis-regulated proteins we found new players in phagocytosis. Based on mis-regulated proteins found in our proteomic screen, we further analysed metabolic changes and showed significant mis-regulation of mitochondrial function and OxPhos protein expression upon loss of IFT. We conclude that primary cilia have a direct influence on RPE phagocytosis, yet the severity of this is dependent on the type of ciliary dysfunction.

KD of *Ift20* and *Ift88* resulted in a POS binding defect in RPE cells. Loss of either disrupts ciliary trafficking and leads to structural changes of the cilia. This effect on phagocytosis could be due to incorrect trafficking of receptors to the apical membrane since we detected reduced expression of indispensable phagocytosis pathway receptors. These include ITGB5 and ITGAV, known to be essential for POS binding, and MerTK, together with multiple GO-Terms and pathways associated with trafficking. IFT proteins are known to facilitate the transport of multiple proteins from the cytoplasm and Golgi apparatus to the base of the cilium base^44,45,69,70^. IFT20 has also been shown to be involved in cilia-independent receptor trafficking in T cells^70^, which led to the accumulation of T cell antigen receptor in RAB5-positive vesicles, suggesting IFT20 plays a role in trafficking receptors from early to recycling endosomes. Therefore, it is feasible that IFT proteins might also facilitate the transport of phagocytosis receptors to the apical membrane of RPE cells. Another possible explanation for the observed phagocytosis phenotype is the alteration in apical microvilli structure, which could also underlie the reduced expression of receptors. The microvilli connect the RPE and POS and are essential for phagocytosis^13^. We previously showed that the *Ift20^null^;Tyrp2-Cre* mouse model displayed morphological changes in the RPE and reduced retinal adhesion^11^. Here we observed changes in microvilli morphology upon ciliary dysfunction in SEM and immunocytochemical staining, particularly pronounced upon KD of *Ift20*, which seemed to have a distinctive effect on overall microvilli structure. In contrast, KD of *Bbs6* did not result in alteration in phagocytosis function. Therefore, it seems as if loss of BBS6, which primarily effects ciliary signalling, does not have a direct effect on RPE functionality. As *Bbs6^-/-^* mice exhibit a slow rate of retinal degeneration, it is likely that BBS6 indirectly influences RPE and retinal health^40^.

While observing the overlap of mis-regulated proteins between the different KD cells we identified three proteins of interest: PAOX and TK2, down-regulated in both *Ift* KD cell models correlating with defective phagocytosis, and PLEKHB2, down-regulated in all three KD cells. To our knowledge, none of these proteins had previously been linked to phagocytosis, yet we demonstrated that KD of either of these targets influenced RPE phagocytosis, displaying novel roles of these proteins in RPE functionality.

Both PAOX and PLEKHB2 mimicked the phenotype of the *Ift* KD cells, however the effect on POS binding was not as severe. This could show an indirect effect of these proteins on the phagocytic pathway, which might be partially rescued by other factors. Interestingly, these proteins are involved in cellular pathways, which can be indirectly linked to RPE homeostasis. PAOX is involved in polyamine metabolism^61,62^. Polyamines are reported to involved in a broad spectrum of important biochemical and cellular signalling pathways, such as synthesis and maintenance of nucleic acids and proteins, DNA binding and transcription, RNA splicing and cytoskeletal function^71–74^. Most of these functions are important for normal cell homeostasis and as such mis-regulation could have an impact on RPE functionality, in particular the alteration of the cytoskeleton. This is due to the fact that each RPE cell lies in contact with multiple POS over its numerous apical actin-based microvilli, therefore, dysregulation in cytoskeletal processes most probably would directly impact POS phagocytosis.

PLEKHB2, also known as EVT-2, is able to bind phosphatidylserines due to its N-terminal pleckstrin homology domain, as do phagocytosis receptor ligands. Additionally, depletion of PLEKHB2 in COS-1 fibroblast cells supressed membrane trafficking of recycling endosomes to the Golgi apparatus, showing a role in retrograde traffic regulation of recycling endosomes^65^. POS are internalised, packed into phagosomes and trafficked towards the basal membrane of the RPE, where they fuse with lysosomes and the degradation process takes place. During the maturation process, the phagosome interacts with early and late endosomes to acquire degradative enzymes, and gradually acidifies^14,75^. Downregulation of PLEKHB2 might therefore cause trafficking issues of endosomes containing these important proteins needed for the maturation of phagosomes, offsetting the equilibrium of the phagocytosis pathway and resulting in impairment of RPE functionality.

Intriguingly, KD of *Tk2* led to an increased rate of phagocytosis. TK2 is essential for mitochondrial DNA synthesis and maintenance, and mitochondrial gene expression^63,64^. The RPE has a high energy demand and therefore a high mitochondria content^37^. In addition, during phagocytosis of POS, energy metabolism, high oxygen partial pressure and light injury all result in the production of reactive oxygen species. If these are not efficiently removed, accumulation can lead to oxidative stress damage and result in mitochondria dysfunction^76–78^. Multiple RPE degeneration-associated diseases have been linked to mitochondria dysfunction, for example age-related macular degeneration or diabetic retinopathy^38,39^. KD of *Tk2* most likely results in mitochondrial stress due to its role in mitochondrial gene expression. This mis- regulation could therefore result in a shift in metabolism. Besides the mitochondria, RPE cells use fatty acids, derived from POS, to generate energy through beta-oxidation and subsequent ketogenesis^14,79^. Hence, upregulation of phagocytosis could be an attempt to get access to more lipids and rescue possible energy deficiencies. Since *Tk2* was downregulated upon ciliary dysfunction, and we also have evidence that the mitochondrial metabolism is directly altered in the ciliary gene KD cells, this data presenting a potential link of the primary cilia not only to RPE functionality, but also directly to RPE metabolism that sustain its activities.

As already mentioned, mitochondrial function and health have been shown to be indispensable for RPE homeostasis and functionality^76–78^. As we found multiple mitochondria-related proteins mis-regulated in our OMIC screen, we took a more precise look at the effect of ciliary dysfunction on mitochondria. The differences in key OxPhos complex expression already gave us a hint, that mitochondrial metabolism might be affected by ciliary dysfunction. Analysing mitochondrial function in depth we could show definite alterations upon loss of either of the IFT proteins. Defective metabolism in the RPE most probably contributes to the defective functionality phenotype, as the cells have not got enough energy to perform the energy-costly process of phagocytosis. The significant increase in ECAR in *Ift88* KD cells after uncoupling the proton gradient indicates a potential switch in metabolism, which was also seen in the energy map with a shift towards energetic metabolism. This coincides with an increase in mitochondrial respiration and glycolysis. This upregulation, especially of glycolysis, might be an attempt to rescue the changes in metabolism due to impaired oxidative phosphorylation. The RPE usually does not use glycolysis as its main energy supply, as it is less efficient in ATP production. In addition, glucose is normally shuttled from the RPE to photoreceptor cells, which use glucose as their main source of energy^14^. The RPE switching its metabolism to also use glycolysis to obtain sufficient energy would, therefore, have a big impact on photoreceptor metabolism, resulting in energy starvation and eventually leading to photoreceptor degeneration. This should however be analysed and confirmed in future studies. Loss of *Bbs6* did not alter metabolism, again hinting towards the hypothesis that ciliary signalling dysfunction on its own only indirectly impacts RPE health.

Our approach involved analysing diverse forms of ciliary dysfunction by investigating different ciliary targets. Both *Ift* KD cells showed the same effect on phagocytosis, whereas *Bbs6* KD cells displayed no phenotype. This demonstrates a greater effect of ciliary structure alterations on RPE functionality, while ciliary signalling dysfunction alone does not seem to have a direct effect. This should be confirmed by using different targets involved in ciliary signalling, for example BBS1, BBS8 or BBS10^80–83^.

The alterations on a molecular level differed between all three targets, as seen in proteomics and mitochondrial function. These differences could point to partial cilia-independent influence of the analysed targets, for which several alternative functions have been described. IFT20 has been shown to play a role in autophagosome formation, planar cell polarity and lysosomal degradation^84–86^, and also to be involved in receptor & signalling molecule trafficking in unciliated T cells^70,87^. IFT88 has been described to be involved in spindle orientation, planar cell polarity establishment, and actin organisation^88–90^. Similarly, BBS6 is also involved in cytoskeleton organisation by regulation of actin via Fascin-1^56,91^. In addition, BBS6 also acts in cell division via centrosomal association^92^ and localises to the nucleus, potentially regulating gene expression^93,94^. These cilia-independent functions are further explanations for how RPE functionality could be affected, or at least contribute to the phenotype. In particular, autophagosome formation, lysosomal degradation and actin cytoskeleton re-modelling are essential for RPE functionality and homeostasis^14,75^. However, as the ciliary functions of these molecules in ciliary signalling and IFT have already been shown to be crucial for RPE maturation, it is very likely that a combination of ciliary and alternative function results in defective functionality. This is supported by the fact that KD of genes essential for ciliary structure (*Ift20* and *Ift88*) showed a stronger phenotype in comparison to KD of *Bbs6*, despite a reduced number of primary cilia still being present in our cell model.

Taken together, we could demonstrate that ciliary dysfunction, caused by defective IFT, alters the apical membrane and changes protein expression and mitochondrial metabolism, leading to a phagocytosis defect in the RPE. Moreover, we identified new proteins involved in the phagocytic pathway. Our data further supports the connection between RPE metabolism and functionality, but for the first time also highlights the role of the primary cilium in this association. The connection between the primary cilium and metabolism is relevant beyond the RPE, revealing new insights into the ciliary influence on cellular processes, particularly in epithelial cells, which needs to be examined further.

## Methods

### Cell culture and siRNA transfection

RPE-J cells (ATCC) were cultivated at 32°C and 5% CO_2_ in Dulbecco’s MEM (4.5 g/L D-glucose, L-glutamine and pyruvate) supplemented with 4% foetal bovine serum (FBS; CELLect Gold, ICN), 1% non-essential amino acids, 1% HEPES and 1 % penicillin-streptomycin (“RPE-J medium”). To silence genes of interest and create knockdowns (KD), the cells were transfected every second day with specific DsiRNAs (Integrated DNA Technologies) using Lipofectamine RNAiMax (Thermo Fisher Scientific) according to the manufacturer’s protocol. As a control, a non-targeting control DsiRNA (NTC) was used that does not recognise any human, mouse or rat sequences. Cells were transfected one and three days after seeding and analysed four days after seeding, if not stated otherwise. To conduct immunofluorescence stainings, cells were seeded on glass coverslips. For trans-epithelial resistance (TEER) and capacitance (Ccl) measurements, cells were seeded on 12-well transwell inserts (PET 0.4 µm, transparent, Sarstedt) with a density of 3*10^5^ cells per transwell and cultivated in RPE-J medium with 10 mM nicotinamide, 0.25 mg/mL taurine, 0.01 nM retinoic acid for 8 days. To conduct Seahorse MitoStress assays, cells were cultivated in Seahorse XFp plates (Agilent, see dedicated paragraph “Metabolic flux analysis”).

### POS phagocytosis assay

Phagocytosis assays were conducted to measure bound and internalised fluorescein isothiocyanate (FITC)-tagged POS *in vitro*^31,95^. Cells were challenged with approximately 10 POS per cell in DMEM media containing no additives for 1.5 h and 4 h. After incubation, cells were washed three times with phosphate-buffered saline (PBS) with 0.2 mM CaCl_2_ and 1 mM MgCl_2_ (PBS-CM) to remove excess POS. Wells to measure internalised POS cells were treated with 0.4% Trypan Blue stain (GIBCO) for 10 min and washed twice. Cells were then fixed in ice-cold methanol for 10 min, rehydrated in PBS-CM for 10 min and then incubated with 4′,6-diamidino-2-phenylindole, dihydrochloride (DAPI; 1:400 in PBS-CM) for 15 min. After two additional washing steps, the fluorescence was measured using the TECAN Spark fluorescence plate reader. Bound POS were calculated by subtracting internalised POS from total POS.

### RNA isolation and real-time quantitative qPCR (RT-qPCR)

Total RNA was isolated using TRIzol™ reagent (ThermoFisher Scientific) according to the manufacturer’s instructions. cDNA was synthesised using the GoTaq® Probe 2-Step RT-qPCR System (Promega) and RT-qPCR was performed using iTaq Universal SYBR-Green® Supermix (BIO-RAD) and QuantStudio 3 system (ThermoFisher Scientific). Primers used are listed in Table S1.

### Cell lysis and western blotting

Cells were lysed using RIPA buffer (50 mM Tris HCl pH=8, 50 mM NaCl, 1% NP-40, 0.5% sodium deoxycholate, 0.1% SDS) supplemented with 1% protease and phosphotase inhibitor cocktail (SIGMA) on ice with periodical vortexing and resuspension, before centrifuging for 10 min at 4°C and 18,400 g. Lysate concentrations were determined via the bicinchoninic acid (BCA) assay (Sigma-Aldrich). We separated 100 µg of proteins with sodium dodecyl sulfate-polyacrylamide (SDS) gels consisting of 5% stacking and 10%/15% separating gels. PVDF membrane blotting was performed and antibodies were detected using the Odyssey® Fc Imaging System (LI-COR Bioscience) with the Image Studio Ver. 5.2 software. Antibodies used are listed in Table S2.

### LC-MS/MS Analysis

Cell lysates were prepared as described above. Proteins were precipitated using the methanol chloroform method, and solubilized proteins were subjected to in-solution tryptic cleavage as described earlier^96^. Samples were purified using StageTips. Briefly, the tip matrix was first equilibrated with 20 µL 0.5% acetic acid containing 80% acetonitrile (ACN) and then rinsed with 20 µL 0.5% acetic acid. Samples were loaded, before washing the matrix with 20 µL 0.5% acetic acid. Peptides were eluted in two steps: first with 20 µL 0.5% acetic acid containing 50% ACN and subsequently with 20 µL 0.5% acetic acid containing 80% ACN. LC-MS/MS analysis was performed as described before^97^. The MaxQuant software 1.6.1.0^98^ was used for label-free quantification. Trypsin/P was used as a digestion enzyme with a maximum of two missed cleavages. Cysteine carbamidomethylation was set as fixed modification, methionine oxidations and N-terminal acetylation as variable modifications. The first search peptide tolerance was set at 20 and the minimum ratio count at 2. Main search peptide tolerance was set to 4.5 ppm and the re-quantify option was selected. The Uniprot database (release 2023_02, #20,422 entries) was used, and the MaxQuant contaminant search was used to detect contaminants. Quantification requirements included a minimum of two peptides with at least one unique and one razor peptide with a minimum length of seven amino acids. Match between runs was performed with an alignment window of 20 min and a match time window of 0.7 min. The Perseus Framework software (version 1.6.15.0) was used for statistical analysis, including t-test, Significance A and ratio calculations^99^. Proteins which were statistically significant with the Student’s t-test (log p > 1.3) were categorised as Tier 2 hits, whereas proteins which were statistically significant with Student’s t-test and Significance A (p ≤ 0.05) were categorised as Tier 1 hits. Therefore, Tier 1 hits are automatically included in Tier 2 lists. VENN diagrams were created using InteractiVenn^100^ and GO-Term and Pathway (REACT and KEGG) analyses were performed using GetGo: A simple gene enrichment analysis tool^101^. For filtering the data for proteins of interest (phagocytosis, phago-/lysosomes, melanogenesis, visual cycle & cytoskeleton), we manually created lists based on Gene Ontology lists and published data (Table S3). MitoCarta 3.0 was used to filter the data for mitochondria-related proteins^68^.

### Transepithelial electrical resistance (TEER) and capacitance (Ccl) measurements

To assess apical membrane functionality, TEER and Ccl of RPE-J cells were measured using the cellZscope2 (Nano Analytics). TEER values are seen as a readout for cellular barrier integrity, which can be interpreted as an indicator of monolayer formation, whereas Ccl is a readout of the degree of the infolding of the apical membrane, symbolising the number of apical microvilli. Measurements were performed 4, 6 and 7 days after seeding. Cells were equilibrated for 10 min before measuring every 10 min for 5 times in total.

### Scanning electron microscopy (SEM)

RPE-J cells cultivated on transwells for 8 days were used for SEM. The cells were first fixed with 2.5% glutaraldehyde in 0.1 M cacodylate buffer with 0.1 M sucrose for 1 h at room temperature. The samples were then washed trice for 10 min with 0.2 M cacodylate buffer with 0.2 M sucrose. Cells were incubated with OsO_4_ in 0.2 M cacodylate buffer with 0.2 M sucrose for 1 h at room temperature, before being washed 6 times for 10 min with destilled H_2_O. After washing, the samples were dehydrated using an ethanol dehydration series with the following 10-min steps each: 1x with 30%, 1x with 50%, 1x with 70%, 1x with 80%, 2x with 96%, 2x with 100% ethanol. Critical point drying was performed hereafter using the BAL-TEC CPD 030. Samples were mounted onto conductive carbon adhesive on top of SEM specimen stubs and sputter coated with platinum (2.5 nm Pt, 15 mA at 2,0x10^-2^ mBar; CCU-010). Samples were stored in a desiccator and then imaged using a Zeiss GeminiSEM 560 with a landing energy of 1.5 kV.

### Immunofluorescence staining and microscopy

RPE-J cells cultivated on glass coverslips were fixed with 4% paraformaldehyde in PBS for 10 min at room temperature, before incubating them with 50 mM NH_4_Cl in PBS for 10 min to quench excess fixative. Cells were then permeabilised with PBS with 0.3% Triton-X100 (PBS-Tx) for 15 min before blocking for 1 h with Fish-Block (0.1% ovalbumin, 0.5% fish gelatine in PBS) with 0.3 % Triton-X100 (Fish-Block-Tx). Primary antibodies were diluted in Fish-Block-Tx and incubated over-night at 4°C. Cells were washed three times with PBS-Tx and then incubated with secondary antibodies and DAPI (1:400) diluted in Fish-Block-Tx for 1 h at room temperature. Cells were subsequently washed twice with PBS-Tx and once with PBS before mounting using Fluoromount-G™ Mounting Medium (ThermoFisher Scientific). To stain apical carbohydrate chains via surface labeling, cells were washed three times with HBSS-CM (0.2 mM Ca^2+^, 0.1 mM Mg^2+^) on ice, before incubating with FITC-labelled wheat germ agglutinin (WGA), diluted in 1:50 HBSS-CM for 45 min. Access antibody was removed with 2 rinses with HBSS-CM, before fixing the cells with trichloroacetic acid (1:10 in ddH_2_O) for 15 min. Cells were returned to RT and washed thrice with PBS-CM. Subsequently, cells were blocked with 1% BSA in PBS-CM for 30 min, incubated with DAPI (1:400 in 1% BSA) for 15 min and rinsed twice with PBS-CM, before mounting using Fluoromount-G™ Mounting Medium (ThermoFisher Scientific). Antibodies are listed in Table S2. Samples were imaged using the CTR6000 confocal microscope, equipped with the DM6000 B laser and the DFC360 FX monochrome digital camera, and LAS-X image software (Leica Microsystems).

### Primary Cilium quantification

Primary cilia percentage of the RPE-J KD cells was quantified. Only cells growing in a properly arranged monolayer, validated by ZO-1 staining, were counted and dividing cells were excluded in the quantification. Primary cilia were defined as a close co-staining of ARL13B and pericentrin (PCNT). 5 biological repetitions were conducted and for each at least 160 cells were counted.

### RPE morphology analysis

Morphological analysis and cell segmentation and was performed using Cellpose^102,103^, followed by a custom python-based pipeline. The analysis integrates the Cellpose segmentation output with the morphological analysis capabilities of Nyxus (v0.8.2) to enable automated analysis of cell morphology. The pipeline is accessible at [https://github.com/Kardelengenc/CellPose-Analysis.git]. Analysed parameters included: cell area, cell perimeter, aspect ratio, neighbour count, hexagonality score, polygonality score (refer to Github for specific parameters and calculations). The analysis was performed using microscopy images of RPE-J cells stained for ZO-1.

### Metabolic flux analysis

RPE-J cells were cultivated in Seahorse XFp FluxPak PDL Cell Culture Miniplates (Agilent Technologies) and equilibrated for 1 h without CO_2_ before starting the assay in Seahorse XF base medium minimal DMEM without phenol red supplemented with 1 mM pyruvate, 2 mM glutamine and 10 mM glucose, and adjusted to pH7.4. The XFp Cell Mito Stress Test Kit (Agilent Technologies) was used with the following mitochondrial inhibitors: oligomycin (1.5 μM), FCCP (carbonyl cyanide 4-(trifluoromethoxy)-phenylhydrazone, 0.5 µM) and rotenone/antimycin A (0.5 µM)^104^. Oxygen Consumption Rate (OCR) and Extracellular Acidification Rate (ECAR) were quantified with the Seahorse Report Generator online software (Agilent Technologies). The original Seahorse program was modified to meet our experiment requirements and is described in Hamieh et al. ^105^. Values were first normalised to protein concentration, determined using the Bradford assay. Then, a second normalisation step was applied to eliminate the variation between assay plates. Therefore, the mean values of each parameter and treatment were divided by the sum of NTC basal respiration and non-mitochondrial oxygen consumption values at the start of the experiment for each experiment. The energy map was generated by plotting ECAR (x-axis) against OCR (y-axis) values in basic (mean value of first 3 measurements) and stressed (mean value after adding FCCP) conditions. The bioenergetics health index was calculated as described in Hamieh et al.^105^.

### Statistical analysis

All experiments were conducted a minimum of 3 times if not stated otherwise. Statistical analysis of the data was performed using GraphPad Prism with suggested tests and post-hoc analysis. Specific tests used for statistical analysis are stated in the respective figure legends. Following significance levels were used: * p < 0.05; ** p < 0.01; *** p < 0.001; **** p < 0.0001.

## Acknowledgements

The authors would especially like to thank Viola Kretschmer, Emily Belina, Jonathan Becker, Alina Frei, Vanessa Maißl, Kardelen Genç, Rike Hähnel-Ewerling and the rest of the AG May-Simera lab for their valuable input and discussion during the preparation of this manuscript. Further we thank Christoph Sieber for technical assistance. This work is dedicated to the memory of Elisabeth Sehn, a valued colleague who is sorely missed.

## Author contributions

<u>Conceptualization:</u> Peter Andreas Matthiessen, Emeline F. Nandrot, Helen Louise May-Simera

<u>Data curation:</u> Peter Andreas Matthiessen, Karsten Boldt, Emeline F. Nandrot, Helen Louise May-Simera

<u>Formal analysis:</u> Peter Andreas Matthiessen

<u>Funding acquisition:</u> Peter Andreas Matthiessen, Helen Louise May-Simera

<u>Investigation:</u> Peter Andreas Matthiessen, Lotta Elisabeth Wagner, Karsten Boldt, Gunnar Glaßer, Ingo Lieberwirth, Emeline F. Nandrot, Helen Louise May-Simera

<u>Methodology:</u> Peter Andreas Matthiessen, Lotta Elisabeth Wagner, Karsten Boldt, Gunnar Glaßer, Ingo Lieberwirth, Emeline F. Nandrot, Helen Louise May-Simera

<u>Project administration:</u> Emeline F. Nandrot, Helen Louise May-Simera

<u>Resources:</u> Emeline F. Nandrot, Helen Louise May-Simera

<u>Supervision:</u> Emeline F. Nandrot, Helen Louise May-Simera

<u>Visualization:</u> Peter Andreas Matthiessen, Emeline F. Nandrot

<u>Writing – original draft:</u> Peter Andreas Matthiessen, Helen Louise May-Simera

<u>Writing – review & editing:</u> Peter Andreas Matthiessen, Lotta Elisabeth Wagner, Gunnar Glaßer, Emeline F. Nandrot, Helen Louise May-Simera

---

**Figure S1** RPE-J cell model characterisation.

(A) siRNA knockdown efficiency of *Ift20*, *Ift88* and *Bbs*6 in RPE-J cells was determined via RT-qPCR. *Rplp0* was used as housekeeping gene and data were normalised to NTC (n = 3). (B) IFT88 knockdown efficiency was validated on protein level via western blot analysis. GAPDH was used as housekeeping protein and data were normalised to NTC (n = 7). (C-D) Primary cilia were imaged and ciliation quantified via immunofluorescence staining. Primary cilia were defined as a close co-staining of ARL13B and PCNT. White squares display area of zoom in images. White arrows in zoom-in images show primary cilia examples. ARL13B staining is shown in magenta, PCNT in grey, ZO-1 in green and nucleus staining DAPI in blue (n = 5). Scale bars represent 10 µm. (E) mRNA expression of ciliary genes *Arl13b*, *Cep290* and *Mks1* in RPE-J KD cells was determined via RT-qPCR. *Rplp0* was used as housekeeping gene and data were normalised to NTC (dashed line, n = 3-5). One-way ANOVA test with Dunnet’s multiple comparison (A, B, D), unpaired t-test (B) and Two-way ANOVA test with Sidak’s multiple comparison (E) were used as statistical analysis with the following p-values: * p < 0.05; ** p < 0.01; *** p < 0.001; **** p < 0.0001. Data are shown as mean ± SEM.

**Figure S2** Cell morphology remains unchanged upon ciliary dysfunction.

Cell morphology of RPE-J KD cells was analysed for following criteria: (A) Cell area. (B) Cell perimeter. (C) Aspect ratio. (D) Number of neighbouring cells. Cells at peripheral region with 0-2 neighbours were excluded. (E) Average hexagonality score. Cells with score of 0 were excluded. (F) Standard deviation of hexagonality. Cells with score of 0 were excluded. (G) Average polygonality score. Cells with score of 0 were excluded. NTC was used as control. Data is shown as mean ± SEM (n = 4). Kruskal-Wallis test with Dunn’s multiple comparison was used as statistical analysis with the following p-values: * p < 0.05; ** p < 0.01; *** p < 0.001; **** p < 0.0001.

**Figure S3** Changes in transepithelial electrical resistance, capacitance and apical carbohydrate chains upon ciliary dysfunction.

TEER (A) and Ccl (B) were measured with RPE-J KD cells after 5, 7 and 8 days of cultivation. NTC was used as control. Data is shown as mean ± SEM (n = 3). Two-way ANOVA test with Tukey’s multiple comparison was used as statistical analysis with the following p-values: * p < 0.05; ** p < 0.01; *** p < 0.001; **** p < 0.0001. TEER, transepithelial electrical resistance; Ccl, capacitance. (C) Apical carbohydrate chains were visualised using WGA-FITC immunofluorescence staining. White squares display area of zoom in images and X-Z axis cross sections. WGA-FITC staining is shown in green, nuclei stained with DAPI in blue. Scale bars represent 10 µm.

**Figure S4** RPE-J *Paox*, *Plekhb2* and *Tk2* KD validation.

SiRNA knockdown efficiency of RPE-J cells was determined via RT-qPCR. *Rplp0* was used as housekeeping gene and NTC as control. Data are shown as mean ± SEM (n = 3). One-way ANOVA test with Dunnet’s multiple comparison was used as statistical analysis as statistical analysis with following p-values: * p < 0.05; ** p < 0.01; *** p < 0.001; **** p < 0.0001.

**Figure S5** Gene ontology and pathway analysis of individual KD cells reveal altered cellular pathway upon ciliary gene knockdown.

(A-C) Bar charts display mis-regulated GO terms of individual RPE-J ciliary gene knockdown cells as indicated. (D-F) Bar charts display pathway terms mis-regulated in individual RPE-J ciliary gene knockdown cells as indicated. Top 20 hits displayed for *Ift20* and *Ift88* KD cells, all hits displayed for *Bbs6* KD cells. Analysis was performed using Tier 2 hits and GetGo (n = 4).

**Figure S6** Ciliary dysfunction alters ECAR and results in mitochondrial metabolism shift in RPE cells.

OCR (A) and ECAR (B) profiles of RPE-J KD cells (n = 5-7). (C) ECAR/OCR energy map RPE-J KD cells under basic (circles) and stressed conditions (squares). Data is shown as mean ± SEM (n = 5-7). (D) Bioenergetic health index of RPE-J KD cells. NTC was used as control. Data is shown as mean ± SEM (n = 5-7). Unpaired t-test with Welch’s correction (A, B) and unpaired t-test (D) was used as statistical analysis as statistical analysis with following p-values: * p < 0.05; ** p < 0.01; *** p < 0.001; **** p < 0.0001.

**Figure S7** Full sized western blots.

Original western blots cropped to membrane size of (A) Figure S1B, (B) Figure 2D and (C) Figure 5C. Black squares display cropped areas shown in single figures.

**Figure S8** Full sized western blots probing phagocytosis pathway proteins in RPE-J *Ift20* KD cells.

Original western blots cropped to membrane size of Figure 1C, left panel. Black squares display cropped areas shown in single figures.

**Figure S9** Full sized western blots probing phagocytosis pathway proteins in RPE-J *Ift88* KD cells.

Original western blots cropped to membrane size of Figure 1C, middle panel. Black squares display cropped areas shown in single figures.

**Figure S10** Full sized western blots probing phagocytosis pathway proteins in RPE-J *Bbs6* KD cells.

Original western blots cropped to membrane size of Figure 1C, right panel. Black squares display cropped areas shown in single figures.

