## Supplementary Files for "Loss of ciliary proteins IFT20 and IFT88 results in defective phagocytosis and metabolism in the RPE"

### Supplementary information

Supplementary Figures S1-S10

Tables S1-S3

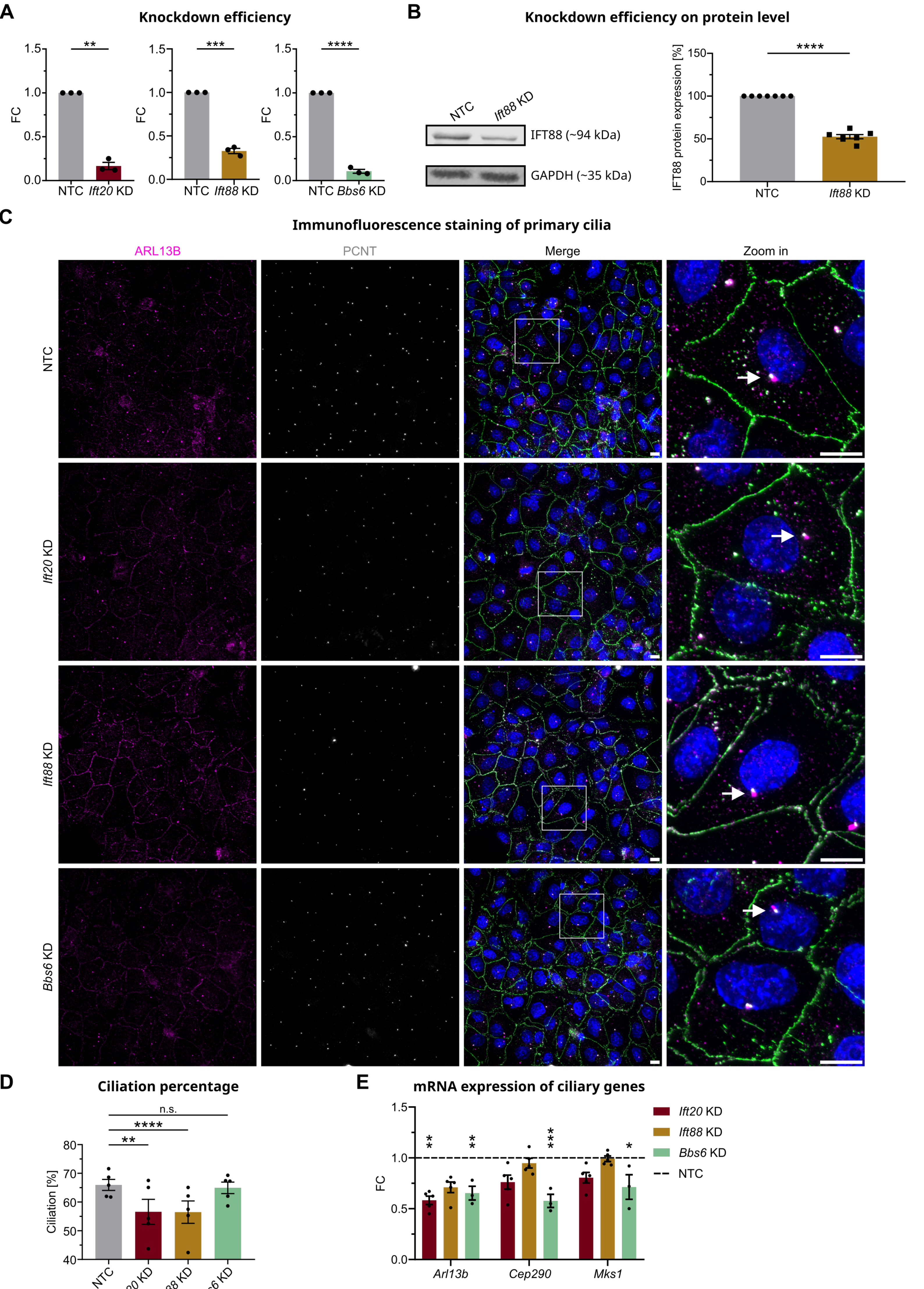
A
Knockdown efficiency
FC
NTC
Ift20 KD
Ift88 KD
Bbs6 KD
B
Knockdown efficiency on protein level
NTC
Ift88 KD
IFT88 (~94 kDa)
GAPDH (~35 kDa)
IFT88 protein expression [%]
C
Immunofluorescence staining of primary cilia
ARL13B
PCNT
Merge
Zoom in
D
Ciliation percentage
Ciliation [%]
n.s.
E
mRNA expression of ciliary genes
Arl13b
Cep290
Mks1
NTC

Figure S1

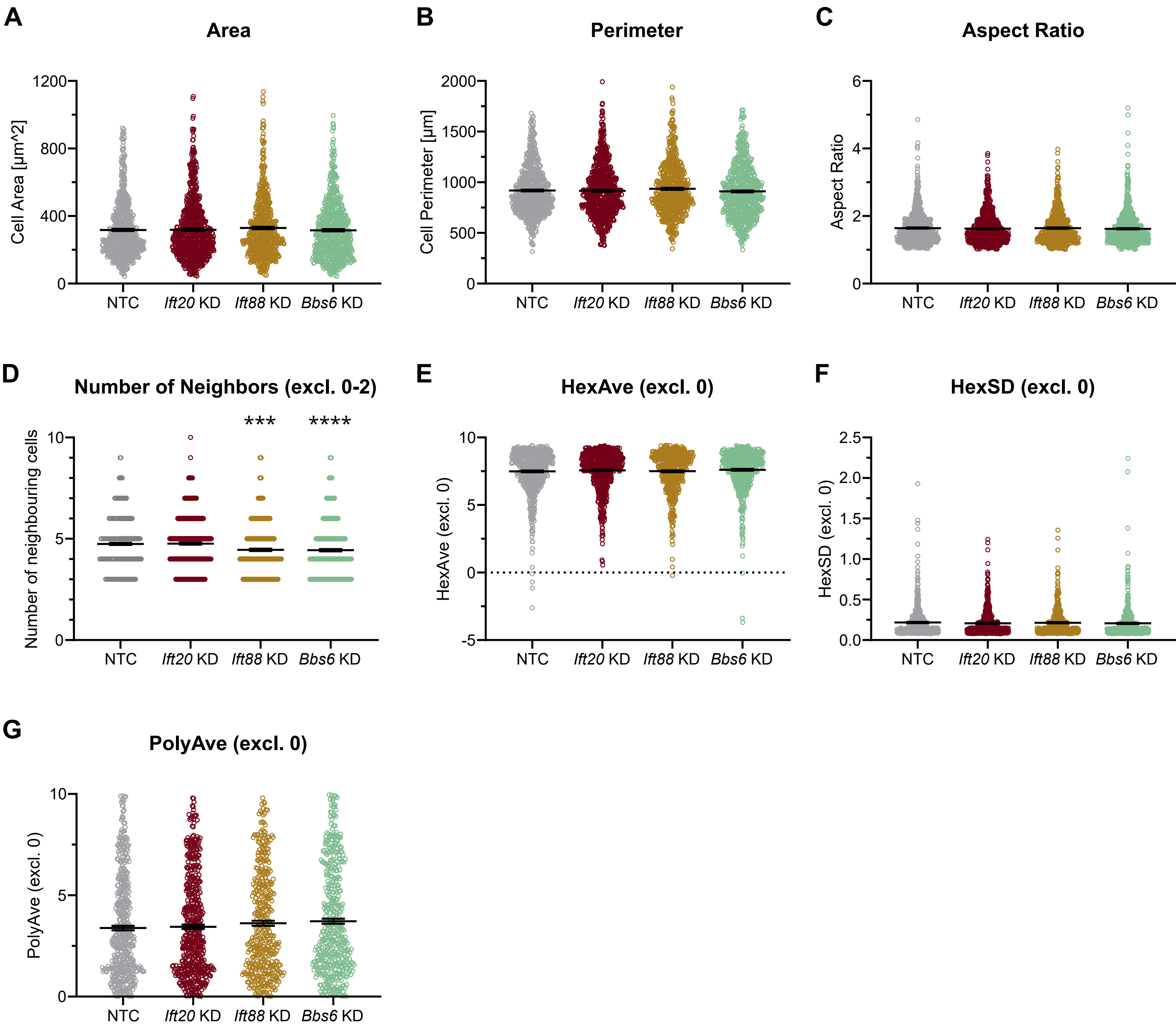

A
Area
Cell Area [µm^2]
1200
800
400
0
NTC
Ift20 KD
Ift88 KD
Bbs6 KD
B
Perimeter
Cell Perimeter [µm]
2000
1500
1000
500
0
C
Aspect Ratio
Aspect Ratio
6
4
2
0
D
Number of Neighbors (excl. 0-2)
Number of neighbouring cells
***
****
10
5
0
E
HexAve (excl. 0)
HexAve (excl. 0)
-5
F
HexSD (excl. 0)
HexSD (excl. 0)
2.5
2.0
1.5
1.0
0.5
0.0
G
PolyAve (excl. 0)
PolyAve (excl. 0)


Figure S2

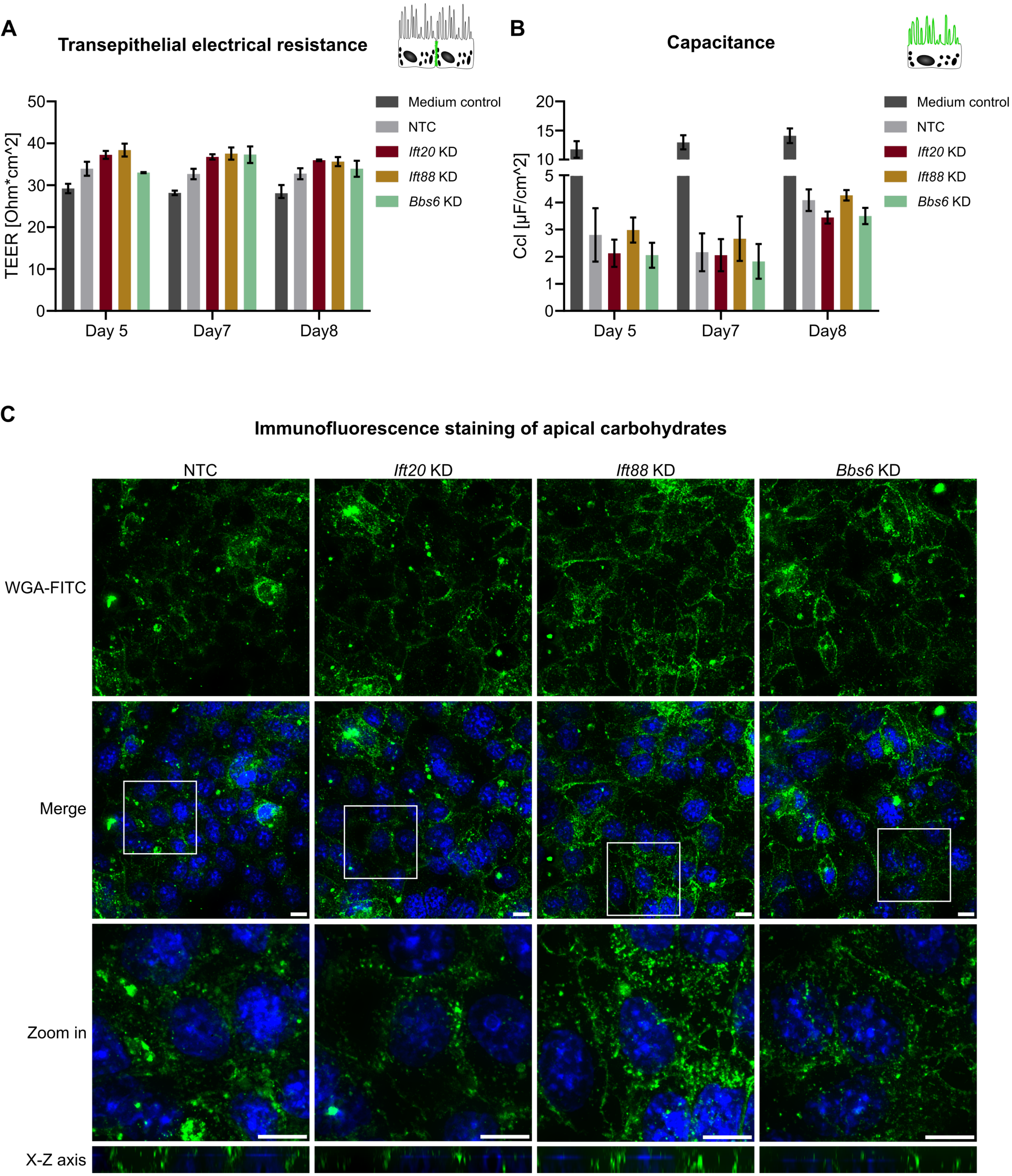
A
Transepithelial electrical resistance
TEER [Ohm*cm^2]
Day 5
Day7
Day8
Medium control
NTC
Ift20 KD
Ift88 KD
Bbs6 KD
B
Capacitance
Ccl [µF/cm^2]
C
Immunofluorescence staining of apical carbohydrates
NTC
Ift20 KD
Ift88 KD
Bbs6 KD
WGA-FITC
Merge
Zoom in
X-Z axis

Figure S3

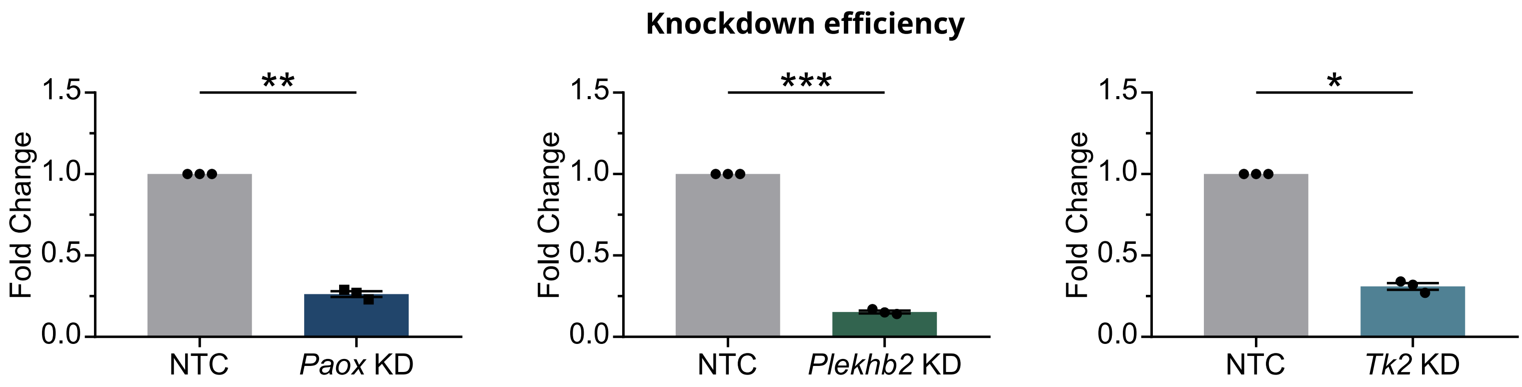
Knockdown efficiency
1.5
1.0
0.5
0.0
Fold Change
**
NTC
Paox KD
***
Plekhb2 KD
*
Tk2 KD

Figure S4

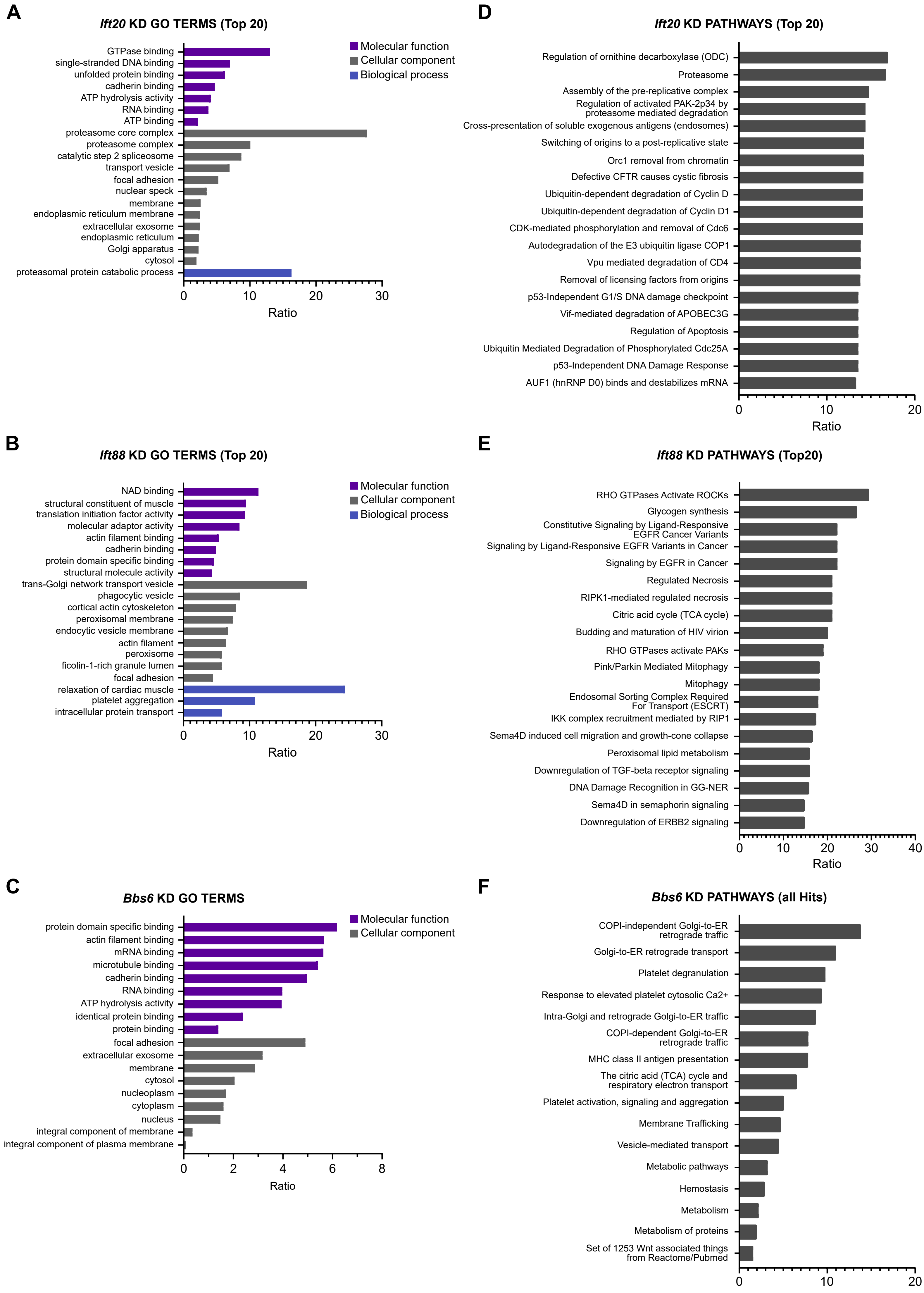
A
Ift20 KD GO TERMS (Top 20)
Molecular function
Cellular component
Biological process
GTPase binding
single-stranded DNA binding
unfolded protein binding
cadherin binding
ATP hydrolysis activity
RNA binding
ATP binding
proteasome core complex
proteasome complex
catalytic step 2 spliceosome
transport vesicle
focal adhesion
nuclear speck
membrane
endoplasmic reticulum membrane
extracellular exosome
endoplasmic reticulum
Golgi apparatus
cytosol
proteasomal protein catabolic process
0
10
20
30
Ratio
D
Ift20 KD PATHWAYS (Top 20)
Regulation of ornithine decarboxylase (ODC)
Proteasome
Assembly of the pre-replicative complex
Regulation of activated PAK-2p34 by proteasome mediated degradation
Cross-presentation of soluble exogenous antigens (endosomes)
Switching of origins to a post-replicative state
Orc1 removal from chromatin
Defective CFTR causes cystic fibrosis
Ubiquitin-dependent degradation of Cyclin D
Ubiquitin-dependent degradation of Cyclin D1
CDK-mediated phosphorylation and removal of Cdc6
Autodegradation of the E3 ubiquitin ligase COP1
Vpu mediated degradation of CD4
Removal of licensing factors from origins
p53-Independent G1/S DNA damage checkpoint
Vif-mediated degradation of APOBEC3G
Regulation of Apoptosis
Ubiquitin Mediated Degradation of Phosphorylated Cdc25A
p53-Independent DNA Damage Response
AUF1 (hnRNP D0) binds and destabilizes mRNA
0
10
20
Ratio
B
Ift88 KD GO TERMS (Top 20)
Molecular function
Cellular component
Biological process
NAD binding
structural constituent of muscle
translation initiation factor activity
molecular adaptor activity
actin filament binding
cadherin binding
protein domain specific binding
structural molecule activity
trans-Golgi network transport vesicle
phagocytic vesicle
cortical actin cytoskeleton
peroxisomal membrane
endocytic vesicle membrane
actin filament
peroxisome
ficolin-1-rich granule lumen
focal adhesion
relaxation of cardiac muscle
platelet aggregation
intracellular protein transport
0
10
20
30
Ratio
E
Ift88 KD PATHWAYS (Top20)
RHO GTPases Activate ROCKs
Glycogen synthesis
Constitutive Signaling by Ligand-Responsive EGFR Cancer Variants
Signaling by Ligand-Responsive EGFR Variants in Cancer
Signaling by EGFR in Cancer
Regulated Necrosis
RIPK1-mediated regulated necrosis
Citric acid cycle (TCA cycle)
Budding and maturation of HIV virion
RHO GTPases activate PAKs
Pink/Parkin Mediated Mitophagy
Mitophagy
Endosomal Sorting Complex Required For Transport (ESCRT)
IKK complex recruitment mediated by RIP1
Sema4D induced cell migration and growth-cone collapse
Peroxisomal lipid metabolism
Downregulation of TGF-beta receptor signaling
DNA Damage Recognition in GG-NER
Sema4D in semaphorin signaling
Downregulation of ERBB2 signaling
0
10
20
30
40
Ratio
C
Bbs6 KD GO TERMS
Molecular function
Cellular component
protein domain specific binding
actin filament binding
mRNA binding
microtubule binding
cadherin binding
RNA binding
ATP hydrolysis activity
identical protein binding
protein binding
focal adhesion
extracellular exosome
membrane
cytosol
nucleoplasm
cytoplasm
nucleus
integral component of membrane
integral component of plasma membrane
0
2
4
6
8
Ratio
F
Bbs6 KD PATHWAYS (all Hits)
COPI-independent Golgi-to-ER retrograde traffic
Golgi-to-ER retrograde transport
Platelet degranulation
Response to elevated platelet cytosolic Ca2+
Intra-Golgi and retrograde Golgi-to-ER traffic
COPI-dependent Golgi-to-ER retrograde traffic
MHC class II antigen presentation
The citric acid (TCA) cycle and respiratory electron transport
Platelet activation, signaling and aggregation
Membrane Trafficking
Vesicle-mediated transport
Metabolic pathways
Hemostasis
Metabolism
Metabolism of proteins
Set of 1253 Wnt associated things from Reactome/Pubmed
0
10
20
Ratio

Figure S5

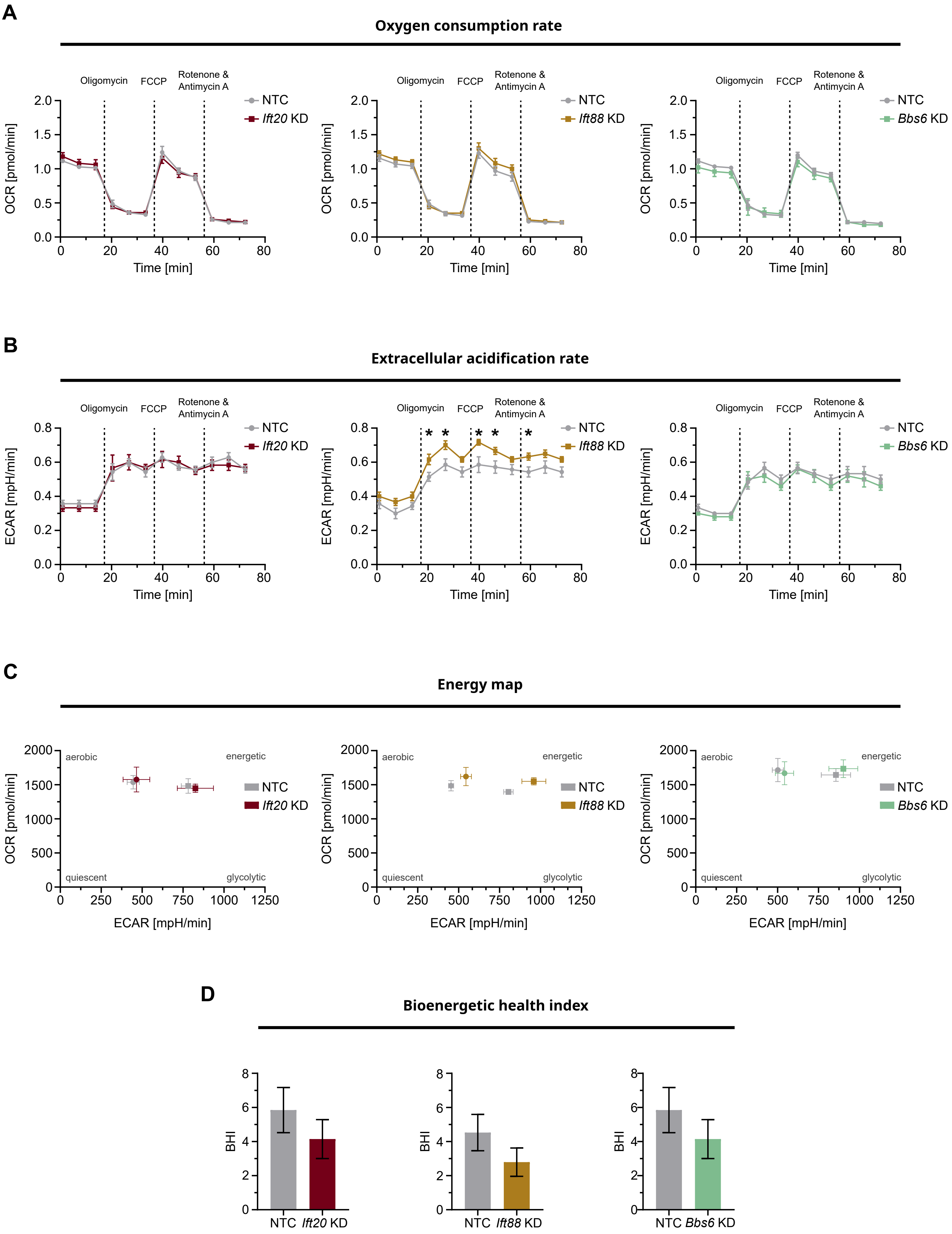
A
Oxygen consumption rate
Oligomycin
FCCP
Rotenone & Antimycin A
OCR [pmol/min]
Time [min]
NTC
Ift20 KD
Ift88 KD
Bbs6 KD
B
Extracellular acidification rate
ECAR [mpH/min]
*
C
Energy map
aerobic
energetic
quiescent
glycolytic
D
Bioenergetic health index
BHI

Figure S6

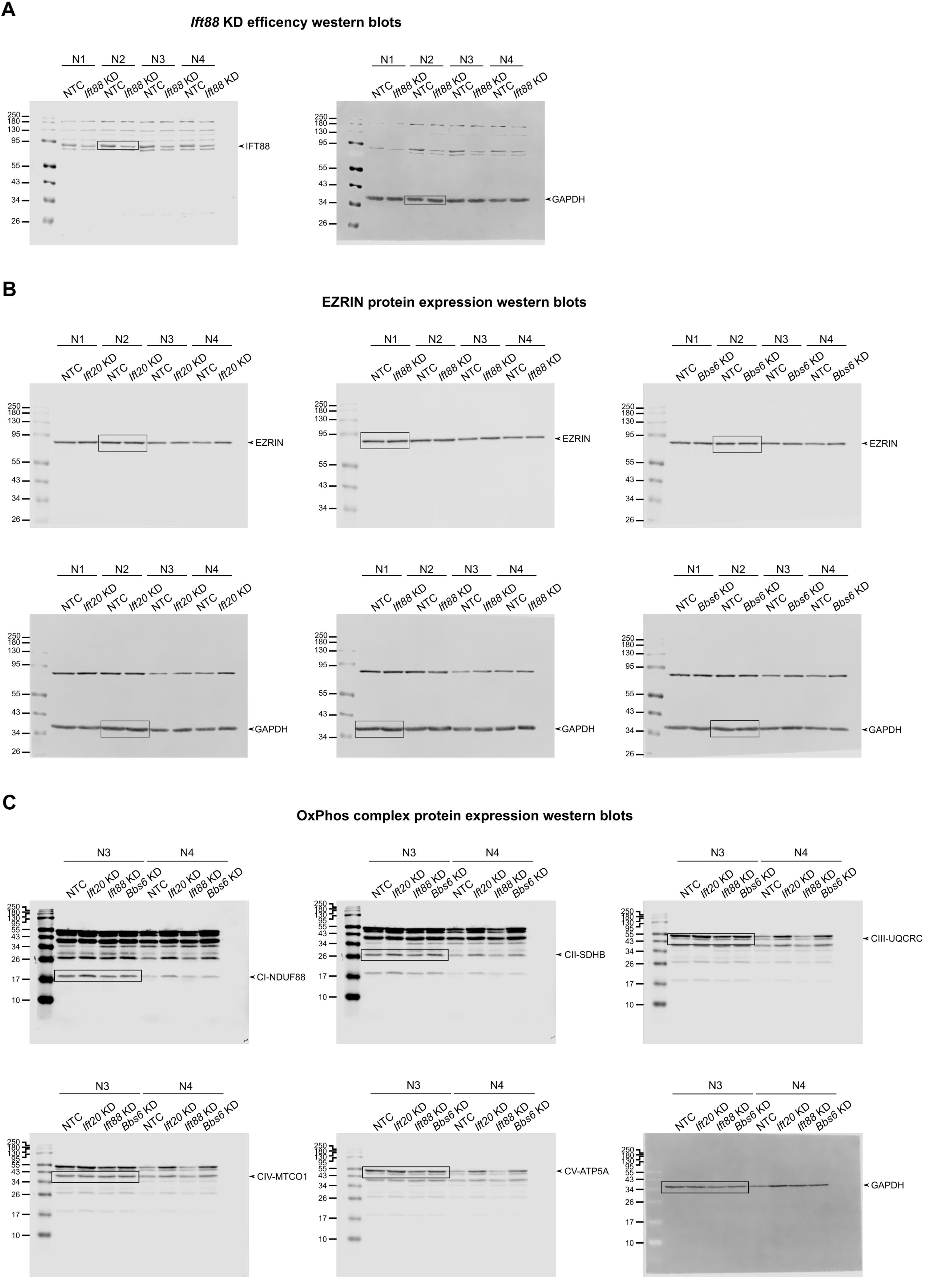
A
Ift88 KD efficency western blots
N1 N2 N3 N4
NTC Ift88 KD NTC Ift88 KD NTC Ift88 KD NTC Ift88 KD
250 180 130 95 55 43 34 26
IFT88
GAPDH
B
EZRIN protein expression western blots
NTC Ift20 KD NTC Ift20 KD NTC Ift20 KD NTC Ift20 KD
NTC Ift88 KD NTC Ift88 KD NTC Ift88 KD NTC Ift88 KD
NTC Bbs6 KD NTC Bbs6 KD NTC Bbs6 KD NTC Bbs6 KD
EZRIN
GAPDH
C
OxPhos complex protein expression western blots
N3 N4
NTC Ift20 KD Ift88 KD Bbs6 KD NTC Ift20 KD Ift88 KD Bbs6 KD
250 180 130 95 55 43 34 26 17 10
CI-NDUF88
CII-SDHB
CIII-UQCRC
CIV-MTCO1
CV-ATP5A
GAPDH

Figure S7

#### *Ift20* KD phagocytosis protein expression western blots

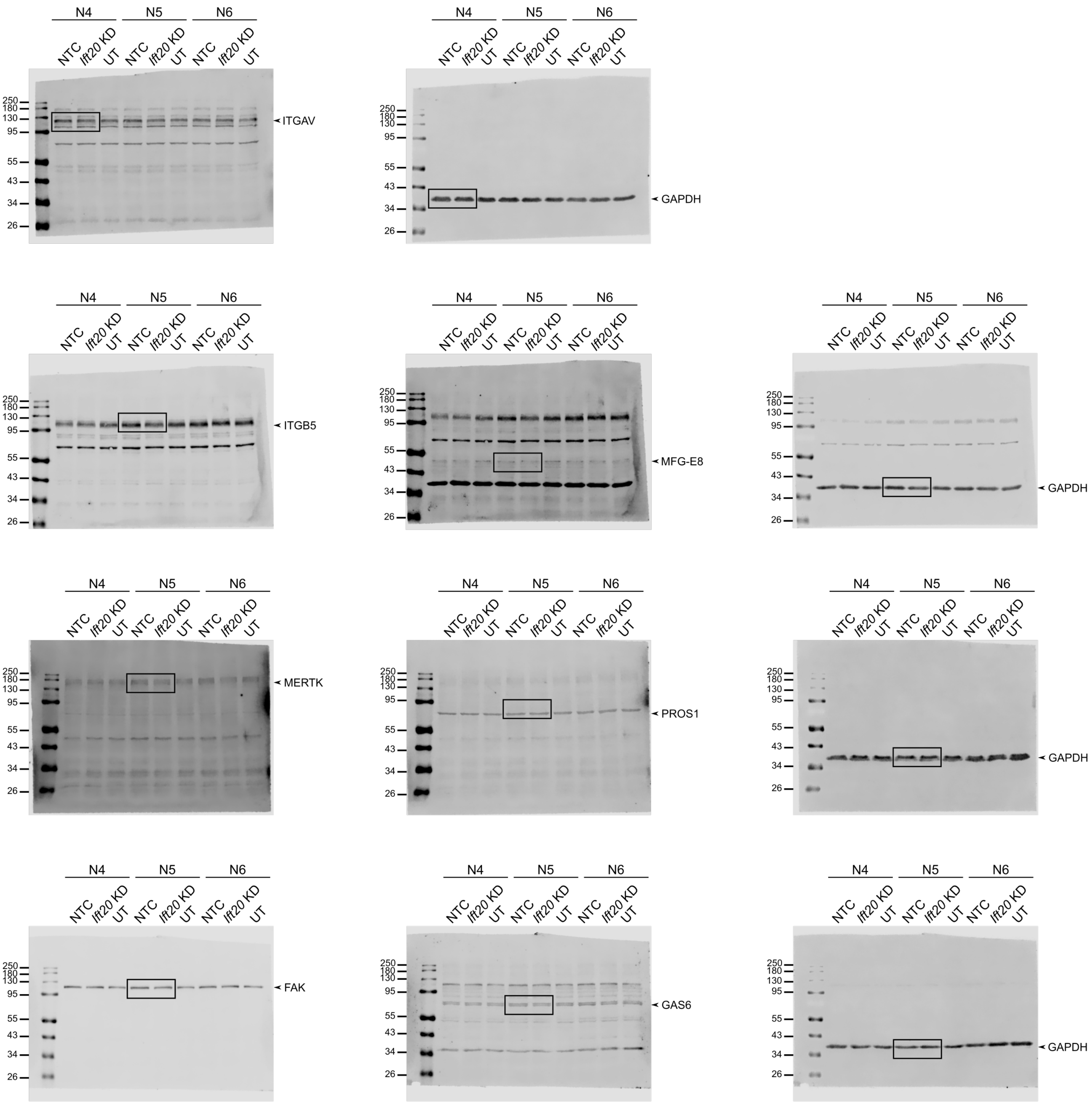


Figure S8

**_Ift88_ KD phagocytosis protein expression western blots**

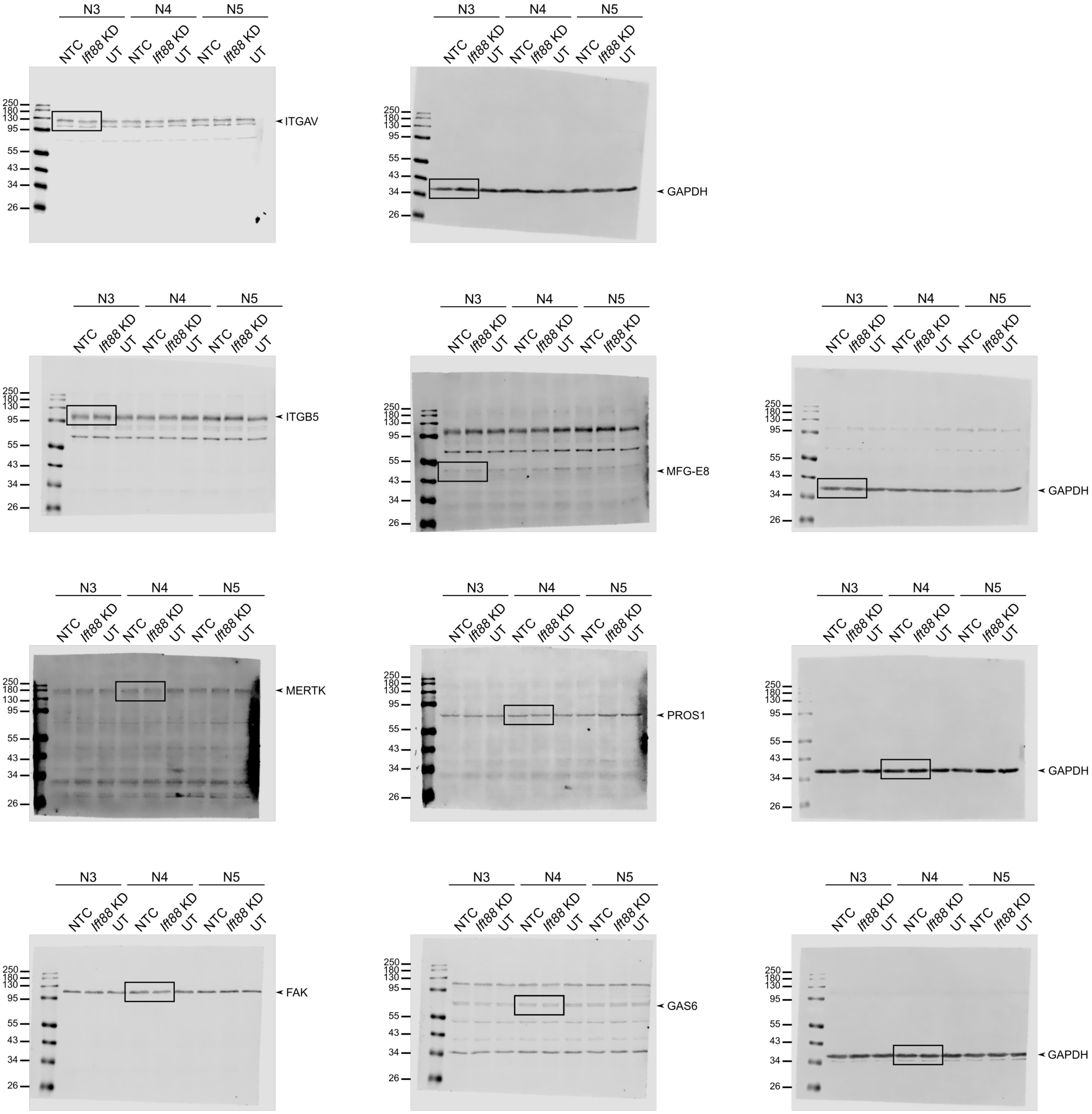


Figure S9

#### *Bbs6* KD phagocytosis protein expression western blots

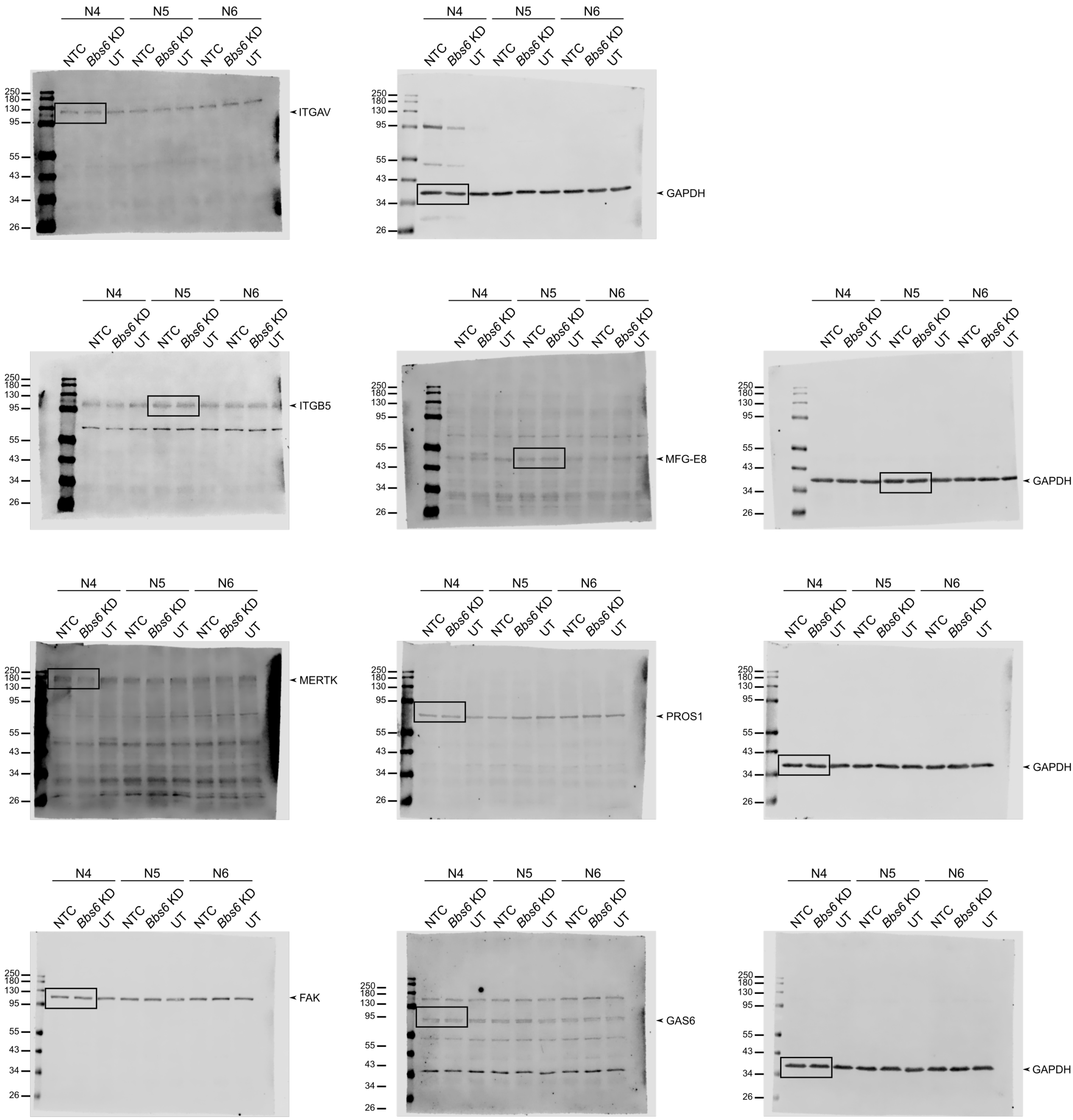


Figure S10

Table S1: List of primers.

| Primer | Sequence | |
|---|---|---|
| | **Fwd** | **Rvs** |
| *Ift20* | TGAACAAGCTTCGGGTGTTG | AAAGTCCTTGCACTCTTCCTTGA |
| *Ift88* | ACCAGGCTGTAGACACATT | TTCTCGTAGTCACCATTTG |
| *Bbs6* | ATGTAGATCTTGTCCTGTGCC | ATAGGGAACCAATAGGCGTTG |
| *Mfge8* | CTCGTCTGTGTATATGGGCTTC | ATCTTCCGCAGAAAGTCCAC |
| *Ptk2* | GAAAGCAGTAATGAGCCAACC | TGAGGCGAAATCCATAGCAG |
| *Pros1* | ACTCCTGGAAGTTACCACTG | TCACATTCGAAGTCTCCTGG |
| *Gas6* | AGGAGGCTAGAGAGGTGTTC | GTGCATTGGTCAGGTAAGTTC |
| *Mertk* | GGAAAGATGGAAAGGAACTGC | GATGTACGACCCATTGTCTGAG |
| *Itgav* | GACAAGCTCACTCCCATCACT | AGTAAAATGTGAGCCTGCCGA |
| *Itgb5* | GGTTTCGGGTCTTTTGTTGAC | ACTCTGTCTGTGAGAGGCAG |
| *Mks1* | GAAGGACCTCATAGACTTGGC | GCGGATTCATTTTGGTACAGG |
| *Cep290* | AAGTACAGGGACGTCTTGCA | GAGCCCCTCCATCTGTTCTT |
| *Arl13b* | ACAGACGGAACTTGATGGGA | GTCCTTCTCCGCAGTCTTCT |
| *Nd4* | GGCAACCAAACAGAACGCTT | GGTGTGTTGTGAGGGAGAGG |
| *Cox4i* | TTCGCTGAGATGAACAAGGG | GATCAAAGGTATGAGGGATGGG |
| *Atp6* | CGTAATTACAGGCTTCCGACA | GGCTGCGTCTTCTTCTTCAT |
| *Plekhb2* | TATGTTGGCTCCGCAATCCT | CCATAGCCGTAGACCTCAGG |
| *Paox* | AGAGTGTTGTGTGAGCGGTA | TGGTCTTCACTGGCTTGTCA |
| *Tk2* | TGCGATTCTGTCTGAGTGGT | TGACTTTCTCCTCTTCCCGG |
| *Rplp0* | GACAATGGCAGCATCTACAG | CATTGATGATGGAGTGAGGC |

Table S2: List of antibodies.

| Antibodies | Company | Product ID | Dilution | Host | Application |
|---|---|---|---|---|---|
| ITGB5 | Proteintech | 28543-1-AP | 1:1000 | rabbit | Western blot |
| FAK | Proteintech | 12636-1-AP | 1:1000 | rabbit | Western blot |
| GAPDH | SIGMA | G8795 | 1:2000 | mouse | Western blot |
| GAPDH | Cell Signaling | 5176 | 1:2000 | rabbit | Western blot |
| GAS6 | Proteintech | 13795-1-AP | 1:1000 | rabbit | Western blot |
| ITGAV | BD Biosciences | 4711 | 1:400 | mouse | Western blot |
| MERTK | R&D Systems | AF591 | 1:1000 | goat | Western blot |
| MFG-E8 | R&D Systems | AF2805 | 1:1000 | goat | Western blot |
| OxPhos Rodent AB Cocktail | Invitrogen | 45-8099 | 1:250 | mouse | Western blot |
| PROTEIN-S | Proteintech | 16910-1-AP | 1:1000 | rabbit | Western blot |
| EZRIN | SIGMA | E8897 | 1:50 | mouse | ICC |
| ARL13B | NeuroMab | ab136648 | 1:400 | mouse | ICC |
| PERICENTRIN | Abcam | Ab4448 | 1:400 | rabbit | ICC |
| WGA-FITC | SIGMA | L4895 | 1:50 | Triticum vulgaris | ICC |
| ZO-1-488 | Invitrogen | 339188 | 1:100 | mouse | ICC |

Table S3: Lists of functionality- and cytoskeleton-related targets used to filter mass spectrometry data.

| Phagocytosis | Phago-/Lysosome | Melanogenesis | Visual Cycle | Cytoskeleton |
|---|---|---|---|---|
| Anxa2 | Abcb6 | Adcy1 | Aipl1 | Akap13 |
| Anxa5 | Adrb2 | Adcy2 | Arrb1 | Als2 |
| Arap1 | Akt1 | Adcy3 | Asic2 | Als2Cl |
| Cd36 | Aktip | Adcy4 | Cabp4 | Arhgef1 |
| Cd81 | Ambra1 | Adcy5 | Cds2 | Arhgef10 |
| Clta | Ankfy1 | Adcy6 | Cngb1 | Arhgef10L |
| Cltb | Ap1g1 | Adcy7 | Gnat1 | Arhgef11 |
| Cltc | Ap3b1 | Adcy8 | Gnat2 | Arhgef12 |
| Gas6 | Ap3d1 | Adcy9 | Gnb1 | Arhgef15 |
| Itga5 | Ap4m1 | Ap1 | Gngt1 | Arhgef16 |
| Itgb5 | Appl1 | Ap2 | Gpr52 | Arhgef17 |
| Lyar | Appl2 | Ap3 | Gpr88 | Arhgef18 |
| Mertk | Arf1 | Asip | Grk1 | Arhgef19 |
| Mfge8 | Arfip2 | Atrn | Guca1a | Arhgef2 |
| Myh2 | Arl8b | Bcl2 | Guca1b | Arhgef25 |
| Pkcd | Asb14 | Bloc1S1 | Lrat | Arhgef26 |
| Pros1 | Atg10 | Bloc1S2 | Opn1mw | Arhgef28 |
| Ptk2 | Atg101 | Calm1 | Opn1sw | Arhgef3 |
| Rac1 | Atg12 | Calm2 | Opn3 | Arhgef33 |
| Rack1 | Atg13 | Calm3 | Opn4 | Arhgef35 |
| Tbp | Atg14 | Calml3 | Opn5 | Arhgef37 |
| Tlr4 | Atg16l1 | Calml5 | Pcp2 | Arhgef38 |
| Uso1 | Atg16l2 | Calml6 | Pde6c | Arhgef39 |
| | Atg2a | Camk2A | Pde6d | Arhgef4 |
| | Atg2b | Camk2B | Rcvrn | Arhgef40 |
| | Atg3 | Camk2D | Rdh | Arhgef5 |
| | Atg4a | Camk2G | Rdh11 | Arhgef6 |
| | Atg4a-ps | Creb1 | Rgr | Arhgef7 |
| | Atg4b | Creb3 | Rho | Arhgef9 |
| | Atg4c | Creb3L1 | Rpe65 | Bcr |
| | Atg4d | Creb3L2 | Rrh | Dnmbp |
| | Atg5 | Creb3L3 | Rs1 | Ect2 |
| | Atg7 | Creb3L4 | Slc24a2 | Ect2L |
| | Atg9a | Crebbp | Slc24a4 | Farp1 |
| | Atg9b | Ctnnb1 | | Farp2 |
| | Atp13a2 | Dct | | Fgd1 |
| | Atp2a2 | Dvl1 | | Fgd2 |
| | Aup1 | Dvl2 | | Fgd3 |
| | Bag3 | Dvl3 | | Fgd4 |
| | Becn1 | Edn1 | | Fgd5 |
| | Becn2 | Ednrb | | Fgd6 |
| | Bin1 | Ep300 | | Itsn1 |
| | Bloc1s1 | Fzd1 | | Itsn2 |
| | Bloc1s2 | Fzd10 | | Kalrn |

| | | | |
|---|---|---|---|
| Borcs5 | Fzd2 | | Mcf2 |
| Borcs6 | Fzd3 | | Mcf2L |
| Borcs7 | Fzd4 | | Mcf2L2 |
| Borcs8 | Fzd5 | | Net1 |
| C9orf72 | Fzd6 | | Ngef |
| Cacng2 | Fzd7 | | Obscn |
| Cacng3 | Fzd8 | | Plekhg1 |
| Cacng4 | Fzd9 | | Plekhg2 |
| Cacng5 | Gnai1 | | Plekhg3 |
| Cacng7 | Gnai2 | | Plekhg4 |
| Cacng8 | Gnai3 | | Plekhg4B |
| Calcoco2 | Gnao1 | | Plekhg5 |
| Calm1 | Gnaq | | Plekhg6 |
| Ccdc91 | Gnas | | Prex1 |
| Cd81 | Gpr143 | | Prex2 |
| Cdx2 | Gsk3B | | Rasgrf1 |
| Chmp1a | Hras | | Rasgrf2 |
| Chmp1b | Icam1 | | Spata13 |
| Chmp1b2 | Kit | | Tiam1 |
| Chmp2a | Kitlg | | Tiam2 |
| Chmp2b | Kras | | Trio |
| Chmp3 | Lef1 | | Vav1 |
| Chmp4b | Litlg | | Vav2 |
| Chmp4c | Map2K1 | | Vav3 |
| Chmp5 | Map2K2 | | 1700012B09Rik |
| Chmp6 | Mapk1 | | 2700049A03Rik |
| Chmp7 | Mapk14 | | 4933427D14Rik |
| Clec16a | Mapk3 | | Aaas |
| Cln3 | Mc1R | | Aamp |
| Clu | Mitf | | Abca2 |
| Coro1a | Mlana | | Abcc3 |
| Ctsd | Nras | | Abi1 |
| Dennd3 | Nurr1 | | Abi2 |
| Dnajc16 | Oca2 | | Abl1 |
| Dtx3l | Onec2 | | Abl2 |
| Eea1 | Pah | | Ablim1 |
| Efnb1 | Pax3 | | Ablim2 |
| Ehd3 | Pik3Cb | | Ablim3 |
| Ehmt2 | Plcb1 | | Abr |
| Elapor1 | Plcb2 | | Abra |
| Emc6 | Plcb3 | | Abraxas2 |
| Entpd4 | Plcb4 | | Acaa2 |
| Entpd4b | Pmel | | Acaca |
| Epg5 | Pomc | | Acads |
| Ephb2 | Prkaca | | Ackr2 |
| Epm2a | Prkacb | | Acot13 |
| Fez1 | Prkacg | | Acta1 |

| | | | |
|---|---|---|---|
| Fez2 | Prkca | | Acta2 |
| Fhip1b | Prkcb | | Actb |
| Flcn | Prkcg | | Actbl2 |
| Fnip1 | Prkx | | Actc1 |
| Fth1 | Raf1 | | Acte1 |
| Ftl1 | Rps6Kb1 | | Actg1 |
| Fyco1 | Slc45A2 | | Actg2 |
| Gaa | Sox10 | | Actl10 |
| Gabarap | Tcf7 | | Actl11 |
| Gabarapl1 | Tcf7L1 | | Actl7A |
| Gabarapl2 | Tcf7L2 | | Actl7B |
| Gak | Tyr | | Actl9 |
| Gba | Tyrp1 | | Actn1 |
| Gcc2 | Usf1 | | Actn2 |
| Gga3 | Wnt1 | | Actn3 |
| Glmp | Wnt10A | | Actn4 |
| Gnptab | Wnt10B | | Actr10 |
| Gpr137b | Wnt11 | | Actr1A |
| Gprasp1 | Wnt16 | | Actr1B |
| Grn | Wnt2 | | Actr2 |
| Hap1 | Wnt2B | | Actr3 |
| Hgs | Wnt3 | | Actr3B |
| Hook1 | Wnt3A | | Actr6 |
| Hook2 | Wnt4 | | Actr8 |
| Hook3 | Wnt5A | | Actrt1 |
| Hrg | Wnt5B | | Actrt2 |
| Hspa8 | Wnt6 | | Actrt3 |
| Htt | Wnt7A | | Adam17 |
| Ift20 | Wnt7B | | Adam8 |
| Ift88 | Wnt8A | | Adcy10 |
| Igtp | Wnt8B | | Adcy8 |
| Irgm1 | Wnt9A | | Adcy9 |
| Irgm2 | Wnt9B | | Add1 |
| Jmy | | | Add2 |
| Kics2 | | | Add3 |
| Kif13a | | | Adora2A |
| Kptn | | | Adrb2 |
| Kxd1 | | | Afap1 |
| Lamp1 | | | Afap1L1 |
| Lamp2 | | | Afg2A |
| Lamtor1 | | | Afg2B |
| Lamtor4 | | | Agbl1 |
| Lamtor5 | | | Agbl2 |
| Laptm4b | | | Agbl3 |
| Laptm5 | | | Agbl4 |
| Lhcgr | | | Agbl5 |
| Lipa | | | Agtpbp1 |

| | | | |
|---|---|---|---|
| Lmbrd1 | | | Ahi1 |
| Lrba | | | Ahnak |
| Lrrk2 | | | Aif1 |
| Lrsam1 | | | Aif1L |
| Lyst | | | Ajuba |
| M6pr | | | Ak1 |
| Map1lc3a | | | Ak5 |
| Map1lc3b | | | Ak6 |
| Mapk15 | | | Ak8 |
| Mcoln1 | | | Akap12 |
| Meak7 | | | Akap13 |
| Mfn2 | | | Akap14 |
| Mfsd1 | | | Akap5 |
| Mgrn1 | | | Akap9 |
| Moap1 | | | Akna |
| Mreg | | | Akt1 |
| Mtm1 | | | Akt3 |
| Mtmr3 | | | Aldoa |
| Mtor | | | Aldob |
| Mvb12a | | | Alms1 |
| Myo7a | | | Alox8 |
| Nbr1 | | | Alpk1 |
| Ncoa4 | | | Als2 |
| Ndp | | | Ambra1 |
| Nedd4 | | | Amot |
| Nsfl1c | | | Amotl1 |
| Nupr1 | | | Amotl2 |
| Osbpl7 | | | Amph |
| P2rx7 | | | Anapc5 |
| Pacs2 | | | Anapc7 |
| Pcdhga3 | | | Ang |
| Peg3 | | | Ang2 |
| Phf23 | | | Ang3 |
| Pik3c2a | | | Ang4 |
| Pik3c2b | | | Ang5 |
| Pik3c3 | | | Ang6 |
| Pik3r4 | | | Ank1 |
| Pikfyve | | | Ank2 |
| Pink1 | | | Ank3 |
| Pip4k2a | | | Ankfn1 |
| Pip4k2b | | | Ankra2 |
| Pip4k2c | | | Ankrd23 |
| Pla2g5 | | | Ankrd26 |
| Plekhf1 | | | Ankrd53 |
| Plekhf2 | | | Ankrd7 |
| Plekhm1 | | | Anks1B |
| Ppp3cb | | | Anln |

| | | | |
|---|---|---|---|
| Prkcd | | | Anxa1 |
| Prkd1 | | | Anxa11 |
| Psen1 | | | Anxa2 |
| Pxk | | | Apbb1Ip |
| Rab12 | | | Apbb3 |
| Rab14 | | | Apc |
| Rab19 | | | Apc2 |
| Rab1a | | | Apex1 |
| Rab1b | | | Apkc |
| Rab20 | | | Apoe |
| Rab23 | | | App |
| Rab24 | | | Appbp2 |
| Rab31 | | | Appl1 |
| Rab32 | | | Aqp0 |
| Rab33b | | | Arap1 |
| Rab34 | | | Arap3 |
| Rab38 | | | Arc |
| Rab39 | | | Arfgef2 |
| Rab3gap1 | | | Arhgap18 |
| Rab3gap2 | | | Arhgap21 |
| Rab43 | | | Arhgap24 |
| Rab5a | | | Arhgap26 |
| Rab7 | | | Arhgap32 |
| Rab7b | | | Arhgap33 |
| Ralb | | | Arhgap35 |
| Rb1cc1 | | | Arhgap39 |
| Rbsn | | | Arhgap4 |
| Rhob | | | Arhgap6 |
| Rilp | | | Arhgef10 |
| Rnf167 | | | Arhgef18 |
| Rnf186 | | | Arhgef2 |
| Rnf5 | | | Arhgef5 |
| Rock2 | | | Arhgef7 |
| Rpn2 | | | Arl13B |
| Rraga | | | Arl2 |
| Rragc | | | Arl2Bp |
| Rtn4 | | | Arl3 |
| Rubcn | | | Arl6 |
| Rubcnl | | | Arl6Ip5 |
| Rufy4 | | | Arl8A |
| Scarb2 | | | Arl8B |
| Scfd1 | | | Armc9 |
| Scyl2 | | | Arpc1A |
| Sec22b | | | Arpc1B |
| Sh3bp4 | | | Arpc2 |
| Sh3glb1 | | | Arpc3 |
| Slamf8 | | | Arpc4 |

| | | | |
|---|---|---|---|
| Slc15a4 | | | Arpc5 |
| Slc4a7 | | | Arpc5L |
| Slc9a9 | | | Arsj |
| Smcr8 | | | Asap1 |
| Smpd1 | | | Asb2 |
| Smurf1 | | | Aspm |
| Snap29 | | | Atat1 |
| Snapin | | | Atf3 |
| Snx16 | | | Atf4 |
| Snx18 | | | Atf5 |
| Snx27 | | | Atf6B |
| Snx3 | | | Atg14 |
| Snx30 | | | Atg16L1 |
| Snx4 | | | Atg5 |
| Snx7 | | | Atg7 |
| Sorl1 | | | Atm |
| Sort1 | | | Atp12A |
| Spg11 | | | Atp2B4 |
| Sqstm1 | | | Atp6V0D1 |
| Srpx | | | Atp6V1D |
| Stbd1 | | | Atxn10 |
| Sting1 | | | Atxn7 |
| Stx12 | | | Aunip |
| Stx17 | | | Aurka |
| Stx7 | | | Aurkb |
| Stx8 | | | Aurkc |
| Syk | | | Auts2 |
| Synpo2 | | | Avil |
| Syt11 | | | Axdnd1 |
| Syt7 | | | Axin1 |
| Szt2 | | | Axin2 |
| Tasl | | | Axl |
| Tbc1d12 | | | Azin1 |
| Tbc1d14 | | | B9D1 |
| Tbc1d25 | | | B9D2 |
| Tbc1d5 | | | Bag2 |
| Tcirg1 | | | Bag3 |
| Tecpr1 | | | Baiap2 |
| Tex264 | | | Baiap2L1 |
| Tgfbrap1 | | | Barx2 |
| Tlr7 | | | Bbln |
| Tlr9 | | | Bbof1 |
| Tmem106b | | | Bbs1 |
| Tmem175 | | | Bbs2 |
| Tmem39a | | | Bbs4 |
| Tmem41b | | | Bbs5 |
| Tom1 | | | Bbs7 |

| | | | |
|---|---|---|---|
| Tpcn1 | | | Bbs9 |
| Tpcn2 | | | Bc048507 |
| Traf6 | | | Bcar1 |
| Trak1 | | | Bcas2 |
| Trak2 | | | Bcas3 |
| Trex1 | | | Bccip |
| Trp53inp1 | | | Bcl10 |
| Trp53inp2 | | | Bcl2L1 |
| Tsg101 | | | Bcl2L10 |
| Ubqln1 | | | Bcl2L11 |
| Ubqln2 | | | Bcl3 |
| Ubqln4 | | | Becn1 |
| Ubxn2a | | | Bex4 |
| Ubxn2b | | | Bex6 |
| Ubxn6 | | | Bfsp1 |
| Uevld | | | Bfsp2 |
| Ulk1 | | | Bicd1 |
| Ulk2 | | | Bicd2 |
| Ulk3 | | | Bicdl1 |
| Unc13b | | | Bin1 |
| Unc93b1 | | | Bin2 |
| Uvrag | | | Bin3 |
| Vamp8 | | | Birc5 |
| Vcp | | | Birc6 |
| Vmp1 | | | Birc7 |
| Vps11 | | | Bloc1S2 |
| Vps16 | | | Bloc1S6 |
| Vps18 | | | Bmerb1 |
| Vps33a | | | Bmf |
| Vps35 | | | Bmyc |
| Vps39 | | | Bnip2 |
| Vps41 | | | Bod1 |
| Vps4a | | | Bora |
| Vps4b | | | Borcs5 |
| Vps53 | | | Bpifb4 |
| Vps54 | | | Braf |
| Vti1a | | | Brca1 |
| Washc1 | | | Brca2 |
| Wdfy3 | | | Brcc3 |
| Wdr45 | | | Brk1 |
| Wdr45b | | | Brsk1 |
| Wdr81 | | | Brsk2 |
| Wipi1 | | | Brwd1 |
| Wipi2 | | | Bsn |
| Zfyve1 | | | Bub1B |
| Zfyve16 | | | Bud31 |
| Zfyve26 | | | Bves |

| | | | | |
|---|---|---|---|---|
| | | | | Bysl |
| | | | | C2Cd3 |
| | | | | C2Cd5 |
| | | | | Cabcoco1 |
| | | | | Cabyr |
| | | | | Cadps2 |
| | | | | Calb1 |
| | | | | Calb2 |
| | | | | Calcoco2 |
| | | | | Cald1 |
| | | | | Calm1 |
| | | | | Calm2 |
| | | | | Calm3 |
| | | | | Calml3 |
| | | | | Camk2B |
| | | | | Camsap1 |
| | | | | Camsap2 |
| | | | | Camsap3 |
| | | | | Cap1 |
| | | | | Capg |
| | | | | Capn10 |
| | | | | Capn6 |
| | | | | Capn7 |
| | | | | Caprin2 |
| | | | | Capza1 |
| | | | | Capza1B |
| | | | | Capza2 |
| | | | | Capza3 |
| | | | | Capzb |
| | | | | Car |
| | | | | Carmil1 |
| | | | | Carmil2 |
| | | | | Casp1 |
| | | | | Casp14 |
| | | | | Cass4 |
| | | | | Catip |
| | | | | Cbx1 |
| | | | | Cbx3 |
| | | | | Cby1 |
| | | | | Cc2D1A |
| | | | | Cc2D2A |
| | | | | Ccar2 |
| | | | | Ccdc102A |
| | | | | Ccdc103 |
| | | | | Ccdc110 |
| | | | | Ccdc112 |
| | | | | Ccdc113 |

| | | | | |
|---|---|---|---|---|
| | | | | Ccdc116 |
| | | | | Ccdc117 |
| | | | | Ccdc120 |
| | | | | Ccdc124 |
| | | | | Ccdc13 |
| | | | | Ccdc14 |
| | | | | Ccdc141 |
| | | | | Ccdc146 |
| | | | | Ccdc15 |
| | | | | Ccdc153 |
| | | | | Ccdc178 |
| | | | | Ccdc18 |
| | | | | Ccdc181 |
| | | | | Ccdc187 |
| | | | | Ccdc22 |
| | | | | Ccdc28B |
| | | | | Ccdc38 |
| | | | | Ccdc39 |
| | | | | Ccdc40 |
| | | | | Ccdc42 |
| | | | | Ccdc50 |
| | | | | Ccdc57 |
| | | | | Ccdc6 |
| | | | | Ccdc61 |
| | | | | Ccdc63 |
| | | | | Ccdc65 |
| | | | | Ccdc66 |
| | | | | Ccdc68 |
| | | | | Ccdc69 |
| | | | | Ccdc77 |
| | | | | Ccdc78 |
| | | | | Ccdc8 |
| | | | | Ccdc81 |
| | | | | Ccdc85B |
| | | | | Ccdc88A |
| | | | | Ccdc88B |
| | | | | Ccdc88C |
| | | | | Ccdc92 |
| | | | | Ccdc96 |
| | | | | Cchcr1 |
| | | | | Ccin |
| | | | | Ccn5 |
| | | | | Ccna1 |
| | | | | Ccna2 |
| | | | | Ccnb1 |
| | | | | Ccnb1-Ps |
| | | | | Ccnb2 |

| | | | | |
|---|---|---|---|---|
| | | | | Ccnd1 |
| | | | | Ccnd2 |
| | | | | Ccnd3 |
| | | | | Ccne1 |
| | | | | Ccne2 |
| | | | | Ccnf |
| | | | | Ccnj |
| | | | | Ccnjl |
| | | | | Ccno |
| | | | | Ccp110 |
| | | | | Ccsap |
| | | | | Ccser2 |
| | | | | Cct2 |
| | | | | Cct3 |
| | | | | Cct4 |
| | | | | Cct5 |
| | | | | Cct6A |
| | | | | Cct7 |
| | | | | Cct8 |
| | | | | Cd180 |
| | | | | Cd274 |
| | | | | Cd2Ap |
| | | | | Cd86 |
| | | | | Cdc14A |
| | | | | Cdc14B |
| | | | | Cdc16 |
| | | | | Cdc20 |
| | | | | Cdc25B |
| | | | | Cdc27 |
| | | | | Cdc42 |
| | | | | Cdc42Bpa |
| | | | | Cdc42Bpb |
| | | | | Cdc42Bpg |
| | | | | Cdc42Ep1 |
| | | | | Cdc42Ep2 |
| | | | | Cdc42Ep3 |
| | | | | Cdc42Ep4 |
| | | | | Cdc42Ep5 |
| | | | | Cdc42Se1 |
| | | | | Cdc42Se2 |
| | | | | Cdc45 |
| | | | | Cdc6 |
| | | | | Cdc7 |
| | | | | Cdca8 |
| | | | | Cdh1 |
| | | | | Cdh10 |
| | | | | Cdh11 |

| | | | | |
|---|---|---|---|---|
| | | | | Cdh12 |
| | | | | Cdh13 |
| | | | | Cdh14 |
| | | | | Cdh15 |
| | | | | Cdh16 |
| | | | | Cdh17 |
| | | | | Cdh18 |
| | | | | Cdh19 |
| | | | | Cdh2 |
| | | | | Cdh20 |
| | | | | Cdh21 |
| | | | | Cdh22 |
| | | | | Cdh23 |
| | | | | Cdh24 |
| | | | | Cdh25 |
| | | | | Cdh26 |
| | | | | Cdh3 |
| | | | | Cdh4 |
| | | | | Cdh5 |
| | | | | Cdh6 |
| | | | | Cdh7 |
| | | | | Cdh8 |
| | | | | Cdh9 |
| | | | | Cdk1 |
| | | | | Cdk10 |
| | | | | Cdk16 |
| | | | | Cdk2 |
| | | | | Cdk2Ap2 |
| | | | | Cdk5 |
| | | | | Cdk5Rap2 |
| | | | | Cdk5Rap3 |
| | | | | Cdk6 |
| | | | | Cdkl2 |
| | | | | Cdkl5 |
| | | | | Cdkn1B |
| | | | | Cenatac |
| | | | | Cenpe |
| | | | | Cenpf |
| | | | | Cenpq |
| | | | | Cenpu |
| | | | | Cenpv |
| | | | | Cep104 |
| | | | | Cep112 |
| | | | | Cep120 |
| | | | | Cep126 |
| | | | | Cep128 |
| | | | | Cep131 |

| | | | | |
|---|---|---|---|---|
| | | | | Cep135 |
| | | | | Cep152 |
| | | | | Cep162 |
| | | | | Cep164 |
| | | | | Cep170 |
| | | | | Cep170B |
| | | | | Cep19 |
| | | | | Cep192 |
| | | | | Cep20 |
| | | | | Cep250 |
| | | | | Cep290 |
| | | | | Cep295 |
| | | | | Cep295Nl |
| | | | | Cep350 |
| | | | | Cep41 |
| | | | | Cep43 |
| | | | | Cep44 |
| | | | | Cep55 |
| | | | | Cep57 |
| | | | | Cep57L1 |
| | | | | Cep63 |
| | | | | Cep68 |
| | | | | Cep70 |
| | | | | Cep72 |
| | | | | Cep76 |
| | | | | Cep78 |
| | | | | Cep83 |
| | | | | Cep85 |
| | | | | Cep85L |
| | | | | Cep89 |
| | | | | Cep95 |
| | | | | Cep97 |
| | | | | Cetn1 |
| | | | | Cetn2 |
| | | | | Cetn3 |
| | | | | Cetn4 |
| | | | | Cfap100 |
| | | | | Cfap107 |
| | | | | Cfap119 |
| | | | | Cfap126 |
| | | | | Cfap141 |
| | | | | Cfap144 |
| | | | | Cfap157 |
| | | | | Cfap161 |
| | | | | Cfap20 |
| | | | | Cfap206 |
| | | | | Cfap210 |

| | | | | |
|---|---|---|---|---|
| | | | | Cfap221 |
| | | | | Cfap251 |
| | | | | Cfap276 |
| | | | | Cfap298 |
| | | | | Cfap300 |
| | | | | Cfap36 |
| | | | | Cfap410 |
| | | | | Cfap43 |
| | | | | Cfap44 |
| | | | | Cfap45 |
| | | | | Cfap47 |
| | | | | Cfap52 |
| | | | | Cfap53 |
| | | | | Cfap54 |
| | | | | Cfap57 |
| | | | | Cfap58 |
| | | | | Cfap61 |
| | | | | Cfap68 |
| | | | | Cfap69 |
| | | | | Cfap70 |
| | | | | Cfap73 |
| | | | | Cfap74 |
| | | | | Cfap77 |
| | | | | Cfap90 |
| | | | | Cfap91 |
| | | | | Cfap95 |
| | | | | Cfap96 |
| | | | | Cfl1 |
| | | | | Cfl2 |
| | | | | Cgn |
| | | | | Cgnl1 |
| | | | | Champ1 |
| | | | | Chd3 |
| | | | | Chd4 |
| | | | | Chek1 |
| | | | | Chmp1A |
| | | | | Chmp1B |
| | | | | Chmp1B2 |
| | | | | Chmp2A |
| | | | | Chmp2B |
| | | | | Chmp3 |
| | | | | Chmp4B |
| | | | | Chmp4C |
| | | | | Chmp5 |
| | | | | Chmp6 |
| | | | | Chmp7 |
| | | | | Chodl |

| | | | | |
|---|---|---|---|---|
| | | | | Chp1 |
| | | | | Chrm1 |
| | | | | Chrm2 |
| | | | | Chrm3 |
| | | | | Chrm4 |
| | | | | Chrm5 |
| | | | | Ciao1 |
| | | | | Ciao2B |
| | | | | Cib1 |
| | | | | Cib2 |
| | | | | Cibar1 |
| | | | | Cibar2 |
| | | | | Cilk1 |
| | | | | Cimap1A |
| | | | | Cimap1B |
| | | | | Cimap1C |
| | | | | Cimap1D |
| | | | | Cimap3 |
| | | | | Cimip2A |
| | | | | Cimip2B |
| | | | | Cimip2C |
| | | | | Cingulin |
| | | | | Cir1 |
| | | | | Cit |
| | | | | Ckap2 |
| | | | | Ckap2L |
| | | | | Ckap4 |
| | | | | Ckap5 |
| | | | | Clasp1 |
| | | | | Clasp2 |
| | | | | Cldn1 |
| | | | | Cldn10 |
| | | | | Cldn11 |
| | | | | Cldn12 |
| | | | | Cldn13 |
| | | | | Cldn14 |
| | | | | Cldn15 |
| | | | | Cldn16 |
| | | | | Cldn17 |
| | | | | Cldn18 |
| | | | | Cldn19 |
| | | | | Cldn2 |
| | | | | Cldn20 |
| | | | | Cldn21 |
| | | | | Cldn22 |
| | | | | Cldn23 |
| | | | | Cldn24 |

| | | | | |
|---|---|---|---|---|
| | | | | Cldn3 |
| | | | | Cldn4 |
| | | | | Cldn5 |
| | | | | Cldn6 |
| | | | | Cldn7 |
| | | | | Cldn8 |
| | | | | Cldn9 |
| | | | | Clic4 |
| | | | | Clic5 |
| | | | | Clip1 |
| | | | | Clip2 |
| | | | | Clip3 |
| | | | | Clip4 |
| | | | | Clk3 |
| | | | | Clmp |
| | | | | Clns1A |
| | | | | Clrn1 |
| | | | | Clta |
| | | | | Cltc |
| | | | | Clu |
| | | | | Cluap1 |
| | | | | Clxn |
| | | | | Cmah |
| | | | | Cnn1 |
| | | | | Cnn2 |
| | | | | Cnn3 |
| | | | | Cnp |
| | | | | Cnr1 |
| | | | | Cntln |
| | | | | Cntrl |
| | | | | Cntrob |
| | | | | Cobl |
| | | | | Corin |
| | | | | Coro1A |
| | | | | Coro1B |
| | | | | Coro1C |
| | | | | Coro2B |
| | | | | Cotl1 |
| | | | | Cpap |
| | | | | Cpeb1 |
| | | | | Cplane2 |
| | | | | Cr1L |
| | | | | Cracr2A |
| | | | | Crb1 |
| | | | | Crb2 |
| | | | | Crb3 |
| | | | | Creb1 |

| | | | | |
|---|---|---|---|---|
| | | | | Crhbp |
| | | | | Crk |
| | | | | Crmp1 |
| | | | | Crocc |
| | | | | Crocc2 |
| | | | | Cryab |
| | | | | Csnk1A1 |
| | | | | Csnk1D |
| | | | | Cspp1 |
| | | | | Csrp1 |
| | | | | Csrp3 |
| | | | | Cstpp1 |
| | | | | Ctdp1 |
| | | | | Ctnna1 |
| | | | | Ctnna2 |
| | | | | Ctnna3 |
| | | | | Ctnnal1 |
| | | | | Ctnnb1 |
| | | | | Ctnnbl1 |
| | | | | ***Ctnnd1*** |
| | | | | Ctsc |
| | | | | Ctsh |
| | | | | Cttn |
| | | | | Cttnbp2 |
| | | | | Cttnbp2Nl |
| | | | | Cul3 |
| | | | | Cul7 |
| | | | | Cxcr2 |
| | | | | Cyb5D1 |
| | | | | Cyba |
| | | | | Cylc1 |
| | | | | Cyld |
| | | | | Cyp2A4 |
| | | | | Cyp2A5 |
| | | | | Cypt1 |
| | | | | Cys1 |
| | | | | Cyth4 |
| | | | | D1Pas1 |
| | | | | D7Ertd443E |
| | | | | Daam1 |
| | | | | Dag1 |
| | | | | Dapk1 |
| | | | | Dapk3 |
| | | | | Daw1 |
| | | | | Daxx |
| | | | | Dbh |
| | | | | Dbn1 |

| | | | | |
|---|---|---|---|---|
| | | | | Dbnl |
| | | | | Dbt |
| | | | | Dcaf1 |
| | | | | Dcaf12 |
| | | | | Dcaf13 |
| | | | | Dcdc2A |
| | | | | Dcdc2B |
| | | | | Dcdc2C |
| | | | | Dclk2 |
| | | | | Dclre1B |
| | | | | Dctn1 |
| | | | | Dctn2 |
| | | | | Dctn3 |
| | | | | Dctn4 |
| | | | | Dctn5 |
| | | | | Dctn6 |
| | | | | Dcun1D5 |
| | | | | Dcx |
| | | | | Dcxr |
| | | | | Ddhd2 |
| | | | | Ddr2 |
| | | | | Ddx11 |
| | | | | Ddx3X |
| | | | | Ddx6 |
| | | | | Ddx60 |
| | | | | Def6 |
| | | | | Dennd1C |
| | | | | Dennd2A |
| | | | | Des |
| | | | | Deup1 |
| | | | | Dgkg |
| | | | | Dgkh |
| | | | | Dgkq |
| | | | | Dhx9 |
| | | | | Diaph1 |
| | | | | Diaph2 |
| | | | | Diaph3 |
| | | | | Dido1 |
| | | | | Dis3L |
| | | | | Disc1 |
| | | | | Dixdc1 |
| | | | | Dlc1 |
| | | | | Dlg1 |
| | | | | Dlg4 |
| | | | | Dlg5 |
| | | | | Dlgap5 |
| | | | | Dmd |

| | | | | |
|---|---|---|---|---|
| | | | | Dmtn |
| | | | | Dnaaf1 |
| | | | | Dnaaf2 |
| | | | | Dnaaf4 |
| | | | | Dnaaf5 |
| | | | | Dnah1 |
| | | | | Dnah10 |
| | | | | Dnah11 |
| | | | | Dnah12 |
| | | | | Dnah14 |
| | | | | Dnah17 |
| | | | | Dnah2 |
| | | | | Dnah3 |
| | | | | Dnah5 |
| | | | | Dnah6 |
| | | | | Dnah7A |
| | | | | Dnah7B |
| | | | | Dnah7C |
| | | | | Dnah8 |
| | | | | Dnah9 |
| | | | | Dnai1 |
| | | | | Dnai2 |
| | | | | Dnai3 |
| | | | | Dnai4 |
| | | | | Dnai7 |
| | | | | Dnaja1 |
| | | | | Dnaja3 |
| | | | | Dnajb13 |
| | | | | Dnajb3 |
| | | | | Dnajc24 |
| | | | | Dnajc7 |
| | | | | Dnal1 |
| | | | | Dnal4 |
| | | | | Dnali1 |
| | | | | Dnhd1 |
| | | | | Dnm1 |
| | | | | Dnm1L |
| | | | | Dnm2 |
| | | | | Dnm3 |
| | | | | Dnmbp |
| | | | | Dock2 |
| | | | | Dock5 |
| | | | | Dpf2 |
| | | | | Dpp9 |
| | | | | Dpysl2 |
| | | | | Dpysl3 |
| | | | | Dr1 |

| | | | | |
|---|---|---|---|---|
| | | | | Drc1 |
| | | | | Drc3 |
| | | | | Drc7 |
| | | | | Drd4 |
| | | | | Dsn1 |
| | | | | Dsp |
| | | | | Dst |
| | | | | Dstn |
| | | | | Dtl |
| | | | | Dtna |
| | | | | Dtnbp1 |
| | | | | Dtx4 |
| | | | | Dusp21 |
| | | | | Dusp22 |
| | | | | Dusp3 |
| | | | | Dvl1 |
| | | | | Dvl2 |
| | | | | Dydc1 |
| | | | | Dync1H1 |
| | | | | Dync1I1 |
| | | | | Dync1I2 |
| | | | | Dync1Li1 |
| | | | | Dync1Li2 |
| | | | | Dync2H1 |
| | | | | Dync2I1 |
| | | | | Dync2I2 |
| | | | | Dync2Li1 |
| | | | | Dynll1 |
| | | | | Dynll2 |
| | | | | Dynlrb1 |
| | | | | Dynlrb2 |
| | | | | Dynlt1A |
| | | | | Dynlt1B |
| | | | | Dynlt1C |
| | | | | Dynlt1F |
| | | | | Dynlt2A1 |
| | | | | Dynlt2B |
| | | | | Dynlt3 |
| | | | | Dynlt4 |
| | | | | Dynlt5 |
| | | | | Dyrk1A |
| | | | | Dyrk2 |
| | | | | Dyrk3 |
| | | | | Dyrk4 |
| | | | | Dysf |
| | | | | Dzank1 |
| | | | | Dzip1 |

| | | | | |
|---|---|---|---|---|
| | | | | Dzip1L |
| | | | | E2F1 |
| | | | | E4F1 |
| | | | | Ecpas |
| | | | | Ect2 |
| | | | | Eef1A1 |
| | | | | Eef1Akmt3 |
| | | | | Efcab2 |
| | | | | Efcab6 |
| | | | | Efhb |
| | | | | Efhc1 |
| | | | | Efhc2 |
| | | | | Efr3B |
| | | | | Egfr |
| | | | | Ehd2 |
| | | | | Eif3A |
| | | | | Eif6 |
| | | | | Elmod3 |
| | | | | Emd |
| | | | | Eml1 |
| | | | | Eml2 |
| | | | | Eml3 |
| | | | | Eml4 |
| | | | | Eml5 |
| | | | | Eml6 |
| | | | | Enah |
| | | | | Enc1 |
| | | | | Enkd1 |
| | | | | Enkur |
| | | | | Entr1 |
| | | | | Epb41 |
| | | | | Epb41L1 |
| | | | | Epb41L2 |
| | | | | Epb41L3 |
| | | | | Epb41L4A |
| | | | | Epb41L4B |
| | | | | Epb41L5 |
| | | | | Epb42 |
| | | | | Epha3 |
| | | | | Eppk1 |
| | | | | Eps8L2 |
| | | | | Erc1 |
| | | | | Erc2 |
| | | | | Ercc2 |
| | | | | Ercc6L2 |
| | | | | Ermn |
| | | | | Esam |

| | | | | |
|---|---|---|---|---|
| | | | | Espl1 |
| | | | | Espn |
| | | | | Esrra |
| | | | | Etl4 |
| | | | | Evc |
| | | | | Evc2 |
| | | | | Evi5 |
| | | | | Evl |
| | | | | Evpl |
| | | | | Exd2 |
| | | | | Exoc4 |
| | | | | Exoc7 |
| | | | | Eya3 |
| | | | | Ezr |
| | | | | Fam107A |
| | | | | Fam110A |
| | | | | Fam110B |
| | | | | Fam110C |
| | | | | Fam161A |
| | | | | Fam161B |
| | | | | Fam184A |
| | | | | Fam234B |
| | | | | Fam83D |
| | | | | Fam83H |
| | | | | Fance |
| | | | | Fank1 |
| | | | | Farp1 |
| | | | | Farp2 |
| | | | | Fbf1 |
| | | | | Fblim1 |
| | | | | Fbxl13 |
| | | | | Fbxl7 |
| | | | | Fbxo31 |
| | | | | Fbxo5 |
| | | | | Fbxw11 |
| | | | | Fbxw8 |
| | | | | Fcgr2B |
| | | | | Fchsd1 |
| | | | | Fer |
| | | | | Fermt1 |
| | | | | Fermt2 |
| | | | | Fermt3 |
| | | | | Fes |
| | | | | Fez1 |
| | | | | Ffar4 |
| | | | | Fgd1 |
| | | | | Fgd2 |

| | | | | |
|---|---|---|---|---|
| | | | | Fgd3 |
| | | | | Fgd4 |
| | | | | Fgd5 |
| | | | | Fgd6 |
| | | | | Fgf13 |
| | | | | Fgr |
| | | | | Fhdc1 |
| | | | | Fhl3 |
| | | | | Fhod1 |
| | | | | Fhod3 |
| | | | | Fign |
| | | | | Filip1 |
| | | | | Filip1L |
| | | | | Firrm |
| | | | | Fkbp15 |
| | | | | Fkbp4 |
| | | | | Fkbp6 |
| | | | | Flacc1 |
| | | | | Flcn |
| | | | | Flii |
| | | | | Flna |
| | | | | Flnb |
| | | | | Flnc |
| | | | | Flot1 |
| | | | | Flot2 |
| | | | | Flt1 |
| | | | | Fmn1 |
| | | | | Fmn2 |
| | | | | Fmr1 |
| | | | | Fnbp1 |
| | | | | Fnbp1L |
| | | | | Fnip2 |
| | | | | Fnta |
| | | | | Fntb |
| | | | | Foxa3 |
| | | | | Frmd3 |
| | | | | Frmd4A |
| | | | | Frmd4B |
| | | | | Frmd5 |
| | | | | Frmd6 |
| | | | | Frmd7 |
| | | | | Frmd8 |
| | | | | Frmpd1 |
| | | | | Frmpd4 |
| | | | | Fry |
| | | | | Fscn1 |
| | | | | Fscn2 |

| | | | | |
|---|---|---|---|---|
| | | | | Fscn3 |
| | | | | Fsd1 |
| | | | | Ftcd |
| | | | | Fuz |
| | | | | Fyn |
| | | | | Fzd6 |
| | | | | G6Pd2 |
| | | | | G6Pdx |
| | | | | Gabarap |
| | | | | Gabarapl1 |
| | | | | Gabrg3 |
| | | | | Gan |
| | | | | Gapdh |
| | | | | Gapdhrt |
| | | | | Gapdhrt2 |
| | | | | Gas2 |
| | | | | Gas2L1 |
| | | | | Gas2L2 |
| | | | | Gas2L3 |
| | | | | Gas7 |
| | | | | Gas8 |
| | | | | Gbp2 |
| | | | | Gbp2B |
| | | | | Gck |
| | | | | Gdpd2 |
| | | | | Gem |
| | | | | Gen1 |
| | | | | Gfap |
| | | | | Gfral |
| | | | | Git1 |
| | | | | Gja1 |
| | | | | Gja2 |
| | | | | Gja3 |
| | | | | Gja4 |
| | | | | Gja5 |
| | | | | Gjb1 |
| | | | | Gjb2 |
| | | | | Gjb3 |
| | | | | Gjb4 |
| | | | | Gjb5 |
| | | | | Gjb6 |
| | | | | Gjb7 |
| | | | | Gjc1 |
| | | | | Gjc2 |
| | | | | Gjc3 |
| | | | | Gjd2 |
| | | | | Gjd3 |

| | | | | |
|---|---|---|---|---|
| | | | | Gjd4 |
| | | | | Gle1 |
| | | | | Glg1 |
| | | | | Gli1 |
| | | | | Gli2 |
| | | | | Gli3 |
| | | | | Gm14137 |
| | | | | Gm5414 |
| | | | | Gm5478 |
| | | | | Gmfb |
| | | | | Gmfg |
| | | | | Gnai1 |
| | | | | Gnai2 |
| | | | | Gnai3 |
| | | | | Gnat3 |
| | | | | Gng12 |
| | | | | Golga2 |
| | | | | Gpaa1 |
| | | | | Gper1 |
| | | | | Gphn |
| | | | | Gpr174 |
| | | | | Gpsm2 |
| | | | | Gpx2 |
| | | | | Gramd2B |
| | | | | Grb2 |
| | | | | Grhl3 |
| | | | | Grin2B |
| | | | | Grip1 |
| | | | | Gsk3A |
| | | | | Gsk3B |
| | | | | Gsn |
| | | | | Gtf2F2 |
| | | | | Gtse1 |
| | | | | Gypc |
| | | | | Gys2 |
| | | | | H1F0 |
| | | | | H2Ax |
| | | | | Hap1 |
| | | | | Harbi1 |
| | | | | Haspin |
| | | | | Haus1 |
| | | | | Haus2 |
| | | | | Haus3 |
| | | | | Haus4 |
| | | | | Haus5 |
| | | | | Haus6 |
| | | | | Haus7 |

| | | | | |
|---|---|---|---|---|
| | | | | Haus8 |
| | | | | Hax1 |
| | | | | Hck |
| | | | | Hcls1 |
| | | | | Hdac3 |
| | | | | Hdac4 |
| | | | | Hdac6 |
| | | | | Hecw2 |
| | | | | Hepacam2 |
| | | | | Herc2 |
| | | | | Hfe |
| | | | | Hid1 |
| | | | | Hip1 |
| | | | | Hip1R |
| | | | | Hipk1 |
| | | | | Hk2 |
| | | | | Hmbox1 |
| | | | | Hmmr |
| | | | | Hnf4G |
| | | | | Hnmt |
| | | | | Hnrnpc |
| | | | | Hnrnpk |
| | | | | Hnrnpu |
| | | | | Hook1 |
| | | | | Hook2 |
| | | | | Hook3 |
| | | | | Hormad2 |
| | | | | Hoxa13 |
| | | | | Hoxb4 |
| | | | | Hoxc8 |
| | | | | Hras |
| | | | | Hsdl1 |
| | | | | Hsf1 |
| | | | | Hspa1A |
| | | | | Hspa1B |
| | | | | Hspa2 |
| | | | | Hspa8 |
| | | | | Hspb1 |
| | | | | Hspb7 |
| | | | | Hsph1 |
| | | | | Htr2A |
| | | | | Htra2 |
| | | | | Htt |
| | | | | Hydin |
| | | | | Hyls1 |
| | | | | Hypk |
| | | | | Id1 |

| | | | | |
|---|---|---|---|---|
| | | | | Iffo1 |
| | | | | Iffo2 |
| | | | | Ift122 |
| | | | | Ift140 |
| | | | | Ift172 |
| | | | | Ift20 |
| | | | | Ift22 |
| | | | | Ift25 |
| | | | | Ift27 |
| | | | | Ift43 |
| | | | | Ift46 |
| | | | | Ift52 |
| | | | | Ift56 |
| | | | | Ift57 |
| | | | | Ift70A1 |
| | | | | Ift70A2 |
| | | | | Ift70B |
| | | | | Ift74 |
| | | | | Ift80 |
| | | | | Ift81 |
| | | | | Ift88 |
| | | | | Igbp1 |
| | | | | Igf2Bp2 |
| | | | | Ik |
| | | | | Ikbkg |
| | | | | Il1Rn |
| | | | | Il4Ra |
| | | | | Ildr1 |
| | | | | Ildr2 |
| | | | | Ilk |
| | | | | Ilrun |
| | | | | Ina |
| | | | | Incenp |
| | | | | Ing4 |
| | | | | Ino80 |
| | | | | Inpp5D |
| | | | | Inpp5E |
| | | | | Inppl1 |
| | | | | Ints6 |
| | | | | Intu |
| | | | | Invs |
| | | | | Ipp |
| | | | | Iqca1 |
| | | | | Iqcb1 |
| | | | | Iqcd |
| | | | | Iqcg |
| | | | | Iqgap1 |

| | | | | |
|---|---|---|---|---|
| | | | | Iqgap2 |
| | | | | Iqub |
| | | | | Irag2 |
| | | | | Irs1 |
| | | | | Ist1 |
| | | | | Itgb1Bp1 |
| | | | | Itgb6 |
| | | | | Itpka |
| | | | | Itpkb |
| | | | | Itsn1 |
| | | | | Itsn2 |
| | | | | Ivl |
| | | | | Ivns1Abp |
| | | | | Jade1 |
| | | | | Jak1 |
| | | | | Jak2 |
| | | | | Jak3 |
| | | | | Jakmip1 |
| | | | | Jam3 |
| | | | | Jam-4 |
| | | | | Jam-A |
| | | | | Jam-B |
| | | | | Jam-C |
| | | | | Jam-L |
| | | | | Jmy |
| | | | | Jpt1 |
| | | | | Jtb |
| | | | | Jup |
| | | | | Kalrn |
| | | | | Kank1 |
| | | | | Kank3 |
| | | | | Kank4 |
| | | | | Kansl2 |
| | | | | Kash5 |
| | | | | Kat14 |
| | | | | Kat2A |
| | | | | Kat2B |
| | | | | Kat5 |
| | | | | Katna1 |
| | | | | Katnal1 |
| | | | | Katnal2 |
| | | | | Katnb1 |
| | | | | Katnbl1 |
| | | | | Katnip |
| | | | | Kazn |
| | | | | Kbtbd8 |
| | | | | Kcnab2 |

| | | | | |
|---|---|---|---|---|
| | | | | Kcnc3 |
| | | | | Keap1 |
| | | | | Keg1 |
| | | | | Khdc3 |
| | | | | Kif11 |
| | | | | Kif12 |
| | | | | Kif13A |
| | | | | Kif13B |
| | | | | Kif14 |
| | | | | Kif15 |
| | | | | Kif16B |
| | | | | Kif17 |
| | | | | Kif18A |
| | | | | Kif18B |
| | | | | Kif19A |
| | | | | Kif19B |
| | | | | Kif1A |
| | | | | Kif1B |
| | | | | Kif1C |
| | | | | Kif20A |
| | | | | Kif20B |
| | | | | Kif21A |
| | | | | Kif21B |
| | | | | Kif22 |
| | | | | Kif23 |
| | | | | Kif24 |
| | | | | Kif26A |
| | | | | Kif26B |
| | | | | Kif27 |
| | | | | Kif28 |
| | | | | Kif2A |
| | | | | Kif2B |
| | | | | Kif2C |
| | | | | Kif3A |
| | | | | Kif3B |
| | | | | Kif3C |
| | | | | Kif4 |
| | | | | Kif5A |
| | | | | Kif5B |
| | | | | Kif5C |
| | | | | Kif6 |
| | | | | Kif7 |
| | | | | Kif9 |
| | | | | Kifap3 |
| | | | | Kifbp |
| | | | | Kifc1 |
| | | | | Kifc2 |

| | | | | |
|---|---|---|---|---|
| | | | | Kifc3 |
| | | | | Kifc5B |
| | | | | Kitl |
| | | | | Kiz |
| | | | | Klc1 |
| | | | | Klc2 |
| | | | | Klc3 |
| | | | | Klc4 |
| | | | | Klf4 |
| | | | | Klhl1 |
| | | | | Klhl12 |
| | | | | Klhl14 |
| | | | | Klhl17 |
| | | | | Klhl2 |
| | | | | Klhl21 |
| | | | | Klhl22 |
| | | | | Klhl3 |
| | | | | Klhl4 |
| | | | | Klhl41 |
| | | | | Klhl42 |
| | | | | Kmt2E |
| | | | | Kmt5B |
| | | | | Kncn |
| | | | | Knstrn |
| | | | | Kntc1 |
| | | | | Kpna7 |
| | | | | Kptn |
| | | | | Krit1 |
| | | | | Krt1 |
| | | | | Krt10 |
| | | | | Krt12 |
| | | | | Krt13 |
| | | | | Krt14 |
| | | | | Krt15 |
| | | | | Krt16 |
| | | | | Krt17 |
| | | | | Krt18 |
| | | | | Krt19 |
| | | | | Krt2 |
| | | | | Krt20 |
| | | | | Krt222 |
| | | | | Krt23 |
| | | | | Krt24 |
| | | | | Krt25 |
| | | | | Krt26 |
| | | | | Krt27 |
| | | | | Krt28 |

| | | | | |
|---|---|---|---|---|
| | | | | Krt31 |
| | | | | Krt32 |
| | | | | Krt33A |
| | | | | Krt33B |
| | | | | Krt34 |
| | | | | Krt35 |
| | | | | Krt36 |
| | | | | Krt39 |
| | | | | Krt4 |
| | | | | Krt40 |
| | | | | Krt42 |
| | | | | Krt5 |
| | | | | Krt6A |
| | | | | Krt6B |
| | | | | Krt7 |
| | | | | Krt71 |
| | | | | Krt72 |
| | | | | Krt73 |
| | | | | Krt74 |
| | | | | Krt75 |
| | | | | Krt76 |
| | | | | Krt77 |
| | | | | Krt78 |
| | | | | Krt79 |
| | | | | Krt8 |
| | | | | Krt80 |
| | | | | Krt81 |
| | | | | Krt82 |
| | | | | Krt83 |
| | | | | Krt84 |
| | | | | Krt85 |
| | | | | Krt86 |
| | | | | Krt87 |
| | | | | Krt9 |
| | | | | Krt90 |
| | | | | Krtap12-1 |
| | | | | Krtap14 |
| | | | | Krtap15-1 |
| | | | | Krtap16-1 |
| | | | | Krtap16-3 |
| | | | | Krtap19-1 |
| | | | | Krtap19-2 |
| | | | | Krtap19-3 |
| | | | | Krtap19-4 |
| | | | | Krtap19-5 |
| | | | | Krtap19-9B |
| | | | | Krtap21-1 |

| | | | | |
|---|---|---|---|---|
| | | | | Krtap26-1 |
| | | | | Krtap29-1 |
| | | | | Krtap3-1 |
| | | | | Krtap3-2 |
| | | | | Krtap3-3 |
| | | | | Krtap4-6 |
| | | | | Krtap5-1 |
| | | | | Krtap5-2 |
| | | | | Krtap5-3 |
| | | | | Krtap5-4 |
| | | | | Krtap5-5 |
| | | | | Krtap6-2 |
| | | | | Krtap6-5 |
| | | | | Krtap7-1 |
| | | | | Krtap8-1 |
| | | | | Krtap9-3 |
| | | | | Ky |
| | | | | Lad1 |
| | | | | Lancl2 |
| | | | | Lasp1 |
| | | | | Lats1 |
| | | | | Lats2 |
| | | | | Lca5 |
| | | | | Lca5L |
| | | | | Lck |
| | | | | Lcp1 |
| | | | | Ldb3 |
| | | | | Ldlrap1 |
| | | | | Lemd2 |
| | | | | Leo1 |
| | | | | Lhcgr |
| | | | | Lima1 |
| | | | | Limch1 |
| | | | | Limd2 |
| | | | | Limk1 |
| | | | | Limk2 |
| | | | | Llgl1 |
| | | | | Llgl2 |
| | | | | Lmna |
| | | | | Lmnb1 |
| | | | | Lmnb2 |
| | | | | Lmntd1 |
| | | | | Lmntd2 |
| | | | | Lmod1 |
| | | | | Lmod2 |
| | | | | Lmod3 |
| | | | | Lpp |

| | | | | |
|---|---|---|---|---|
| | | | | Lpxn |
| | | | | Lrguk |
| | | | | Lrif1 |
| | | | | Lrp1 |
| | | | | Lrp8 |
| | | | | Lrpprc |
| | | | | Lrrc10 |
| | | | | Lrrc23 |
| | | | | Lrrc25 |
| | | | | Lrrc26 |
| | | | | Lrrc45 |
| | | | | Lrrc49 |
| | | | | Lrrc51 |
| | | | | Lrrc7 |
| | | | | Lrrcc1 |
| | | | | Lrriq1 |
| | | | | Lrrk2 |
| | | | | Lrwd1 |
| | | | | Lsm14A |
| | | | | Lsr |
| | | | | Lurap1 |
| | | | | Luzp1 |
| | | | | Lyst |
| | | | | Lztfl1 |
| | | | | Lzts2 |
| | | | | Lzts3 |
| | | | | Macf1 |
| | | | | Macroh2A1 |
| | | | | Mad1L1 |
| | | | | Mad2L1 |
| | | | | Mad2L1Bp |
| | | | | Mad2L2 |
| | | | | Maea |
| | | | | Magi-1 |
| | | | | Magi2 |
| | | | | Mak |
| | | | | Mamld1 |
| | | | | Map10 |
| | | | | Map1A |
| | | | | Map1B |
| | | | | Map1Lc3A |
| | | | | Map1Lc3B |
| | | | | Map1S |
| | | | | Map2 |
| | | | | Map2K1 |
| | | | | Map2K2 |
| | | | | Map2K5 |

| | | | | |
|---|---|---|---|---|
| | | | | Map2K6 |
| | | | | Map3K1 |
| | | | | Map3K11 |
| | | | | Map4 |
| | | | | Map6 |
| | | | | Map6D1 |
| | | | | Map7 |
| | | | | Map7D1 |
| | | | | Map7D2 |
| | | | | Map7D3 |
| | | | | Map9 |
| | | | | Mapk1 |
| | | | | Mapk14 |
| | | | | Mapk15 |
| | | | | Mapk1Ip1 |
| | | | | Mapk3 |
| | | | | Mapk6 |
| | | | | Mapk8 |
| | | | | Mapkapk2 |
| | | | | Mapkapk5 |
| | | | | Mapkbp1 |
| | | | | Mapre1 |
| | | | | Mapre2 |
| | | | | Mapre3 |
| | | | | Mapt |
| | | | | Marchf7 |
| | | | | Marcks |
| | | | | Marcksl1 |
| | | | | Mark1 |
| | | | | Mark2 |
| | | | | Mark4 |
| | | | | Marveld1 |
| | | | | Marveld2 |
| | | | | Marveld3 |
| | | | | Mast1 |
| | | | | Mast2 |
| | | | | Mastl |
| | | | | Matcap1 |
| | | | | Mbip |
| | | | | Mcm3 |
| | | | | Mcph1 |
| | | | | Mdh1 |
| | | | | Mdm1 |
| | | | | Mdm2 |
| | | | | Mdn1 |
| | | | | Mecp2 |
| | | | | Med28 |

| | | | | |
|---|---|---|---|---|
| | | | | Mefv |
| | | | | Meig1 |
| | | | | Mfap1A |
| | | | | Mfap1B |
| | | | | Mfn2 |
| | | | | Mib1 |
| | | | | Mical1 |
| | | | | Mical2 |
| | | | | Mical3 |
| | | | | Micall1 |
| | | | | Micall2 |
| | | | | Mid1 |
| | | | | Mid1Ip1 |
| | | | | Mid2 |
| | | | | Mis12 |
| | | | | Misp |
| | | | | Mkks |
| | | | | Mks1 |
| | | | | Mlf1 |
| | | | | Mllt11 |
| | | | | Mlph |
| | | | | Mme |
| | | | | Mmp14 |
| | | | | Mms19 |
| | | | | Mns1 |
| | | | | Mphosph9 |
| | | | | Mplkip |
| | | | | Mplkipl1 |
| | | | | Mpp1 |
| | | | | Mpp2 |
| | | | | Mpp4 |
| | | | | Mprip |
| | | | | Msn |
| | | | | Msra |
| | | | | Msrb1 |
| | | | | Mst1R |
| | | | | Mt3 |
| | | | | Mta1 |
| | | | | Mtcl1 |
| | | | | Mtcl2 |
| | | | | Mtpn |
| | | | | Mtrr |
| | | | | Mtss1 |
| | | | | Mtss2 |
| | | | | Mtus1 |
| | | | | Mtus2 |
| | | | | Mupp1 |

| | | | | |
|---|---|---|---|---|
| | | | | Mvb12A |
| | | | | Mvp |
| | | | | Mx2 |
| | | | | Myadm |
| | | | | Mybpc1 |
| | | | | Mybpc2 |
| | | | | Mybpc3 |
| | | | | Myc |
| | | | | Mycbp2 |
| | | | | Myf6 |
| | | | | Myh1 |
| | | | | Myh10 |
| | | | | Myh11 |
| | | | | Myh13 |
| | | | | Myh14 |
| | | | | Myh15 |
| | | | | Myh2 |
| | | | | Myh3 |
| | | | | Myh4 |
| | | | | Myh6 |
| | | | | Myh7 |
| | | | | Myh7B |
| | | | | Myh8 |
| | | | | Myh9 |
| | | | | Myl1 |
| | | | | Myl12A |
| | | | | Myl12B |
| | | | | Myl2 |
| | | | | Myl3 |
| | | | | Myl4 |
| | | | | Myl6 |
| | | | | Myl6B |
| | | | | Myl7 |
| | | | | Myl9 |
| | | | | Mylip |
| | | | | Mylk |
| | | | | Mylk3 |
| | | | | Mylpf |
| | | | | Myo10 |
| | | | | Myo15A |
| | | | | Myo16 |
| | | | | Myo18A |
| | | | | Myo18B |
| | | | | Myo19 |
| | | | | Myo1A |
| | | | | Myo1B |
| | | | | Myo1C |

| | | | | |
|---|---|---|---|---|
| | | | | Myo1D |
| | | | | Myo1E |
| | | | | Myo1F |
| | | | | Myo1G |
| | | | | Myo1H |
| | | | | Myo3A |
| | | | | Myo3B |
| | | | | Myo5A |
| | | | | Myo5B |
| | | | | Myo5C |
| | | | | Myo6 |
| | | | | Myo7A |
| | | | | Myo7B |
| | | | | Myo9A |
| | | | | Myo9B |
| | | | | Myof |
| | | | | Myom1 |
| | | | | Myom2 |
| | | | | Myot |
| | | | | Myoz1 |
| | | | | Myoz2 |
| | | | | Myoz3 |
| | | | | Myrip |
| | | | | Myzap |
| | | | | Mzt1 |
| | | | | Mzt2 |
| | | | | Naa11 |
| | | | | Naa12 |
| | | | | Naa40 |
| | | | | Narf |
| | | | | Nav1 |
| | | | | Nav3 |
| | | | | Nckap1 |
| | | | | Nckap5 |
| | | | | Nckap5L |
| | | | | Ncoa5 |
| | | | | Ncor1 |
| | | | | Ndc1 |
| | | | | Ndc80 |
| | | | | Nde1 |
| | | | | Ndel1 |
| | | | | Ndn |
| | | | | Ndor1 |
| | | | | Ndrg1 |
| | | | | Neb |
| | | | | Nebl |
| | | | | Nedd1 |

| | | | | |
|---|---|---|---|---|
| | | | | Nedd9 |
| | | | | Nefh |
| | | | | Nefl |
| | | | | Nefm |
| | | | | Neil1 |
| | | | | Neil2 |
| | | | | Nek1 |
| | | | | Nek2 |
| | | | | Nek4 |
| | | | | Nek6 |
| | | | | Nek7 |
| | | | | Nek8 |
| | | | | Nek9 |
| | | | | Nes |
| | | | | Neurl1B |
| | | | | Neurl4 |
| | | | | Nexn |
| | | | | Nf2 |
| | | | | Nfe2L2 |
| | | | | Nfs1 |
| | | | | Ngrn |
| | | | | Nicn1 |
| | | | | Nin |
| | | | | Ninl |
| | | | | Nisch |
| | | | | Nit2 |
| | | | | Nlrc3 |
| | | | | Nlrc5 |
| | | | | Nlrp3 |
| | | | | Nme1 |
| | | | | Nme2 |
| | | | | Nme3 |
| | | | | Nme5 |
| | | | | Nme7 |
| | | | | Nme8 |
| | | | | Nme9 |
| | | | | Nod2 |
| | | | | Nol9 |
| | | | | Nos1 |
| | | | | Nos1Ap |
| | | | | Nos2 |
| | | | | Nos3 |
| | | | | Nostrin |
| | | | | Notch1 |
| | | | | Nox4 |
| | | | | Npffr2 |
| | | | | Nphp1 |

| | | | | |
|---|---|---|---|---|
| | | | | Nphp4 |
| | | | | Npm1 |
| | | | | Npm3 |
| | | | | Nr0B1 |
| | | | | Nr1I2 |
| | | | | Nr1I3 |
| | | | | Nr3C1 |
| | | | | Nrp1 |
| | | | | Nsfl1C |
| | | | | Nsl1 |
| | | | | Nsmce1 |
| | | | | Nsmf |
| | | | | Nsun2 |
| | | | | Ntn1 |
| | | | | Nuak1 |
| | | | | Nubp1 |
| | | | | Nubp2 |
| | | | | Nudc |
| | | | | Nudcd2 |
| | | | | Nudcd3 |
| | | | | Nudt21 |
| | | | | Numa1 |
| | | | | Nup62 |
| | | | | Nup85 |
| | | | | Nup93 |
| | | | | Nusap1 |
| | | | | Obsl1 |
| | | | | Ocm |
| | | | | Ocrl |
| | | | | Odad1 |
| | | | | Odad2 |
| | | | | Odad3 |
| | | | | Odad4 |
| | | | | Odam |
| | | | | Odf1 |
| | | | | Odf2 |
| | | | | Odf2L |
| | | | | Odf4 |
| | | | | Ofcc1 |
| | | | | Ofd1 |
| | | | | Ola1 |
| | | | | Onecut2 |
| | | | | Opa1 |
| | | | | Ophn1 |
| | | | | Or2A7 |
| | | | | Orc2 |
| | | | | Osbpl10 |

| | | | | |
|---|---|---|---|---|
| | | | | Ovgp1 |
| | | | | P2Rx7 |
| | | | | P4Hb |
| | | | | Pacrg |
| | | | | Pacsin2 |
| | | | | Padi6 |
| | | | | Pafah1B1 |
| | | | | Pak1 |
| | | | | Palld |
| | | | | Pals1 |
| | | | | Par3 |
| | | | | Par6 |
| | | | | Pard3 |
| | | | | Pard6A |
| | | | | Parp3 |
| | | | | Parp4 |
| | | | | Parva |
| | | | | Parvb |
| | | | | Parvg |
| | | | | Patj |
| | | | | Pawr |
| | | | | Pax2 |
| | | | | Pbxip1 |
| | | | | Pcgf5 |
| | | | | Pcif1 |
| | | | | Pclaf |
| | | | | Pclo |
| | | | | Pcm1 |
| | | | | Pcna |
| | | | | Pcnt |
| | | | | Pcp4 |
| | | | | Pdcd6Ip |
| | | | | Pde4B |
| | | | | Pde4D |
| | | | | Pde4Dip |
| | | | | Pde6D |
| | | | | Pdlim1 |
| | | | | Pdlim2 |
| | | | | Pdlim3 |
| | | | | Pdlim4 |
| | | | | Pdlim5 |
| | | | | Pdlim7 |
| | | | | Pdxp |
| | | | | Pdzd2 |
| | | | | Pdzd7 |
| | | | | Pea15A |
| | | | | Peak1 |

| | | | | |
|---|---|---|---|---|
| | | | | Pfdn1 |
| | | | | Pfdn5 |
| | | | | Pfn1 |
| | | | | Pfn2 |
| | | | | Pfn3 |
| | | | | Pgm5 |
| | | | | Phf1 |
| | | | | Phldb2 |
| | | | | Phlpp2 |
| | | | | Pias1 |
| | | | | Pibf1 |
| | | | | Pick1 |
| | | | | Pierce1 |
| | | | | Pierce2 |
| | | | | Piezo1 |
| | | | | Piezo2 |
| | | | | Pik3R4 |
| | | | | Pik3R5 |
| | | | | Pin4 |
| | | | | Pink1 |
| | | | | Pinx1 |
| | | | | Pitpnm2 |
| | | | | Pja2 |
| | | | | Pjvk |
| | | | | Pkd2 |
| | | | | Pkd2L1 |
| | | | | Pkhd1 |
| | | | | Pkn2 |
| | | | | Pknox2 |
| | | | | Pkp2 |
| | | | | Pkp4 |
| | | | | Pla2G3 |
| | | | | Pla2G4C |
| | | | | Pla2G6 |
| | | | | Plag1 |
| | | | | Plec |
| | | | | Plek2 |
| | | | | Plekha7 |
| | | | | Plekhg3 |
| | | | | Plekhg6 |
| | | | | Plekhh1 |
| | | | | Plekhh2 |
| | | | | Plekhh3 |
| | | | | Plekhn1 |
| | | | | Plk1 |
| | | | | Plk2 |
| | | | | Plk3 |

| | | | | |
|---|---|---|---|---|
| | | | | Plk4 |
| | | | | Plk5 |
| | | | | Pls1 |
| | | | | Pls3 |
| | | | | Pmf1 |
| | | | | Pmm2 |
| | | | | Pnma5 |
| | | | | Poc1A |
| | | | | Poc1B |
| | | | | Poc5 |
| | | | | Podxl |
| | | | | Pof1B |
| | | | | Pola2 |
| | | | | Polb |
| | | | | Poldip2 |
| | | | | Polr3H |
| | | | | Popdc2 |
| | | | | Popdc3 |
| | | | | Ppl |
| | | | | Ppp1Cc |
| | | | | Ppp1R12A |
| | | | | Ppp1R12B |
| | | | | Ppp1R12C |
| | | | | Ppp1R18 |
| | | | | Ppp1R35 |
| | | | | Ppp1R42 |
| | | | | Ppp1R9A |
| | | | | Ppp1R9B |
| | | | | Ppp2Ca |
| | | | | Ppp2Cb |
| | | | | Ppp2R2B |
| | | | | Ppp2R3C |
| | | | | Ppp2R5A |
| | | | | Ppp4C |
| | | | | Ppp4R2 |
| | | | | Ppp4R3A |
| | | | | Ppp4R3B |
| | | | | Ppp4R4 |
| | | | | Pqbp1 |
| | | | | Prc1 |
| | | | | Prepl |
| | | | | Prickle4 |
| | | | | Prkaa1 |
| | | | | Prkaa2 |
| | | | | Prkaca |
| | | | | Prkacb |
| | | | | Prkar1A |

| | | | | |
|---|---|---|---|---|
| | | | | Prkar2A |
| | | | | Prkar2B |
| | | | | Prkca |
| | | | | Prkcb |
| | | | | Prkce |
| | | | | Prkci |
| | | | | Prkcq |
| | | | | Prkcz |
| | | | | Procr |
| | | | | Proser3 |
| | | | | Prpf19 |
| | | | | Prpf6 |
| | | | | Prph |
| | | | | Psen1 |
| | | | | Psen2 |
| | | | | Pskh1 |
| | | | | Psma1 |
| | | | | Psmb4 |
| | | | | Psmb5 |
| | | | | Psmc4 |
| | | | | Psmd10 |
| | | | | Psme3 |
| | | | | Psrc1 |
| | | | | Pstpip1 |
| | | | | Pstpip2 |
| | | | | Ptges3 |
| | | | | Ptk2 |
| | | | | Ptk2B |
| | | | | Ptp4A1 |
| | | | | Ptpn11 |
| | | | | Ptpn12 |
| | | | | Ptpn13 |
| | | | | Ptpn14 |
| | | | | Ptpn20 |
| | | | | Ptpn21 |
| | | | | Ptpn23 |
| | | | | Ptpn3 |
| | | | | Ptpn4 |
| | | | | Ptpn7 |
| | | | | Pttg1 |
| | | | | Pvalb |
| | | | | Pxk |
| | | | | Pxn |
| | | | | Pycard |
| | | | | Pycr3 |
| | | | | Rab10 |
| | | | | Rab11A |

| | | | | |
|---|---|---|---|---|
| | | | | Rab11Fip3 |
| | | | | Rab11Fip5 |
| | | | | Rab22A |
| | | | | Rab23 |
| | | | | Rab28 |
| | | | | Rab29 |
| | | | | Rab34 |
| | | | | Rab3D |
| | | | | Rab3Ip |
| | | | | Rab5A |
| | | | | Rab6A |
| | | | | Rab6B |
| | | | | Rab8A |
| | | | | Rabep2 |
| | | | | Rabgap1 |
| | | | | Rabl2 |
| | | | | Rabl6 |
| | | | | Rac1 |
| | | | | Rac2 |
| | | | | Rac3 |
| | | | | Racgap1 |
| | | | | Rad18 |
| | | | | Rad51 |
| | | | | Rad51D |
| | | | | Radil |
| | | | | Rae1 |
| | | | | Rai14 |
| | | | | Ralbp1 |
| | | | | Ran |
| | | | | Ranbp1 |
| | | | | Ranbp10 |
| | | | | Ranbp9 |
| | | | | Rangap1 |
| | | | | Rap1Gap2 |
| | | | | Rapgef3 |
| | | | | Rapgef6 |
| | | | | Rapsn |
| | | | | Rara |
| | | | | Rassf1 |
| | | | | Rassf10 |
| | | | | Rassf3 |
| | | | | Rassf5 |
| | | | | Rassf7 |
| | | | | Rb1 |
| | | | | Rbbp6 |
| | | | | Rbm39 |
| | | | | Rcc2 |

| | | | | |
|---|---|---|---|---|
| | | | | Rdx |
| | | | | Reep1 |
| | | | | Reep2 |
| | | | | Reep3 |
| | | | | Reep4 |
| | | | | Relb |
| | | | | Rell1 |
| | | | | Rflna |
| | | | | Rflnb |
| | | | | Rgcc |
| | | | | Rgs14 |
| | | | | Rgs22 |
| | | | | Rhbg |
| | | | | Rhoa |
| | | | | Rhobtb1 |
| | | | | Rhobtb2 |
| | | | | Rhof |
| | | | | Rhog |
| | | | | Rhoq |
| | | | | Rhou |
| | | | | Ribc1 |
| | | | | Ribc2 |
| | | | | Ric8B |
| | | | | Rif1 |
| | | | | Rigi |
| | | | | Rilp |
| | | | | Rilpl1 |
| | | | | Rilpl2 |
| | | | | Rimbp3 |
| | | | | Rin1 |
| | | | | Rinl |
| | | | | Ripk2 |
| | | | | Ripor2 |
| | | | | Rita1 |
| | | | | Rlbp1 |
| | | | | Rmdn1 |
| | | | | Rmdn2 |
| | | | | Rmdn3 |
| | | | | Rnd1 |
| | | | | Rnf128 |
| | | | | Rnf19A |
| | | | | Rnf4 |
| | | | | Rock1 |
| | | | | Rock2 |
| | | | | Ropn1L |
| | | | | Ror1 |
| | | | | Ror2 |

| | | | | |
|---|---|---|---|---|
| | | | | Rp1 |
| | | | | Rp1L1 |
| | | | | Rp2 |
| | | | | Rpap3 |
| | | | | Rpgr |
| | | | | Rpgrip1 |
| | | | | Rpgrip1L |
| | | | | Rpp25 |
| | | | | Rps3 |
| | | | | Rps6Ka1 |
| | | | | Rps6Ka2 |
| | | | | Rps7 |
| | | | | Rragd |
| | | | | Rrm1 |
| | | | | Rrp7A |
| | | | | Rsph1 |
| | | | | Rsph10B |
| | | | | Rsph14 |
| | | | | Rsph3A |
| | | | | Rsph3B |
| | | | | Rsph4A |
| | | | | Rsph6A |
| | | | | Rsph9 |
| | | | | Rtkn |
| | | | | Rtn2 |
| | | | | Rtraf |
| | | | | Rttn |
| | | | | Rusc1 |
| | | | | Ruvbl1 |
| | | | | Ruvbl2 |
| | | | | S100A8 |
| | | | | S100A9 |
| | | | | S100B |
| | | | | Saa1 |
| | | | | Saa2 |
| | | | | Sac3D1 |
| | | | | Samd14 |
| | | | | Sap30Bp |
| | | | | Sarm1 |
| | | | | Sass6 |
| | | | | Saxo1 |
| | | | | Saxo2 |
| | | | | Saxo4 |
| | | | | Sbds |
| | | | | Scin |
| | | | | Sclt1 |
| | | | | Scn8A |

| | | | | |
|---|---|---|---|---|
| | | | | Scnn1A |
| | | | | Sctr |
| | | | | Scyl1 |
| | | | | Sdcbp |
| | | | | Sdccag8 |
| | | | | Sele |
| | | | | Selenos |
| | | | | Sema4D |
| | | | | Septin1 |
| | | | | Septin10 |
| | | | | Septin11 |
| | | | | Septin12 |
| | | | | Septin14 |
| | | | | Septin2 |
| | | | | Septin3 |
| | | | | Septin4 |
| | | | | Septin5 |
| | | | | Septin6 |
| | | | | Septin7 |
| | | | | Septin8 |
| | | | | Septin9 |
| | | | | Serinc5 |
| | | | | Serp1 |
| | | | | Sestd1 |
| | | | | Sfi1 |
| | | | | Sfr1 |
| | | | | Sgca |
| | | | | Sgcb |
| | | | | Sgcd |
| | | | | Sgce |
| | | | | Sgcg |
| | | | | Sgcz |
| | | | | Sgf29 |
| | | | | Sgo1 |
| | | | | Sgpp1 |
| | | | | Sh2B2 |
| | | | | Sh3Gl1 |
| | | | | Sh3Kbp1 |
| | | | | Sh3Pxd2A |
| | | | | Sh3Pxd2B |
| | | | | Shank2 |
| | | | | Shcbp1 |
| | | | | Shcbp1L |
| | | | | Shld2 |
| | | | | Shmt2 |
| | | | | Shroom1 |
| | | | | Shroom2 |

| | | | | |
|---|---|---|---|---|
| | | | | Shroom3 |
| | | | | Shroom4 |
| | | | | Shtn1 |
| | | | | Sipa1L1 |
| | | | | Sipa1L3 |
| | | | | Sirt2 |
| | | | | Ska1 |
| | | | | Ska2 |
| | | | | Ska3 |
| | | | | Ski |
| | | | | Skp1 |
| | | | | Slain1 |
| | | | | Slain2 |
| | | | | Slc16A1 |
| | | | | Slc18A2 |
| | | | | Slc1A4 |
| | | | | Slc1A6 |
| | | | | Slc25A5 |
| | | | | Slc25A54 |
| | | | | Slc2A1 |
| | | | | Slc30A9 |
| | | | | Slc34A1 |
| | | | | Slc4A1 |
| | | | | Slc8A1 |
| | | | | Slc8A2 |
| | | | | Slc8A3 |
| | | | | Slf1 |
| | | | | Slmap |
| | | | | Smad3 |
| | | | | Smad4 |
| | | | | Smad6 |
| | | | | Smad7 |
| | | | | Smarca2 |
| | | | | Smc1A |
| | | | | Smc3 |
| | | | | Smc6 |
| | | | | Smg6 |
| | | | | Smo |
| | | | | Smtn |
| | | | | Snap25 |
| | | | | Snap29 |
| | | | | Snapin |
| | | | | Snca |
| | | | | Sncg |
| | | | | Snph |
| | | | | Snta1 |
| | | | | Sntb1 |

| | | | | |
|---|---|---|---|---|
| | | | | Sntb2 |
| | | | | Sntg1 |
| | | | | Sntg2 |
| | | | | Snx10 |
| | | | | Snx4 |
| | | | | Snx9 |
| | | | | Sorbs1 |
| | | | | Sorbs2 |
| | | | | Sorbs3 |
| | | | | Spaca9 |
| | | | | Spag16 |
| | | | | Spag17 |
| | | | | Spag4 |
| | | | | Spag5 |
| | | | | Spag6 |
| | | | | Spag6L |
| | | | | Spag8 |
| | | | | Spag9 |
| | | | | Spast |
| | | | | Spata13 |
| | | | | Spata4 |
| | | | | Spata7 |
| | | | | Spatc1 |
| | | | | Spatc1L |
| | | | | Spdl1 |
| | | | | Specc1 |
| | | | | Specc1L |
| | | | | Spef1 |
| | | | | Spef1L |
| | | | | Spef2 |
| | | | | Spice1 |
| | | | | Spin1 |
| | | | | Spire1 |
| | | | | Spire2 |
| | | | | Spmip10 |
| | | | | Spmip11 |
| | | | | Spmip4 |
| | | | | Spmip5 |
| | | | | Spmip6 |
| | | | | Spmip8 |
| | | | | Spmip9 |
| | | | | Spout1 |
| | | | | Sppl2B |
| | | | | Spry2 |
| | | | | Spta1 |
| | | | | Sptan1 |
| | | | | Sptb |

| | | | | |
|---|---|---|---|---|
| | | | | Sptbn1 |
| | | | | Sptbn2 |
| | | | | Sptbn4 |
| | | | | Sptbn5 |
| | | | | Sra1 |
| | | | | Src |
| | | | | Srcin1 |
| | | | | Srprb |
| | | | | Ss18 |
| | | | | Ssh1 |
| | | | | Ssh2 |
| | | | | Ssh3 |
| | | | | Ssna1 |
| | | | | Ssx2Ip |
| | | | | Stag1 |
| | | | | Stag2 |
| | | | | Stap1 |
| | | | | Stard9 |
| | | | | Stau2 |
| | | | | Steep1 |
| | | | | Stil |
| | | | | Stim1 |
| | | | | Sting1 |
| | | | | Stk11 |
| | | | | Stk17B |
| | | | | Stk3 |
| | | | | Stk33 |
| | | | | Stk36 |
| | | | | Stk38L |
| | | | | Stmn1 |
| | | | | Stn1 |
| | | | | Stom |
| | | | | Stoml2 |
| | | | | Stox1 |
| | | | | Stpg3 |
| | | | | Strbp |
| | | | | Stx1A |
| | | | | Stx1B |
| | | | | Svbp |
| | | | | Svil |
| | | | | Swap70 |
| | | | | Sybu |
| | | | | Syce1L |
| | | | | Sympk |
| | | | | Syn1 |
| | | | | Sync |
| | | | | Syne1 |

| | | | | |
|---|---|---|---|---|
| | | | | Syne2 |
| | | | | Synj1 |
| | | | | Synj2 |
| | | | | Synm |
| | | | | Synpo |
| | | | | Synpo2 |
| | | | | Synpo2L |
| | | | | Tacc1 |
| | | | | Tacc2 |
| | | | | Tacc3 |
| | | | | Tada2A |
| | | | | Tada3 |
| | | | | Taf1A |
| | | | | Taf1D |
| | | | | Taf5 |
| | | | | Tagln2 |
| | | | | Tagln3 |
| | | | | Taok1 |
| | | | | Taok2 |
| | | | | Tap1 |
| | | | | Tapt1 |
| | | | | Tax1Bp3 |
| | | | | Tbc1D21 |
| | | | | Tbc1D30 |
| | | | | Tbc1D31 |
| | | | | Tbc1D7 |
| | | | | Tbca |
| | | | | Tbcb |
| | | | | Tbccd1 |
| | | | | Tbcd |
| | | | | Tbce |
| | | | | Tbcel |
| | | | | Tbck |
| | | | | Tbl1X |
| | | | | Tbl1Xr1 |
| | | | | Tcea2 |
| | | | | Tchp |
| | | | | Tcp1 |
| | | | | Tcp11L1 |
| | | | | Tcp11L2 |
| | | | | Tcte1 |
| | | | | Tctn1 |
| | | | | Tctn2 |
| | | | | Tec |
| | | | | Tedc1 |
| | | | | Tedc2 |
| | | | | Tek |

| | | | | |
|---|---|---|---|---|
| | | | | Tekt1 |
| | | | | Tekt2 |
| | | | | Tekt3 |
| | | | | Tekt4 |
| | | | | Tekt5 |
| | | | | Tektip1 |
| | | | | Tektl1 |
| | | | | Tenm1 |
| | | | | Tent5C |
| | | | | Terf1 |
| | | | | Tesk1 |
| | | | | Tex35 |
| | | | | Tex9 |
| | | | | Tfdp2 |
| | | | | Tfpt |
| | | | | Tgfb1I1 |
| | | | | Tgif2 |
| | | | | Tiam1 |
| | | | | Tjp1 |
| | | | | Tjp2 |
| | | | | Tjp3 |
| | | | | Tlk2 |
| | | | | Tln1 |
| | | | | Tln2 |
| | | | | Tlnrd1 |
| | | | | Tm9Sf2 |
| | | | | Tmc2 |
| | | | | Tmem201 |
| | | | | Tmem214 |
| | | | | Tmem216 |
| | | | | Tmem232 |
| | | | | Tmem237 |
| | | | | Tmem63A |
| | | | | Tmem63B |
| | | | | Tmem67 |
| | | | | Tmem9 |
| | | | | Tmod1 |
| | | | | Tmod2 |
| | | | | Tmod3 |
| | | | | Tmod4 |
| | | | | Tmsb10 |
| | | | | Tmsb15B2 |
| | | | | Tmsb15L |
| | | | | Tmsb4X |
| | | | | Tmub1 |
| | | | | Tnik |
| | | | | Tnks |

| | | | | |
|---|---|---|---|---|
| | | | | Tnks1Bp1 |
| | | | | Tnks2 |
| | | | | Tnnc1 |
| | | | | Tnnc2 |
| | | | | Tnni1 |
| | | | | Tnni2 |
| | | | | Tnni3 |
| | | | | Tnnt1 |
| | | | | Tnnt2 |
| | | | | Tnnt3 |
| | | | | Tns1 |
| | | | | Tns3 |
| | | | | Tns4 |
| | | | | Togaram1 |
| | | | | Togaram2 |
| | | | | Top2A |
| | | | | Topbp1 |
| | | | | Topors |
| | | | | Tor1A |
| | | | | Tpgs1 |
| | | | | Tpgs2 |
| | | | | Tpm1 |
| | | | | Tpm2 |
| | | | | Tpm3 |
| | | | | Tpm3-Rs7 |
| | | | | Tpm4 |
| | | | | Tppp |
| | | | | Tppp2 |
| | | | | Tppp3 |
| | | | | Tpr |
| | | | | Tpt1 |
| | | | | Tpx2 |
| | | | | Tradd |
| | | | | Traf3Ip1 |
| | | | | Traf4 |
| | | | | Traf5 |
| | | | | Trappc14 |
| | | | | Trat1 |
| | | | | Trib2 |
| | | | | Trim32 |
| | | | | Trim36 |
| | | | | Trim46 |
| | | | | Trim54 |
| | | | | Trim67 |
| | | | | Trim69 |
| | | | | Trim75 |
| | | | | Trim9 |

| | | | | |
|---|---|---|---|---|
| | | | | Triobp |
| | | | | Trip10 |
| | | | | Trip4 |
| | | | | Trip6 |
| | | | | Trmt10A |
| | | | | Trp53 |
| | | | | Trpc4 |
| | | | | Trpv4 |
| | | | | Tsc1 |
| | | | | Tsen2 |
| | | | | Tsg101 |
| | | | | Tsga10 |
| | | | | Tsks |
| | | | | Tspan5 |
| | | | | Tssk2 |
| | | | | Tssk6 |
| | | | | Ttbk1 |
| | | | | Ttbk2 |
| | | | | Ttc12 |
| | | | | Ttc17 |
| | | | | Ttc21B |
| | | | | Ttc23L |
| | | | | Ttc28 |
| | | | | Ttc29 |
| | | | | Ttc39A |
| | | | | Ttc8 |
| | | | | Ttl |
| | | | | Ttll1 |
| | | | | Ttll10 |
| | | | | Ttll11 |
| | | | | Ttll12 |
| | | | | Ttll13 |
| | | | | Ttll2 |
| | | | | Ttll3 |
| | | | | Ttll4 |
| | | | | Ttll5 |
| | | | | Ttll6 |
| | | | | Ttll7 |
| | | | | Ttll8 |
| | | | | Ttll9 |
| | | | | Ttn |
| | | | | Tuba1A |
| | | | | Tuba1B |
| | | | | Tuba1C |
| | | | | Tuba3A |
| | | | | Tuba3B |
| | | | | Tuba4A |

| | | | | |
|---|---|---|---|---|
| | | | | Tuba8 |
| | | | | Tubal3 |
| | | | | Tubb1 |
| | | | | Tubb2A |
| | | | | Tubb2B |
| | | | | Tubb3 |
| | | | | Tubb4A |
| | | | | Tubb4B |
| | | | | Tubb5 |
| | | | | Tubb6 |
| | | | | Tubd1 |
| | | | | Tube1 |
| | | | | Tubg1 |
| | | | | Tubg2 |
| | | | | Tubgcp2 |
| | | | | Tubgcp3 |
| | | | | Tubgcp4 |
| | | | | Tubgcp5 |
| | | | | Tubgcp6 |
| | | | | Tulp3 |
| | | | | Twf1 |
| | | | | Twf2 |
| | | | | Txndc2 |
| | | | | Txndc9 |
| | | | | Tyk2 |
| | | | | Uaca |
| | | | | Ubn1 |
| | | | | Ubr4 |
| | | | | Ubxn11 |
| | | | | Ubxn2B |
| | | | | Ubxn6 |
| | | | | Ucma |
| | | | | Umod |
| | | | | Unc119 |
| | | | | Upf3B |
| | | | | Upp2 |
| | | | | Ush1C |
| | | | | Ush1G |
| | | | | Ush2A |
| | | | | Usp10 |
| | | | | Usp2 |
| | | | | Usp20 |
| | | | | Usp26 |
| | | | | Usp33 |
| | | | | Usp50 |
| | | | | Usp9X |
| | | | | Utrn |

| | | | | |
|---|---|---|---|---|
| | | | | Uvrag |
| | | | | Uxt |
| | | | | Vangl2 |
| | | | | Vapa |
| | | | | Vash2 |
| | | | | Vasp |
| | | | | Vcam1 |
| | | | | Vcl |
| | | | | Vcp |
| | | | | Vil1 |
| | | | | Vill |
| | | | | Vim |
| | | | | Vmac |
| | | | | Vps11 |
| | | | | Vps16 |
| | | | | Vps18 |
| | | | | Vps37A |
| | | | | Vps41 |
| | | | | Vps4A |
| | | | | Vps4B |
| | | | | Wapl |
| | | | | Was |
| | | | | Wasf1 |
| | | | | Wasf2 |
| | | | | Wasf3 |
| | | | | Washc1 |
| | | | | Wasl |
| | | | | Wbp2Nl |
| | | | | Wdpcp |
| | | | | Wdr1 |
| | | | | Wdr11 |
| | | | | Wdr13 |
| | | | | Wdr19 |
| | | | | Wdr35 |
| | | | | Wdr44 |
| | | | | Wdr47 |
| | | | | Wdr5 |
| | | | | Wdr62 |
| | | | | Wdr73 |
| | | | | Wdr90 |
| | | | | Whamm |
| | | | | Whrn |
| | | | | Wipf1 |
| | | | | Wipf2 |
| | | | | Wipi1 |
| | | | | Wnk1 |
| | | | | Wrap73 |

| | | | | |
|---|---|---|---|---|
| | | | | Wrn |
| | | | | Xirp1 |
| | | | | Xirp2 |
| | | | | Xrcc2 |
| | | | | Ybx3 |
| | | | | Yeats2 |
| | | | | Yes1 |
| | | | | Ypel5 |
| | | | | Ythdf2 |
| | | | | Ywhae |
| | | | | Zbed6 |
| | | | | Zbtb49 |
| | | | | Zc3H12A |
| | | | | Zfp12 |
| | | | | Zfp131 |
| | | | | Zfp185 |
| | | | | Zfp207 |
| | | | | Zfp322A |
| | | | | Zfp330 |
| | | | | Zfp365 |
| | | | | Zfp397 |
| | | | | Zfp804A |
| | | | | Zfyve19 |
| | | | | Zfyve26 |
| | | | | Zmynd10 |
| | | | | Zonab |
| | | | | Zw10 |
| | | | | Zyx |
| | | | | Zzz3 |
